# Transformer neural network for protein-specific de novo drug generation as a machine translation problem

**DOI:** 10.1101/863415

**Authors:** Daria Grechishnikova

## Abstract

Drug discovery for a protein target is a very laborious, long and costly process. Machine learning approaches and, in particular, deep generative networks can substantially reduce development time and costs. However, the majority of methods imply prior knowledge of protein binders, their physicochemical characteristics or the three-dimensional structure of the protein. The method proposed in this work generates novel molecules with predicted ability to bind a target protein by relying on its amino acid sequence only. We consider target-specific de novo drug design as a translational problem between the amino acid “language” and SMILES (Simplified Molecular Input Line Entry System) representation of the molecule. To tackle this problem, we apply Transformer neural network architecture, a state-of-the-art approach in sequence transduction tasks. Transformer is based on a self-attention technique, which allows the capture of long-range dependencies between items in sequence. The model generates realistic diverse compounds with structural novelty. The computed physicochemical properties and common metrics used in drug discovery fall within the plausible drug-like range of values.

## Introduction

Drug development is a multistage process that requires many resources. Bringing a drug to market may take up to 20 years [1]. The total cost may vary from US$0.5 billion to US$2.6 billion [2]. The estimated amount of drug-like molecule space is 10^60^, while the number of synthesized compounds is on the order of 10^8^ [3]. Therefore, the search for a promising molecule that may bind to a target protein is a challenging task for chemists. A high-throughput screening technique allows testing of millions of molecules in vitro to determine compounds that may act on the protein of interest [4]. However, this method is expensive and time-consuming. Virtual screening is used to search libraries of billions of molecules in silico [5]. This method requires information about compounds that are active against the protein or knowledge of the protein three-dimensional structure and operates on already known molecules, which span only a small part of the synthetically accessible molecule space. In de novo drug design, one has to create a molecule that is active toward the desired biological target from scratch. Existing computational methods often generate molecules that are hard to synthesize or restrict accessible chemical space via coded rules [6]. Despite all efforts, targeted generation of molecules remains a challenging task. Recently, machine learning methods were proposed to tackle this problem [7].

Most of the deep learning models for molecule generation are based on recurrent neural network (RNN). RNN is commonly used for modeling sequence data. The main feature of RNN allowing it to work with sequential data is the ability to make use of information from preceding steps. RNN can reveal links between distant elements of a sequence [8]. Unfortunately, RNNs suffer from the problem of vanishing gradients, which significantly limits their ability to work with long sequences. Long short-term memory and gated recurrent units partially solve this issue [9]. Recently, several works introduced recurrent neural networks based on the long short-term memory for de novo molecule generation [10–12]. They use Simplified Molecular-Input Line-Entry (SMILES) strings as input. Fine-tuning on a smaller dataset with compounds known to be active against biological targets force the models to generate focused molecule libraries with the desired activity toward the same target. Several research groups applied a reinforcement learning approach to bias the generator to produce molecules with desired properties [13–20]. In the reinforcement learning paradigm, the agent (generator in de novo drug generation problem) takes some action (choosing the next character during new SMILES string generation) to maximize reward (function computed after SMILES string completion). Olivecrona et al. fine-tuned the RNNs to generate compounds binding Dopamine Receptor Type 2 (DRD2). To predict molecule activity, they built a Support Vector Machine (SVM) classifier with a Gaussian kernel and trained it on the DRD2 activity dataset [13]. The output of this model was used to formulate the reward function. Popova et al. [14] suggested training separately two neural networks – generative and predictive – and then using them jointly to generate novel libraries of compounds with the desired properties, e.g., targeted toward Janus kinase 2. Several research groups applied the generative adversarial network concept to design compounds with optimized properties, but they did not consider activity against any biological target [17,18].

Another fundamental approach to de novo compound generation is based on autoencoder architecture [21–28]. Autoencoder consists of encoder and decoder networks [8]. The former one converts the input data into a latent representation (vector of fixed dimension), and the second one reconstructs the initial object from the latent code. The hidden layer forming the latent representation vector is an informational bottleneck, which induces the model to capture the most important features of the input object [8]. Variational and adversarial autoencoders are two types of autoencoders that are widely used to generate molecules. In variational autoencoders, a prior distribution, usually normal, is imposed on latent space to make it smooth and suitable for sampling [29]. In adversarial autoencoders, the discriminator neural network is introduced into architecture to force the distribution of latent codes to follow arbitrary prior distribution [30]. Gómez-Bombarelli et al. [21] suggested a variational autoencoder extended by attaching a multilayer perceptron to the latent layer for property prediction. Joint training of this enlarged model forces the latent space to organize by property values. On top of this model, authors trained the Gaussian process to predict target compound properties using the latent space representation as input. In a recent publication [22], the authors compared several autoencoder architectures including variational and adversarial ones. The adversarial autoencoder provides the highest fraction of valid SMILES strings. The authors trained the SVM classifier to predict activity against DRD2. They used this probability as the objective function and maximized it during the latent space Bayesian optimization. Additionally, an autoencoder can be used for a conditional generation [31–33]. In these studies, properties were directly imposed on latent space during the training. Polykovskiy et al. introduced a conditional adversarial autoencoder to design compounds with specified properties [33]. After training on a set of Janus kinase 2 (JAK2) and Janus kinase 3 (JAK3) inhibitors and conditioning on the selective activity against JAK2, the model generated a compound that turned out to be active toward JAK2 during in vitro tests. Recently, Zhavoronkov et al. developed a discoidin domain receptor 1 (DDR1) inhibitor in 21 days using a variational autoencoder fine-tuned with the reinforcement learning approach [25]. One molecule successfully passed experiments in mice.

However, all these methods imply prior knowledge of protein binders and their physicochemical characteristics. Structure-based drug design approaches require the three-dimensional structure of the protein. In this work, we introduce an approach to targeted drug generation that uses only the protein amino acid sequence as input. We consider the target-specific de novo drug generation problem as a translation from the amino acid “language” to SMILES representation of the molecule. Recently, Transformer-based models demonstrated state-of-the-art results on neural machine translation tasks [34,35]. We adopt Transformer to generate molecules. The network takes amino acid sequence as input and generates molecules with predicted ability to bind the protein target. The model outputs valid structures with plausible values of computed physicochemical characteristics, a drug-likeness metric, and a synthetic accessibility score.

The main contributions of our work are as follows:

1. We formulate the targeted drug generation problem as a translational task and apply the Transformer architecture. This application allows molecules generation based on protein amino acid sequence only.
2. Our approach requires neither prior knowledge of protein binders nor preparation of libraries of ligands active against the target.
3. The proposed model is based on a self-attention technique that allows better capture of long-range dependencies than recurrent neural networks.

## Methods

### Data

We retrieved data from BindingDB [36]. BindingDB contains a measured binding affinity of interactions between proteins and drug-like molecules. The full database version was downloaded. The raw dataset contained over 1.5 million data records. We selected records from the raw dataset using the following criteria:

1. The field “Target Source Organism According to Curator or DataSource” equals “Homo sapiens” or “Rattus norvegicus” or “Mus musculus” or “Bos taurus”.
2. The record has an IC50 value less than 100 nm; if the IC50 is missing, then Kd is less than 100 nm; if both are missing, then EC50 is less than 100 nm.
3. The record has a chemical identifier (PubChem CID).
4. The record has SMILES representation.
5. The molecular weight is less than 1000 Da.
6. The record has a protein identifier (Uniprot ID).
7. Protein amino acid sequence length is greater than 80 and lower than 2050.

This yielded a results dataset containing 238147 records. There were 1613 unique amino acid sequences and 154924 unique ligand SMILES strings. All SMILES strings used in this work were canonicalized using RDKit. We created five different splits into test and training parts.

In each split we required the similarity between proteins in the test and the ones in the training dataset to be less than 20%. This additional condition leaded to the removal of some proteins from the dataset, so final sizes of the training and test datasets can be found in Table 1.

**Table 1.**
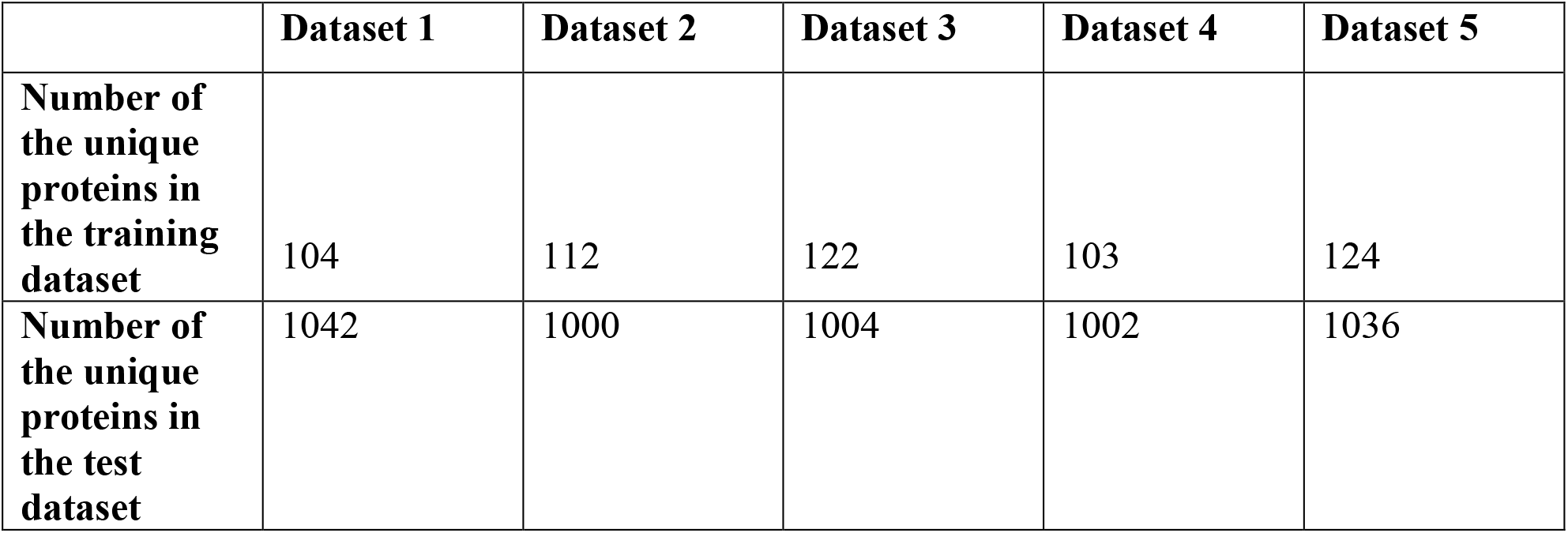
The number of unique proteins in the training and test datasets for each split.

In each split we used a proportion common for machine learning tasks – roughly 90% for train part and 10% for test part. To compute similarities, we used the Needleman-Wunsch global alignment algorithm from the EMBOSS package [37]. The distribution of pairwise sequence similarities for the first split is shown in Figure 1.

**Figure 1.**
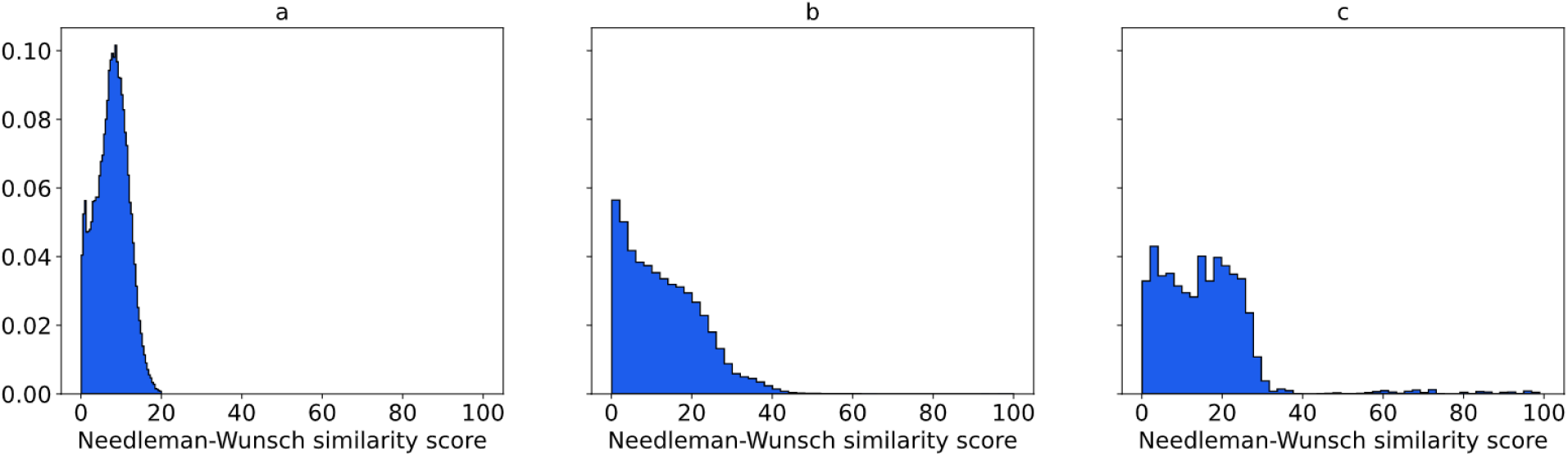
Distributions of sequence similarities between proteins used for model training and validation. (a) Sequence similarities between proteins in the test dataset and the ones in the training dataset, (b) sequence similarities of proteins within the test dataset, (c) sequence similarities of proteins within the training dataset.

The distributions for other splits are analogous. Figure 1a shows that protein sequences from the test dataset share less than 20% similarity with those in the training dataset. At the same time, the protein sequences within the test and training sets are also diverse enough to train and test the model. The majority of sequences share less than 40% similarity (Figures 1b and 1c).

### Data representation

We considered each symbol in an amino acid sequence or in a SMILES string as a token. The vocabulary was determined by the dataset and contained 71 symbols. Each token was converted into a vector using trainable embedding in the first layer of the encoder.

### Model

We adopted the Transformer model for targeted drug generation using the original implementation described in [35]. Transformer has an encoder-decoder structure. The encoder maps a protein amino acid sequence (*a_1_*,…,*a_n_*) to a sequence of continuous representations *z*=(*z_1_*, …, *z_n_*). Then, the decoder takes *z* as input and generates a SMILES string in autoregressive manner. At every step of generation, the decoder may attend to any elements of *z* due to the attention mechanism. The latent code *z* may be considered as a “protein context” used by the decoder to generate a molecule structure. The model yields a probability distribution over each element in the vocabulary for each position in the output sequence. Transformer is based on an attentional mechanism only. It lacks any kind of convolutional or recurrent neural network components. Transformer uses self-attention to compute the representations of input and output sequences. Self-attention refers to different components of a single sequence in relation to other components to compute sequence representation. Each layer of the encoder is composed of a multihead self-attention sublayer and feed-forward sublayer. In addition to these, each layer of the decoder has a multihead attention layer attending the encoder output.

The self-attention mechanism successfully copes with long-range dependencies while being faster than recurrent layers. The attention layer at first calculates three vectors from each “word” of a “sentence” – key, query and value. To process all words in a sentence simultaneously, key vectors are packed together into matrix *K*, and queries and values produce matrices *Q* and *V* correspondingly. In our task definition, “words” are amino acid residuals or characters in SMILES strings. The attention is computed as follows:

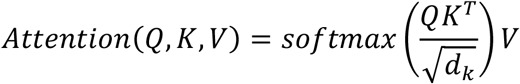
where *d_k_* is a scaling factor.

The multihead attention mechanism produces *h* different representations of *Q, K, V* values and computes an attention function for each representation:

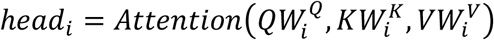

The outputs are concatenated and projected one more time, yielding final values:

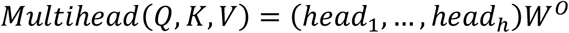
where 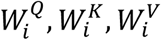 are matrices of learned weights.

Since the model lacks any recurrent component, it has no information about the order of tokens in a sequence. To address this lack of information, the model adds position-dependent signals to the input embedding. There are many possible choices for signal functions. In Transformer, sine and cosine functions are used:

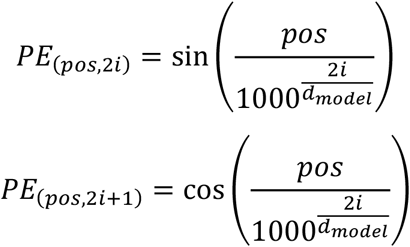

where *pos* is the position, *i* is the dimension and *d_model_* is the size of embedding.

We use beam search to decode SMILES strings. While constructing a sequence, the beam search evaluates all possible next steps and keeps the top n candidates with the highest probability, where n is the user-specified parameter referred to as beam size. If beam size is equal to one, the beam search becomes the greedy search. If beam size is greater than one, the output sequences differ only slightly from each other. It might be beneficial if a generated molecule is good enough, and small structure optimizations are needed. However, in the process of target-specific de novo drug generation, it would be better to have more diverse variants per certain protein. There are several ways to achieve potential improvement. We discuss them in the “Results and Discussion” section. For each protein, we ran beam search with beam sizes of 4 and 10. In the first case, we left only one SMILES string with the highest probability (one per one mode). In the second case, we left all ten generated molecules for subsequent analysis (ten per one mode).

All work was performed in Google Colaboratory. We used the open-source tensor2tensor library for building, training and testing the model [38]. We experimented with different numbers of attentional heads, layers, and their sizes. The optimal proportion between the amount of valid and unique SMILES strings gives the model containing four layers of size 128 and four attention heads. We used the Adam optimizer and learning rate decay scheme proposed in [35], and the batch size was set to 4096 tokens. We trained the model for 600K epochs using one GPU.

To test the model, we performed Monte-Carlo cross validation. We split all proteins so that test dataset contains only sequences sharing less than 20% similarity with those in the training dataset. Then, we trained the model and tested it on selected proteins. This procedure was repeated five times.

### Model evaluation

We used RDKit [39] to check chemical validity, calculate properties, compute similarity scores and produce SMILES canonical representation of molecule structures. Molecules known to be active against given target proteins and the generated ones were docked in binding sites using SMINA with default settings [40]. Protein PDB structures were downloaded from the Protein Data Bank [41]. We followed the standard procedure to prepare protein for docking, heteroatoms were removed, and hydrogens were added via PyMol [42]. We utilized OpenBabel [43] to generate three-dimensional conformers.

## Results and Discussion

### Chemical feasibility

This section demonstrates the effectiveness of the proposed approach for the generation of valid realistic molecules. We created five different divisions of the initial dataset to train and test parts.

For each division, we performed training of the model followed by validation on the corresponding test set. At first, we ran the model in one per one mode (see Methods). For each protein in test datasets, the model generated a molecule. We checked the chemical validity of molecules with RDKit software, analyzed uniqueness and searched the ZINC15 database for generated compounds. All characteristics were averaged across five test sets. Approximately 90% of generated molecules were valid, and 92% were unique (Table 2). Approximately 30% of compounds were found in the ZINC15 database. The entire list of generated SMILES strings can be found as Supplementary Table S1. We also provide the figures with their graph representations (see Supplementary Figure S2).

**Table 2.** Percentages of valid, unique and ZINC15 database-matched SMILES strings generated by the model in one per one mode.

|  | Dataset 1 | Dataset 2 | Dataset 3 | Dataset 4 | Dataset 5 | Average |
| --- | --- | --- | --- | --- | --- | --- |
| **Total number of generated SMILES strings (one per target protein)** | 104 | 112 | 122 | 103 | 124 | 113 |
| **Valid (%)** | 92.3 | 90.2 | 87.7 | 92.2 | 88.7 | 90.2 |
| **Unique (%)** | 93.3 | 90.2 | 90.9 | 94.2 | 92.7 | 92.3 |
| **Match with ZINC15 database (%)** | 33.6 | 22.3 | 27.0 | 31.0 | 39.5 | 30.6 |

In the case of generating one ligand per protein, the outputted compound might be considered as a valid starting point for a subsequent modification during the drug discovery process. Nevertheless, it would be useful to obtain more drug candidates for the given target protein. To achieve this aim, we expanded beam size to ten, allowing the model to output the ten most likely variants of the compound for inputted protein (ten per one mode). In this mode, the model generated almost 83% valid SMILES strings and 82% unique SMILES strings on average across five datasets (Table 3). Over 17% of novel compounds matched the ZINC15 database.

The number of valid and unique SMILES strings is lower in ten per one mode. We assume that this is caused by the problem of performance degradation in the beam search. A recently proposed method may possibly increase the performance [44]. However, this improvement is outside the scope of our work.

**Table 3.** Percentages of valid, unique and ZINC15 database-matched SMILES strings generated by the model in ten per one mode.

|  | Dataset 1 | Dataset 2 | Dataset 3 | Dataset 4 | Dataset 5 | Average |
| --- | --- | --- | --- | --- | --- | --- |
| **Total number of generated SMILES strings (ten per target protein)** | 1040 | 1120 | 1030 | 1220 | 1240 | 1130 |
| **Valid** | 80.8 | 80.5 | 79.6 | 86.2 | 86.1 | 82.6 |
| **Unique** | 88.4 | 78.4 | 71.7 | 85.0 | 85.0 | 81.7 |
| **Match with ZINC15 database (%)** | 17.9 | 16.3 | 10.7 | 19.5 | 21.2 | 17.1 |

### Testing the binding affinity between generated compounds and target proteins

In this section, our goal is to assess whether generated molecules could bind the target protein. At first, we randomly shuffled the test dataset (split 1), which contains 104 proteins. All proteins share less than 20% sequence similarity with those in the training dataset. Then, we consequently checked each protein and selected the ones satisfying the criteria:

- Protein is from human
- More than 100 known binders were selected from BindingDB using criteria from the Data section
- Protein contains one druggable cavity

The last criterion was related to technical limitations. Molecular docking is a very resource-consuming procedure. We were able to analyze several PDB structures only for a pair of proteins with one druggable cavity. Docking of many ligands into several structures with many binding pockets requires a lot more computational time than we possess. The vast majority of proteins in the test dataset have many binding pockets. To satisfy the last criterion, we had to choose proteins with one well-known binding pocket, which is mostly used as a target site for small molecule drugs. Therefore, we selected two proteins from the receptor tyrosine kinases family. They contain an extracellular ligand-binding region, transmembrane domain, and an intracellular region with a tyrosine kinase domain [45]. Binding of a specific ligand to an extracellular region induces phosphorylation process, leading to structural transformation within the kinase domain. This results in activation of a corresponding signal pathway. The vast majority of reported kinase inhibitors binds to the catalytic domain essential for kinase activity [45]. The first selected protein is the Insulin-like growth factor 1 receptor (IGF-1R). IGF-1R is a transmembrane receptor with tyrosine kinase activity. It can bind three ligands: insulin and the two insulin-like growth factors (IGF-I, IGF-II) [46]. It is involved in the development of many tissues and plays a key role in the regulation of overall growth and metabolism. IGF-1R is known to contribute to the pathophysiology of cancer via involvement in cell transformation, proliferation and metastatic events [47]. This involvement makes IGF-1R a valuable target for drug development. One of the strategies aimed at blocking IGF-1R activity is to use small molecules as IGF-1R tyrosine kinase inhibitors [46].

The second protein is Vascular endothelial growth factor receptor 2 (VEGFR2). VEGFR2 is a cell-surface receptor with tyrosine kinase activity [48]. Three growth factors bind to VEGFR2: VEGFA, VEGFC, and VEGFD. Ligand binding initiates a phosphorylation process leading to an enhancement of endothelial cell proliferation and migration. VEGFR2 is expressed on vascular endothelial cells and lymphatic endothelial cells. It plays a critical role in physiologic and pathologic angiogenesis, vascular development and permeability and embryonic hematopoiesis. It is involved in the development of many diseases including cancer, arthritis, and diabetes [48].

For each protein, we composed four sets of ligands – known binders, compounds randomly chosen from BindingDB, molecules generated for a selected protein and molecules generated for other targets in the test dataset. We collected 1148 known binders from BindingDB for IGF-1R and 3782 compounds for VEGFR2 using the criteria mentioned in the Data section. We could not dock all of them to proteins due to technical limitations. Therefore, we randomly selected 100 ligands for docking experiments. The second set contains 100 compounds randomly selected from BindingDB. The third set includes 11 generated molecules (one in one per one mode and ten in ten per one mode) for each protein. To test whether generated compounds can bind to the “wrong” target (cross-docking), we also formed a set of 100 molecules generated for other proteins in the test dataset. The binding scores between ligands and target protein active sites were computed using SMINA. For each protein, we downloaded protein structures bound to ligands from the PDB database. We obtained 11 PDB files (2OJ9, 3D94, 3I81, 3NW5, 3NW6, 3NW7, 3O23, 4D2R, 5FXQ, 5FXR, 5FXS) for IGF-1R and 20 (1Y6A, 1Y6B, 2P2H, 2QU5, 2RL5, 2XIR, 3BE2, 3EWH, 3U6J, 3VHE, 3VHK, 3VID, 3VNT, 3VO3, 4AG8, 4AGC, 4AGD, 4ASD, 4ASE, 6GQP) for VEGFR2. We examined all structures to ensure that they contain binding pockets. Then, we aligned them via PyMol. All ligands were extracted and combined into separate PDB files. We used them to define search space for SMINA. The docking requires many computational resources; therefore, we were not able to analyze all PDB structures for each protein. Thus, we selected structures that represent discrimination ability between known binders and randomly selected compounds. We utilized the ROC curve and corresponding area under curve (AUC) of the scores calculated by SMINA to evaluate whether the docking tool could discriminate between them. We checked six structures for IGF-1R (2OJ9, 3O23, 3I81, 4D2R, 5FXQ, 5FXR) and four structures for VEGFR2 (2P2H, 3BE2, 4ASE, 6GQP). Structures with PDB codes 3I81, 5FXQ, 5FXR, and 6GQP failed in the discrimination of active and randomly selected compounds (AUC<0.6). We removed them from the subsequent analysis.

We further assessed whether the docking tool could discriminate between binders and molecules generated for other targets, between generated and randomly selected compounds and between generated and known binders. The structures with PDB codes 3O23 and 3BE2 demonstrate the best discriminative ability between known binders and randomly selected compounds. Figure 2 shows ROC curves and corresponding AUC values for several combinations of molecule sets for both PDB structures.

**Figure 2.**
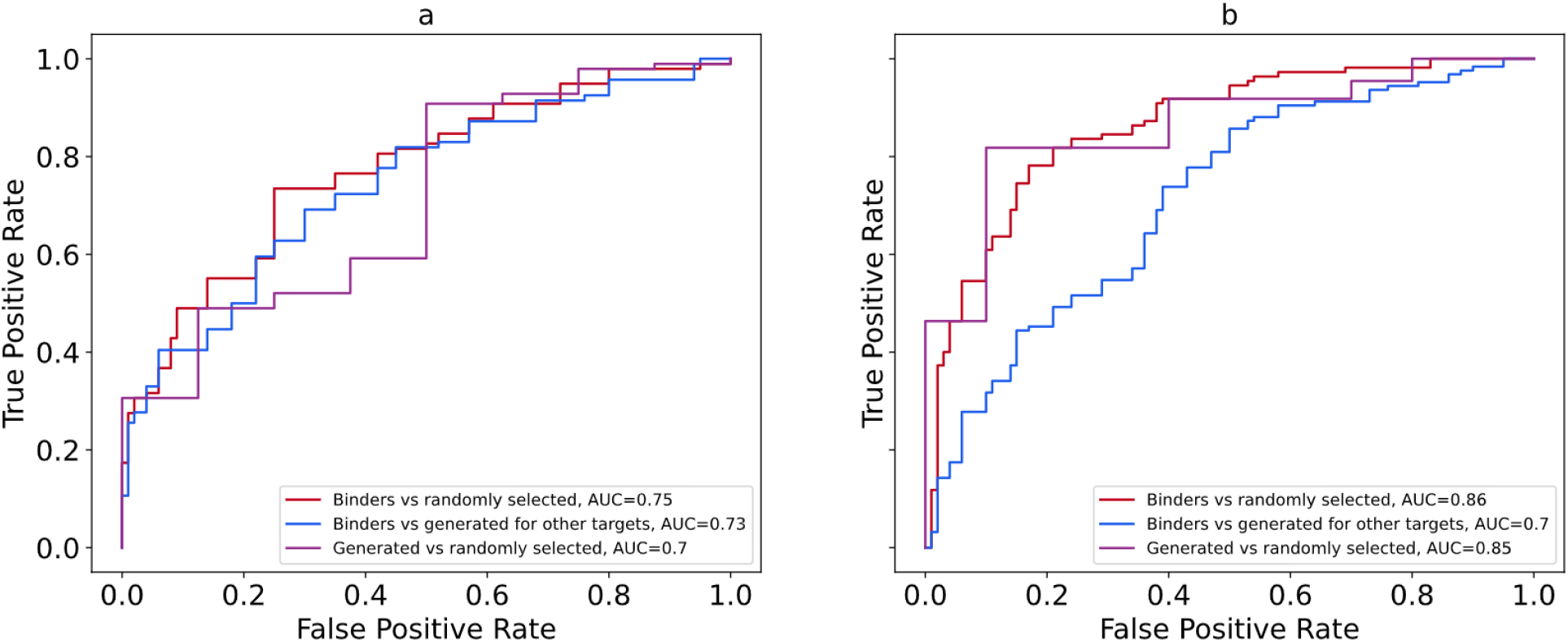
ROC curves and corresponding AUC for the following structures: (a) structure of IGF-1R with PDB code 3O23, (b) structure of VEGFR2 with PDB code 3BE2.

All AUC values are considerably higher than random baseline (0.5). These results indicate that the tool more likely classifies compounds generated for the IGF-1R and VEGFR2 as binders. At the same time, it less likely classifies compounds generated for other targets as binders. Interestingly, for the VEGFR2 the AUC value computed for the group of generated compounds versus group of randomly selected compounds is very close to the one computed for the set of known binders versus randomly selected molecules. Four other structures (4D2R, 2OJ9 for the IGF-1R target and 4ASE, 2P2H for the VEGFR2 target) present slightly lower discrimination ability; however, AUC values are very close to those computed for 3O23 and 3BE2 respectively. The ROC curves and their corresponding AUC values can be found as Supplementary Figure S3. We also build ROC curves to evaluate whether the tool could discriminate between compounds generated for analyzed structures and known binders (Figure 3 and Supplementary Figure S4). The AUC values are close to 0.5 in all cases meaning that the tool is unable to distinguish between these groups of molecules. It is well known that AUC values are directly connected to the U statistics of the Mann-Whitney test:

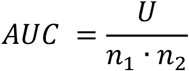

**Figure 3.**
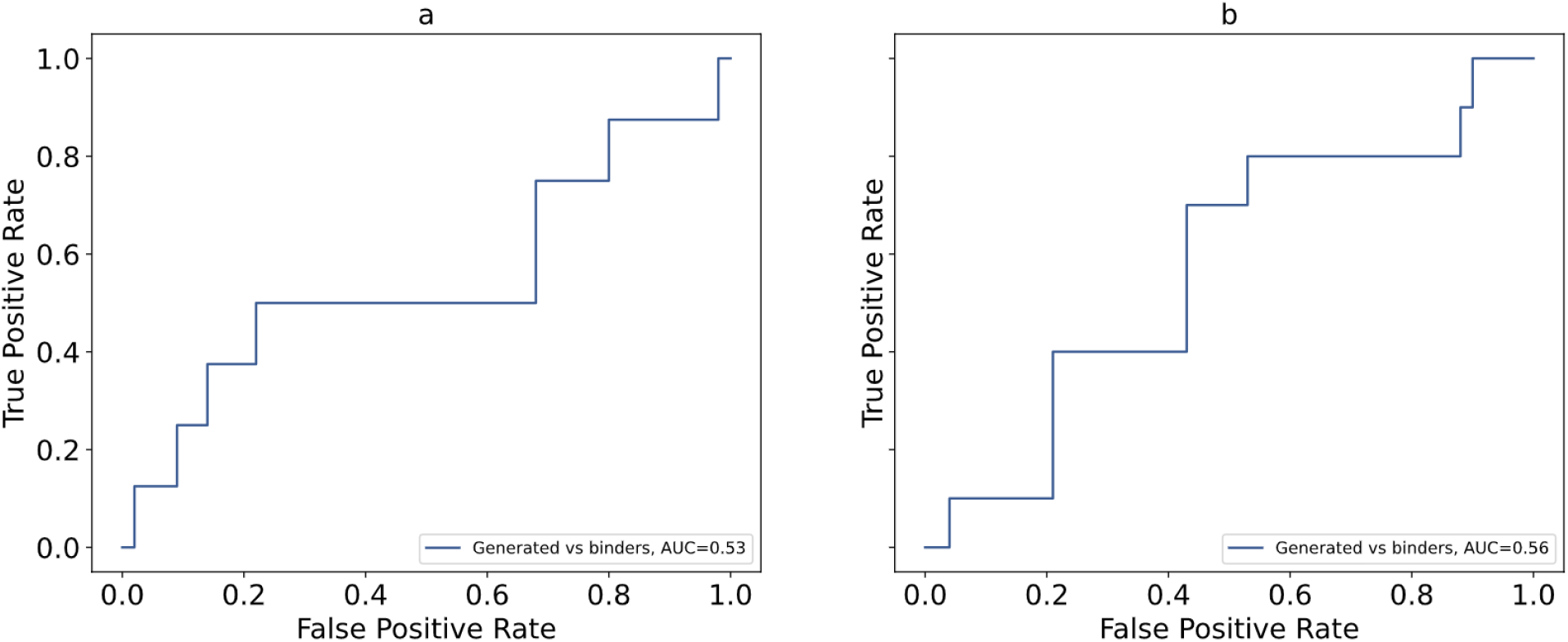
ROC comparison of known binders versus molecules generated for IGF-1R and VEGFR2 and corresponding AUC for the following structures: (a) structure of IGF-1R with PDB code 3O23, (b) structure of VEGFR2 with PDB code 3BE2.

Where *n*_1_ and *n*_2_ are sizes of the classes used. We assess significance of the difference between classes in each pair by p-value of the U statistics and present those values in the Table 4. It is clearly seen, that known binders are significantly different from the randomly selected compounds and compounds generated for other targets. Significance of the difference between randomly selected compounds and the ones generated for this target is smaller, but still valuable. In opposite, difference between known binders and compounds generated for this target is not significant, meaning that the model could not distinguish them.

**Table 4.** P-values of the Mann-Whitney test for all molecule sets used in the analysis.

| PDB code | P-values |  |  |  |
| --- | --- | --- | --- | --- |
|  | Known binders vs randomly selected compounds | Known binders vs compounds generated for other targets | Compounds generated for the analyzed protein vs randomly selected ones | Compounds generated for the analyzed protein vs known binders |
| 3O23 | $3.4 \cdot 10^{-10}$ | $2.2 \cdot 10^{-8}$ | $3.0 \cdot 10^{-2}$ | 0.40 |
| 4D2R | $1.5 \cdot 10^{-8}$ | $1.4 \cdot 10^{-6}$ | $2.9 \cdot 10^{-2}$ | 0.33 |
| 2OJ9 | $5.5 \cdot 10^{-7}$ | $5.2 \cdot 10^{-5}$ | $4.5 \cdot 10^{-2}$ | 0.44 |
| 3BE2 | $2.5 \cdot 10^{-19}$ | $8.4 \cdot 10^{-8}$ | $1.1 \cdot 10^{-4}$ | 0.26 |
| 4ASE | $1.6 \cdot 10^{-10}$ | $2.3 \cdot 10^{-5}$ | $4.1 \cdot 10^{-3}$ | 0.31 |
| 2P2H | $2.2 \cdot 10^{-12}$ | $3.6 \cdot 10^{-4}$ | $3.1 \cdot 10^{-3}$ | 0.17 |

These results suggest that generated molecules for IGF-1R and VEGFR2 can be considered binders, while the molecules generated for other targets are more likely to be nonbinders for both proteins.

We also visualized complexes of both proteins and generated ligands using PyMol software. Figure 4 shows the docking poses of generated molecules with the lowest scores in the binding pocket of the corresponding targets.

**Figure 4.**
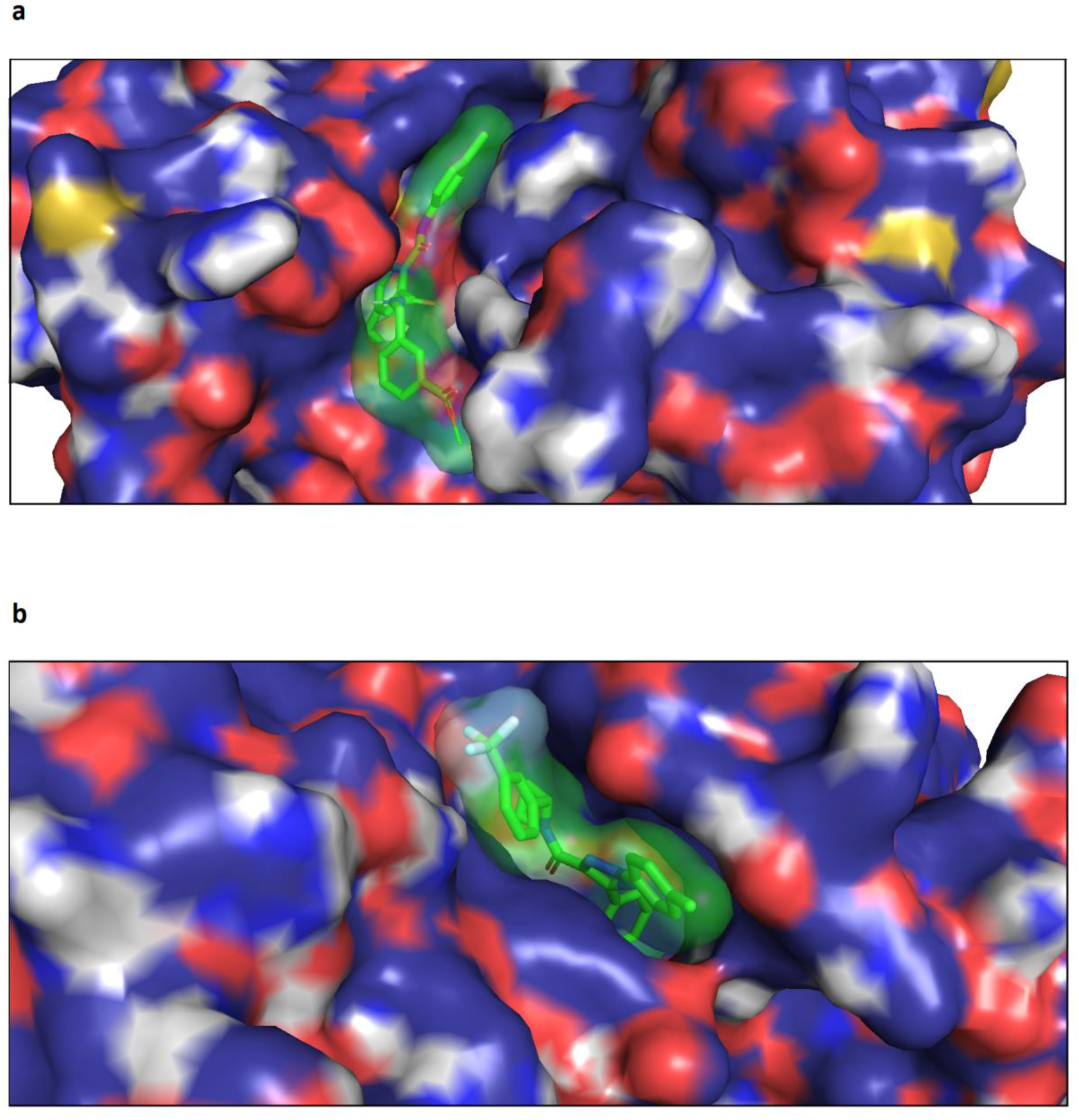
Positions of the generated molecules with the lowest scores in the binding sites of the following proteins: (a) Insulin-like growth factor 1 receptor, (b) Vascular endothelial growth factor receptor 2.

We realize that accurate estimation of the binding ability of generated molecules requires the analysis of many diverse proteins using in silico docking and/or in vitro assays. However, these are separate and quite complicated tasks, which we consider a direction for future work. We believe that the analysis described here is sufficient for the proof of concept.

### Physicochemical properties and metrics

It is not enough for the model to output chemically valid molecules active against a certain target. The model should also take care of parameters crucial for a molecule to be a potential drug. We computed several important metrics and physicochemical properties for generated compounds and compared them with corresponding characteristics of the molecules from the training dataset. The goal was to access the ability of the model to generate compounds satisfying typical drug-likeness metrics. According to the famous Lipinski’s rule of five, the water-octanol partition coefficient (logP) of a potential orally active drug should not exceed five. Molecules with molecular weight less than 500 show better permeability and solubility. The numbers of hydrogen donors, acceptors and rotatable bonds have to be no more than 5, 10 and 10, respectively [49,50]. Although Lipinski’s rule was developed for oral medications, it gives a good reference point for evaluating the properties of the generated molecules.

The Topological Polar Surface Area (TPSA) is another important characteristic of a drug candidate. Chemists assume that molecules having a topological polar surface area greater than 140Å^2^ are absorbed poorly [50]. To overcome the blood-brain barrier, a molecule should have a TPSA less than 90 Å^2^ [51]. Quantitative Estimate of Drug-likeness (QED) is based on the desirability functions for molecular properties and is widely used to select appropriate compounds during the early stages of drug discovery. In other words, QED is the measure of drug-likeness [52]. It ranges from zero to one, where zero indicates a totally unsuitable molecule, while one corresponds to molecules with favorable characteristics. The Synthetic Accessibility Score (SAS) is of great importance, as many computational approaches often yield molecules that tend to be difficult to synthesize (SAS > 6) [53]. Table 5 summarizes data about the compliance of the generated molecules with the rules mentioned above across five datasets used for Monte-Carlo cross-validation. For each constraint, the majority of generated compounds lie in acceptable drug-like molecule boundaries. Figure 5 shows the distributions of logP, the number of H-donors, H-acceptors, and rotatable bonds, QED, SAS, TPSA, molecular weight and length for the first test dataset.

**Figure 5.**
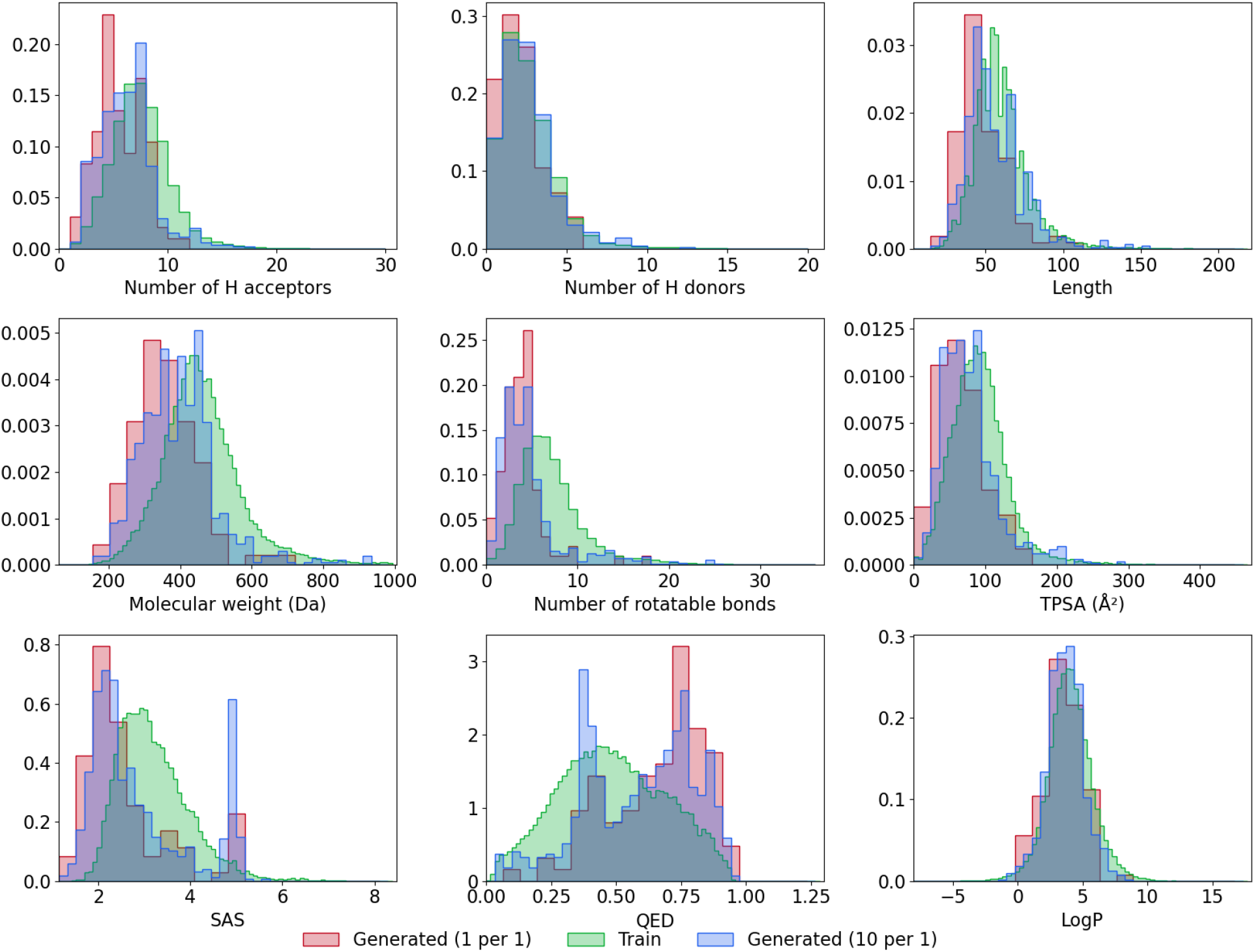
Distribution of properties for the generated molecules. Properties include: water-octanol partition coefficient (logP), the number of H-donors, the number of H-acceptors, the number of rotatable bonds, Quantitative Estimation of Drug-likeness (QED), the synthetic accessibility score (SAS), total polar surface area, molecular weight and length.

**Table 5.** Percentage of generated molecules falling within plausible drug-like molecule ranges of values.

| Property name | Constraints | Structures satisfying the constraints (%) |  |
| --- | --- | --- | --- |
|  |  | Generated molecules (one per one) ** | Generated molecules (ten per one) ** |
| logP | <5 | 84.4 | 85.6 |
| Molecular weight (Da) | <500 | 95.8 | 88.9 |
| Number of hydrogen donors | <5 | 95.8 | 91.9 |
| Number of hydrogen acceptors | <10 | 97.9 | 93.5 |
| Number of rotational bonds | <10 | 97.9 | 91.2 |
| Topological polar surface area (Å<sup>2</sup>) | <140 | 98.0 | 92.7 |
| Quantitative Estimate of Drug-likeness (QED) * |  | 0.66 ± 0.19 | 0.58 ± 0.21 |
| Synthetic accessibility score | <6 | 99.9 | 100.0 |

\* There is no common threshold for QED. QED varies in a range [0,1]. The higher a QED value is, the better. The columns show mean values and standard deviations.
\*\*averaged across five cross validational datasets

The distributions for the four remaining test datasets are almost identical. We analyzed computed characteristics of molecules from three datasets: structures generated in one per one mode, ten per one mode and the training set. For each parameter, the histograms display almost complete overlap between datasets. This overlap indicates that the model reproduces the property distribution of molecules in the training set very well.

The favorable values of these parameters do not necessarily indicate that the generated compound will become a drug. It can be checked only in an experiment. Nevertheless, we can conclude that generated molecules may be considered starting points for developing novel drugs with activity against given protein targets.

We assessed the structural diversity between generated molecules and molecules from the training dataset by calculating the Tanimoto similarity score implemented in RDKit. Figure 6 shows the distributions of the nearest neighbor Tanimoto coefficients over all pairs of these molecules.

**Figure 6.**
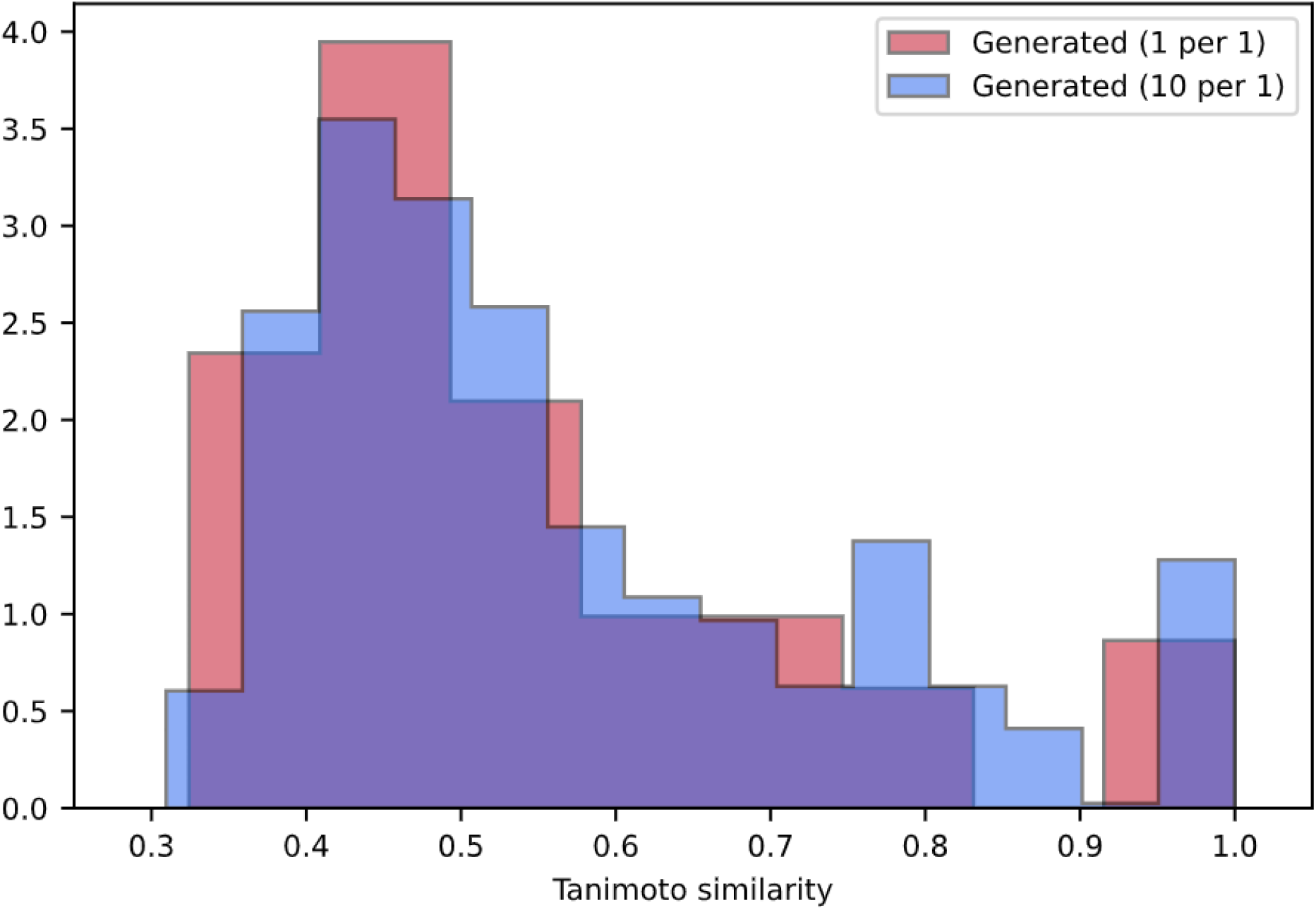
Tanimoto similarity of generated molecules to the nearest neighbor in the training dataset.

Only 8% of all generated structures have a Tanimoto score above the similarity threshold (Tanimoto score > 0.85) and can be considered similar to structures from the training dataset. The majority of generated molecules (51 %) has a Tanimoto score lower than 0.5, which suggests that this part of the generated compounds differ significantly from those in the training dataset. A high Tanimoto score usually indicates small differences in the molecule structure. However, even small differences in structure may lead to significant changes in functionality. Figure 7 demonstrates the distributions of the nearest neighbor Tanimoto similarities over all pairs of ligands in the training dataset.

**Figure 7.**
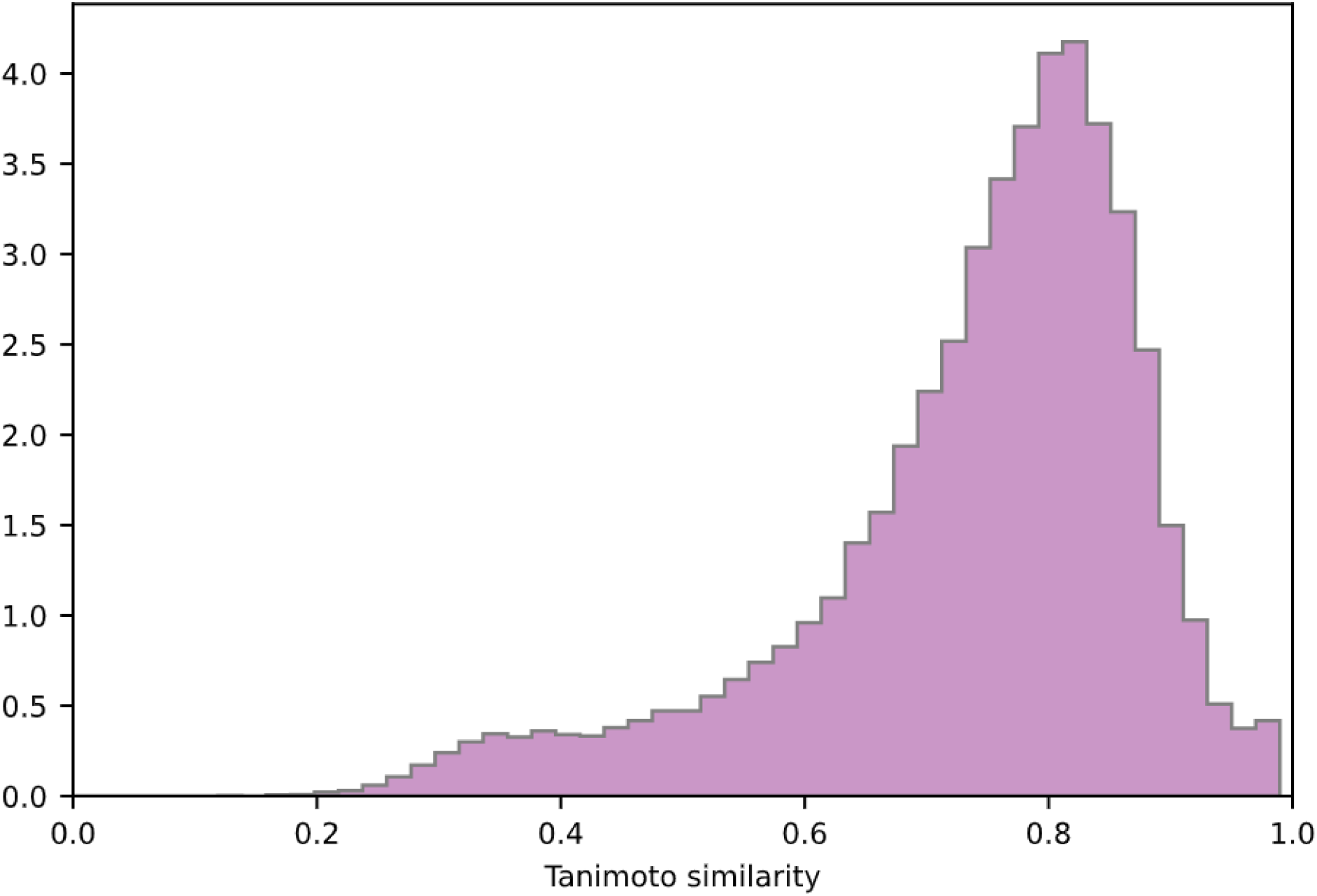
Tanimoto similarity of molecules which are nearest neighbors in the training dataset.

Mean and standard deviation values are shown in Table 6.

**Table 6.** Mean and standard deviation values of Tanimoto similarity distributions.

| Distribution | Mean | Standard deviation |
| --- | --- | --- |
| Tanimoto similarity of the generated molecules to the nearest neighbor in the training dataset (1 per 1 mode) | 0.54 | 0.17 |
| Tanimoto similarity of the generated molecules to the nearest neighbor in the training dataset (10 per 1 mode) | 0.57 | 0.18 |
| Tanimoto similarity of the molecules which are nearest neighbors in the training dataset | 0.74 | 0.14 |

The mean value of similarities between generated molecules and those in the training set is much lower than the mean value of similarities between compounds in the training dataset. Compared to the input dataset, the model achieves the generation of more diverse molecules, demonstrating the ability to create novel structures outside the training dataset.

### Transformer applicability to drug generation tasks

Deep learning methods usually need a library of molecules with known activity against a certain protein to generate ligand binding with the target. The specific library is used to fine-tune the model or to train a predictive network that assigns a reward to generator output in a reinforcement learning approach (e.g., [10,14,22]). In several research works, authors used a seed molecule to generate structures with the desired activity (e.g., [27,28]). In other words, these approaches demand some prior information about compounds that are active against a given target. The method proposed in this work does not imply knowledge of active ligands or any kind of chemical descriptors of the molecule. At the same time, the method does not rely on information about the three-dimensional structure of the protein of interest. Usually, protein three-dimensional structure determination is not an easy task. Additionally, it may be quite costly. Therefore, the usage of an amino acid sequence as input may substantially simplify one of the initial stages of drug discovery – the search for a lead compound – and can be very fruitful in the case of targeting proteins with limited or no information about inhibitors and three-dimensional structure.

To the best of our knowledge, this paper is the first attempt to present the de novo drug generation problem as a translational task between protein sequence and SMILES representation of the molecule.

The method has benefited from the recent progress in the neural machine translation field, where the Transformer architecture demonstrated state-of-the-art results [34]. Recently, Transformer also exhibited very promising results in predicting the products of chemical reactions and retrosynthesis [54,55]. One of the key features of Transformer is self-attention layers. They reduce the length of the paths that the signal should travel during deep network learning. This reduction allows the model to maintain long-range dependencies in sequence much better than in recurrent neural networks. The self-attention in Transformer architecture operates on both the input amino acid sequence and the already generated part of the SMILES string, giving access to any part of them at any time. Intuitively, self-attention is a good choice for translation between protein and molecule. First, a protein sequence may be quite long - dozens of times longer than a SMILES string. Second, three-dimensional structural features of the protein may be formed by amino acid residues located far from each other in the sequence representation. That is why it is so important for the algorithm to reference elements coming long before the current one. The multihead self-attention mechanism allows the model to jointly attend to different aspects of positions that are important in relation to proceeding elements. In language translation tasks, this ability means that Transformer may capture, for example, both the semantic and grammatical meaning of a particular word. Intuitively, it appears that this ability may be helpful in capturing 3D features of a protein binding pocket. For example, a model may consider a certain residue simultaneously in two aspects: forming the pocket and interacting directly with the drug. This is just our assumption and requires additional checking. Currently, the vast majority of deep learning approaches to the drug generation task use the similarity of organic chemistry structures and natural human language. Chemists understand molecule structure much like a human understands words. Segler et al. introduced encoder-decoder RNN architecture for the construction of a chemical language model, i.e., the probability distribution over a sequence of characters in SMILES notation [10]. Others implemented variational and adversarial autoencoders to create a continuous latent representation of chemical spaces (e.g., [22]). This creation allows easy sampling of latent codes and decoding them into SMILES strings corresponding to novel molecules. The reinforcement learning technique and fine-tuning of specific datasets were proposed to bias the probability distribution toward desired properties (e.g., [13]). In all of these approaches, the source “language” and the target “language” should ideally have the same distribution, and deep learning methods are used to construct the best fitting between them. Unlike previous studies, in our approach, we attempt to tackle the problem where source language and target language have different distributions. This approach allows the creation of a molecule with intended binding affinity using minimum information about the target, i.e., amino acid sequence only. As a proof of concept, we investigated several types of end points: chemical feasibility, physical properties, and predicted biological activity, and achieved promising results for each of them. However, the method can be improved in several directions. One of them is the generation of more diverse valid variants per protein. The Diverse Beam Search may be beneficial in this respect as it optimizes the objective containing dissimilarity term [56]. However, a more fundamental approach is to couple Transformer with a variational or adversarial autoencoder. These networks can be trained on large datasets of molecule structures to produce a latent continuous representation of chemical space. Joint training Transformer with such an additional component will allow usage of benefits from both approaches: sampling from continuous representation and conditioning on the target protein. Another improvement is to increase the number of novel structures that are not present in databases. The model learns distribution in chemical space from the training dataset and then uses it to generate a SMILES string. Typically, in deep learning, more diverse input in the training phase causes more diverse output during the generation phase. During our experiments, we noticed that the number of structures found in the ZINC15 database is lower for models trained on four organisms than for models trained only on human. Along the same lines, the Tanimoto scores between generated compounds and those from the training dataset are lower on average (0.57±0.I8 for generated molecules and 0.74±0.I4 for ones in train dataset). We anticipate that model pretraining on a much larger set of molecules (~1.5 million items from ChEMBL, for example) may substantially reduce the fraction of molecules found in databases. It also may help to increase the diversity of generated molecules from those in the training dataset. However, such improvement requires technical resources that we do not yet possess. Therefore, this optimization was out of the scope of our work. Another important improvement is an increase in the model interpretability. A visualizable interpretation may provide valuable biological insights and substantially improve understanding of the protein-ligand interaction.

## Conclusion

In this work, we introduced a deep neural network based on the Transformer architecture for protein-specific de novo molecule design. Computational experiments demonstrated the efficiency of the proposed method in terms of predicted binding affinity of generated ligands to the target protein, percentages of valid diverse structure, drug-likeness metrics and synthetic accessibility. Our model is based solely on protein sequence. This basis may be beneficial in the early stage of drug discovery, i.e., during identification of a lead compound for a protein target. The proposed method may be useful if information about the 3D protein structure is inaccessible due to difficulties in protein expression, purification and crystallization. However, our approach can be extended to yield a more interpretable model. We will address this improvement in our future studies.

## Data Availability

The code and data are available at https://github.com/dariagrechishnikova/molecule_structure_generation

## Acknowledgements

The author would like to thank L. Grechishnikov for his expertise, help in code creation, in the visualization of the figures and support. The author thanks Professor V. Tverdislov, M. Poptsova, I. Bannikova, I. Volodina, S. Pavlischev for fruitful discussions and support.

## Author contributions statement

DG conceived the presented idea, performed the computations, analyzed the results and wrote the manuscript.

