## Supplementary materials for "Transformer neural network for protein-specific de novo drug generation as a machine translation problem"

### SMILES representation for generated compounds.

1. O=C(N[C@H]1C[C@@H]1c1ccc(C(F)(F)F)cc1)N1CCN(S(=O)(=O)c2ccc(C(F)(F)F)cc2)CC1
2. COc1ccc(CN(Cc2ccc(-c3ccc(C(=O)O)cc3)cc2)Cc2ccc(C(C)(C)C)cc2)cc1
3. COc1ccc(CN(Cc2ccc(-c3ccc(C(=O)O)cc3)cc2)c2ccc(C(=O)O)cc2)cc1
4. O=C(N[C@@H]1C[C@H]1c1ccc(O)cc1)Nc1ccc(Cl)cc1
5. Cc1ccccc1NC(=O)Nc1ccc(S(=O)(=O)Nc2ccc(N3CCN(C)CC3)cc2)cc1
6. CC(C)[C@H](CC(=O)NO)C(=O)N[C@@H](CC(C)C)C(=O)N[C@@H](Cc1ccccc1)C(=O)N[C@@H](Cc1 ccccc1)C(=O)N[C@@H](CC(C)C)C(=O)N[C@@H](CC(C)C)C(=O)O
7. O=C(O)c1ccc(NC(=O)c2nc3ccccc3[nH]2)cc1
8. Cc1cc(C(=O)Nc2ccc(F)cc2)cc(-c2ccc(C(=O)O)cc2)c1
9. O=C(c1ccccc1)c1ccc(O)c(O)c1
10. COc1ccc(-c2nc(Nc3ccc(F)c(Cl)c3)c3c(n2)CCNC3)cc1
11. COc1ccc(-c2cc(=O)oc3ccccc23)cc1
12. O=C(O)c1ccc(Oc2ccc(Cl)cc2)c(Cl)c1
13. O=C(N[C@@H]1C[C@@H]1c1ccccc1)c1ccc(Cl)cc1
14. O=C(O)c1ccc(N2CCN(c3ccc4c(c3)OCO4)CC2)cc1
15. CN[C@@H]1C[C@H]2O[C@@](C)([C@@H]1OC)n1c3ccccc3c3c4c(c5c6ccccc6n2c5c31)C(=O)NC4
16. CN(C)C(=O)Nc1ccc2c(c1)S(=O)(=O)c1ccccc1N2
17. C[C@@H](NC(=O)[C@H](Cc1ccccc1)NC(=O)[C@H](Cc1c[nH]c2ccccc12)NC(=O)[C@H](Cc1ccccc1) NC(=O)[C@H](Cc1ccccc1)NC(=O)[C@H](Cc1ccccc1)NC(=O)[C@@H](C)C(N)=O)C(N)=O
18. COc1cc2c(cc1OC)C(=O)c1c(-c3ccccc3)n[nH]c1-2
19. COc1ccc([C@H]2C[C@H]3[C@@H](O)CC[C@H]3C2)cc1
20. CNC(=O)c1ccc2c(c1)[nH]c1ccccc12
21. O=C(N[C@@H]1C[C@@H]1c1ccccc1)N1CC[C@@H](c2ccc(Cl)cc2)[C@@H]1C(F)(F)F
22. COc1ccc(C(=O)Nc2ccc(Cl)cc2)cc1Cl
23. COC(=O)C(=O)N1CCN(C(=O)c2ccc(-c3ccc4c(c3)CC(=O)O4)cc2)CC1
24. COc1ccc2c(c1)nc(Nc1ccc(C(=O)O)cc1)c1ccccc1C2
25. CN(C)C(=O)c1ccc2c(c1)[C@]1(C)CC[C@@H](C2)[C@@H]1C(=O)O
26. CNC(=O)Oc1ccc2c(c1)[C@]1(C)CCN(C)[C@@H]1N2C
27. COc1ccc2[nH]cc(C)c2c1
28. COc1cc2c(cc1OC)C(c1ccccc1)N(C)C(=O)C2
29. O=C(c1ccccc1)N1CCC2(CC1)OC(=O)c1cc(O)ccc1O2
30. COc1ccc(C(=O)Nc2ccc(Cl)c(Cl)c2)cc1
31. COc1cc2c(cc1OC)C(=O)N(CC(=O)O)C2=O
32. C[C@]12O[C@H](C[C@]1(O)CO)n1c3ccccc3c3c4c(c5c6ccccc6n2c5c31)CNC4=O
33. CC(C)[C@H](NC(=O)[C@H](Cc1ccccc1)NC(=O)c1ccc(-c2ccccc2)cc1)C(=O)O
34. COc1ccc(C(=O)N2CCN(c3ccc4c(c3)OCO4)CC2)cc1
35. COc1ccc(S(=O)(=O)NC(=O)c2c(Cl)cc(Cl)cc2Cl)cc1
36. COc1ccccc1N1CCN(CCCNC(=O)c2cc(OC)c(OC)c(OC)c2)CC1
37. CNC[C@@H](O)[C@H](c1ccccc1)N(C)C
38. NC(=O)c1ccc(Oc2ccccc2)cc1
39. CN(C)C(=O)c1ccc(S(=O)(=O)N2CCN(Cc3ccc4c(c3)OCO4)CC2)cc1

40. O=C(Nc1ccc2ccccc2c1)S(=O)(=O)c1ccc2ccccc2c1
41. O=C(O)c1ccc(O)c(O)c1
42. CN(C)C(=O)c1ccc2c(c1)[nH]c1ccc(C#N)cc12
43. CC(=O)N[C@@H]1CC[C@H](C(=O)O)CC1
44. CC(=O)N1CCN(c2ccc([N+](=O)[O-])cc2)CC1
45. CC(C)(C)c1ccc(S(=O)(=O)Nc2ccc3c(c2)OCO3)cc1
46. O=C(O)c1ccc(-c2ccccc2)cc1
47. COc1cc2c(cc1OC)C(=O)c1c(-c3cnc[nH]3)c[nH]c1-2
48. CC(C)(C)c1ccc2c(c1)OCO2
49. O=C(O)c1ccc(-c2ccc(O)cc2)cc1
50. O=C(N[C@@H]1C[C@@H]1c1ccc(OCc2ccccc2)cc1)N1Cc2ccccc2C1
51. CC(C)(C)NC(=O)c1ccc(Oc2ccccc2)cc1
52. COc1ccc(-n2c(=O)n(-c3ccc(Cl)cc3)c3cc(C(=O)NS(C)(=O)=O)ccc32)cc1
53. CO[C@@H]1[C@H](N(C)C(=O)c2ccccc2)C[C@H]2O[C@]1(C)n1c3ccccc3c3c4c(c5c6ccccc6n2c5c31)C(=O)NC4
54. Cc1cc(C(=O)Nc2ccc(-c3ccccc3)cc2)nc2ccccc12
55. NC(=O)c1ccc(Oc2cn(Cc3ccc4c(c3)OCO4)nc2)cc1
56. C[C@@H]1NC(C)(C)CO[C@@]1(O)c1ccc(Cl)c(Cl)c1
57. COc1ccc(NC(=O)Nc2ccc(Cl)c(Cl)c2)c(Cl)c1
58. CC(=O)N[C@@H]1C[C@H]2O[C@@](C)([C@@H]1OC)n1c3ccccc3c3c4c(c5c6ccccc6n2c5c31)C(=O)NC4
59. O=C(O)c1ccc(Oc2ccc(-c3ccc(C(F)(F)F)cc3)cc2)cc1
60. CC(C)C[C@H](NS(=O)(=O)c1ccccc1)C(=O)O
61. COc1cccc(NC(=S)Nc2cccc(Cl)c2)c1
62. COc1cccc(N2CCN(Cc3ccccc3)CC2)c1
63. CC(=O)N[C@@H](CC(=O)O)C(=O)N[C@H](Cc1ccccc1)C(=O)N[C@H](Cc1ccccc1)C(=O)O
64. O=C(O)c1ccc(Oc2ccc(Oc3ccccc3)cc2)cc1
65. COc1ccc2c(c1)C(=Cc1ccc(C(=O)O)cc1)C(=O)N2C
66. COC(=O)c1ccc2c(c1)NC(=O)/C2=C(\Nc1ccc(F)cc1)c1ccccc1
67. O=C(Nc1nc(-c2ccc(OC(F)(F)F)cc2)cs1)c1nc(C(F)(F)F)nc(N2CCOCC2)c1Cl
68. COc1cc2c(cc1OC)CN(Cc1ccccc1)C2
69. COc1ccc(S(=O)(=O)c2ccccc2)cc1
70. O=C(O)c1ccc2c(c1)C(=O)N(Cc1ccccc1)C2
71. COc1cc(Nc2nc(Sc3ccc(NC(=O)CN4CCOCC4)cc3)nc(N3CCNCC3)n2)cc(OC)c1OC
72. O=C(O)c1ccc(OC(F)(F)F)cc1
73. COc1ccc(-c2ccc(CSc3nc(-c4ccccc4)c[nH]3)cc2)cc1
74. O=C(N[C@@H]1C[C@H]1c1ccccc1)c1ccc2c(c1)OCO2
75. COc1cccc(CNC(=O)Nc2ccc(Cl)cc2)c1
76. CCCCCCCCCCCCCCCCCCCC
77. O=C(NCc1ccccc1)c1ccc(C(F)(F)F)cc1
78. CCN(CC)C(=O)[C@H]1CC[C@H]2[C@@H]3C[C@@H]4C[C@@](O)(C4)[C@H]3CC[C@@]21C
79. COc1ccc2oc(=O)c(O)c(O)c2c1
80. O=c1[nH]c(=O)n([C@H]2C[C@@H](c3ccc4c(c3)OCO4)O2)c2ccccc12
81. N#Cc1ccc(-c2noc(-c3ccc4c(c3)OCO4)n2)cc1

82. O=C(N[C@@H](Cc1ccccc1)C(=O)N1CCC[C@H]1c1ccccc1)c1ccccc1
83. COc1ccc2nc(N3CCN(Cc4ccc(Cl)c(Cl)c4)CC3)[nH]c2c1
84. O=C(Nc1ccc(Cl)cc1)c1ccc(Cl)c(Cl)c1
85. O=C(O)c1ccc2c(c1)[nH]c1ccc(Cl)cc12
86. CC[C@H](C)[C@H](NC(=O)[C@H](Cc1c[nH]c2ccccc12)NC(=O)[C@@H]1CCCN1C(=O)[C@@H](N)C CCN=C(N)N)C(=O)N[C@@H](Cc1c[nH]c2ccccc12)C(N)=O
87. CC(C)(C)NC(=O)c1ccc(OCCCSc2nc(-c3ccccc3)no2)cc1
88. NC(=O)c1sc(-c2cccs2)nc1-c1ccc2c(c1)OCO2
89. CN(C)c1ccc(C2CCN(Cc3ccc(Cl)cc3)CC2)cc1
90. O=C(c1ccccc1)N1CCC(Nc2ncnc3ccccc23)CC1
91. COc1cc2c(cc1OC)/C(=C/c1ccc(Nc3ccccc3)cc1)C(=O)NC2
92. Cc1ccc(C(=O)Nc2cccc(CNC(=O)c3ccccc3)c2)cc1
93. COc1cc2c(cc1OC)-c1[nH]c3ccccc3c1C2
94. COc1ccc2c(c1)C(=O)Nc1ccc(-c3ccccc3)cc1C2
95. COc1ccc(CNC(=O)c2ccc3c(c2)c(C)c(C)n3C)cc1
96. O=C(N[C@@H](Cc1ccccc1)C(=O)c1nc(-c2ccccc2)c(-c2ccccc2)[nH]1)c1ccccc1
97. C[C@H]1NC(C)(C)CO[C@@]1(O)c1ccc(Cl)c(Cl)c1
98. COc1cc2c(cc1OC)-c1nc(Nc3ccc(F)cc3)ncc1CC2
99. COC(=O)c1cccc(NS(=O)(=O)c2ccc(Cl)cc2)c1
100. NC(=O)c1cc2c(Oc3ccc([N+](=O)[O-])cc3)cncc2s1
101. COc1cc2c(cc1OC)C(=O)N(CC(=O)Nc1ccccc1)C2=O
102. O=c1n(CCN2CCN(Cc3ccc(Cl)cc3)CC2)n(C2CCN(C(=O)c3ccccc3)CC2)n1C(Cl)c1ccc(Cl)cc1
103. Cc1ccc(-c2noc(C)n2)c(-c2ccc(NS(C)(=O)=O)cc2)n1
104. C[C@@H](NC(=O)c1ccc(C(F)(F)F)cc1)C(=O)N[C@H]1CCC(=O)N(c2ccc(C(F)(F)F)cc2)C1
105. CC(C)c1nc(C(=O)Nc2ccc(C(F)(F)F)cc2)oc1C(C)(C)C
106. COC(=O)c1cccc(CNC(=O)c2cc(-c3ccccc3)ccc2C(=O)Nc2ccc(Cl)cc2)c1
107. Cc1c(C#N)cccc1-c1noc(-c2ccc(C(F)(F)F)cc2)n1
108. CC(C)c1cc(NC(=O)[C@H]2CCN(c3ccc(C(F)(F)F)cc3)CC2)nc2ccccc12
109. Cc1cccc(NC(=O)c2ccc(N3CCOCC3)cc2)c1
110. C[C@]12CC[C@@H](C[C@@H]1OC)n1c3ccccc3c3c4c(c5c6ccccc6n2c5c31)CNC4=O
111. Cc1ccccc1NC(=O)c1ccc(N2CCCCC2)cc1
112. Cc1ccc(-c2ccc(C(F)(F)F)cc2)nc1
113. COc1ccc(NC(=O)[C@@H]2CCCN2C(=O)[C@@H](c2ccccc2)c2ccccc2)cc1
114. O=C(N[C@@H]1CC[C@H](O)CC1)c1ccc2c(c1)C(=O)CCC2
115. C[C@]12CC[C@@H]3c4ccc(O)cc4CC[C@H]3[C@@H]1CC[C@@H]2O
116. CN1CCN(c2ccc(Nc3ncnc4cc(-c5ccc(F)cc5)ncc34)cc2)CC1
117. Nc1ncc(-c2ccc3c(c2)OCO3)nc1-c1ccc2c(c1)OCO2
118. CC(C)(C)c1ccc(NC(=O)Nc2ccc(C(=O)O)cc2)cc1
119. CC(C)NC(=O)c1ccc2c(c1)[C@@]1(C)CC[C@H](O)[C@@H]1N2C
120. COc1cc(NC(=O)C(C)(C)C)ccc1C(=O)Nc1ccc(Cl)cc1
121. NC(=O)c1cccc(OC[C@H]2CN3CCC[C@@H]3C2)c1
122. NC(=O)c1ccc(-c2ccccc2Cl)cc1
123. COc1ccc([C@H]2C[C@H](O)[C@@H](O)[C@H]2O)cc1
124. O=C1NC(=O)C(c2ccc3ccccc3c2)C1=O

125. NC(=O)c1ccc(Nc2nccc(-c3ccc4c(c3)NC(=O)C4=O)n2)cc1
126. COc1ccc(-c2ccc(CC(=O)O)cc2)cc1
127. O=C(Nc1ccc(Oc2ccccc2)cc1)c1cnc2ccccc2c1
128. CN(C)C(=O)c1ccc(Oc2ccccc2Cl)c(Cl)c1
129. COc1cc2c(cc1OC)C(O)C(=O)N(C)C(=O)c1ccc(C(F)(F)F)cc1S2(=O)=O
130. Nc1ncc(C(=O)Nc2ccc(S(N)(=O)=O)cc2)s1
131. CN(C)C(=O)c1ccc2c(=O)n(Cc3ccccc3)c(=O)c2c1
132. C[C@@H]1N(C)CCN1c1ccccc1Cl
133. COc1ccc(-c2c(-c3ccccc3)nc(C)n2-c2ccc(Cl)cc2)cc1
134. COC(=O)c1ccc2c(c1)[C@@]1(C)CCN(C(=O)c3ccccc3)[C@@H]1C2
135. COc1ccc2c(c1)C(C(=O)Nc1ccc(Cl)cc1)C(=O)N2
136. NS(=O)(=O)c1ccc(-c2ccc(N3CCNCC3)cc2)cc1
137. CC(C)(C)C(=O)N[C@@H](Cc1c[nH]c2ccccc12)C(=O)N[C@@H](Cc1ccccc1)C(=O)O
138. COc1ccc(-c2n[nH]c3c2C(=O)c2ccc(-c4ccccc4)cc2-3)cc1
139. COc1ccc(C2=C(c3ccc(F)cc3)c3ccccc3N(C)C2=O)cc1
140. CC(C)(C)c1nn(-c2ccc(F)cc2Cl)c2c1C(=O)CCC2
141. CC(C)[C@H](NC(=O)[C@H](Cc1c[nH]c2ccccc12)NC(=O)OCc1ccccc1)C(=O)O
142. O=C(Nc1nc2ccccc2s1)c1ccccc1
143. CC(C)[C@H](NC(=O)[C@H](Cc1ccccc1)NC(=O)[C@H](Cc1ccccc1)NC(=O)[C@H](Cc1ccccc1)NC(=O)
OCc1ccccc1)C(=O)O
144. COc1ccc(S(=O)(=O)N2CCN(C(=O)c3ccc4c(c3)OCO4)CC2)cc1
145. CC(C)[C@H](NC(=O)c1ccccc1)C(=O)N[C@H]1C[C@H]2O[C@](C)([C@@H]1OC)n1c3ccccc3c3c4c(c
5c6ccccc6n2c5c31)C(=O)NC4
146. O=C(Nc1ccccc1)N1CCC(N2CCCC2)CC1
147. Cc1ccc(-c2sc(C(F)(F)F)nc2C(=O)c2ccc(Cl)cc2)cc1
148. CC(C)(C)c1cnc(N2CCN(c3ccc(C(F)(F)F)cc3)CC2)s1
149. COC(=O)c1ccc2[nH]ncc2c1
150. O=C(Nc1nc(-c2ccccc2)ns1)c1ccc(N2CCCCC2)cc1
151. O=C(O)c1ccc(OCCN2CCN(c3cccc(Cl)c3)CC2)cc1
152. COc1cc(NC(=O)C(C)(C)C)ccc1C(=O)Nc1ccc(Cl)c(Cl)c1
153. NC(=O)c1ccc2c(c1)C(=Nc1ccc(S(N)(=O)=O)cc1)C(=O)N2
154. C[C@H]1CC[C@@H](NC(=O)Nc2cccc(C(F)(F)F)c2Cl)CC1
155. Nc1ncnc(N2CCC(Nc3ncnc4[nH]cc(-c5ccc(C(F)(F)F)cc5)c34)CC2)n1
156. C[C@]12CC[C@H]3[C@@H](CC[C@H]4CCCC[C@@]43C)[C@@H]1CC[C@@H]2O
157. Nc1nc2ccccc2c(=O)[nH]1
158. COc1ccc(Nc2nccc(-c3ccc4c(c3)NC(=O)C4=O)n2)cc1
159. COC(=O)c1cccc(-c2nc(NC(C)=O)c(NC(C)=O)o2)c1
160. COc1ccc(-c2c(C)noc2-c2ccc3nc(N)n(C)c3c2)cc1
161. COc1ccc(CNC(=O)c2cc(-c3ccc4c(c3)OCO4)n[nH]2)cc1
162. COc1cc2c(cc1OC)-c1[nH]nc(-c3ccc(N4CCOCC4)cc3)c1C2
163. Cc1ccc(-c2cc(C(F)(F)F)ccc2C(=O)Nc2ccc(-c3ccccc3)cc2)cc1
164. NC(=O)c1ccc2nc(-c3ccc(NC(=O)c4ccc5c(c4)OCO5)cc3)[nH]c2c1
165. C[C@@H](NC(=O)[C@H](Cc1ccccc1)NC(=O)[C@H](CC(C)C)NC(=O)[C@H](Cc1ccccc1)NC(=O)OC(C)
(C)C)C(C)C

166. C[C@@H]1C[C@H]2[C@@H]3CCC4=CC(=O)CC[C@]4(C)[C@@H]3CC[C@@]21C
167. COc1ccc(C(=O)Nc2ccc(-c3ccccc3)cc2)cc1
168. COc1ccc(-n2nnnc2C(=O)c2ccc(-n3cnc4ccccc43)cc2)cc1
169. COc1ccc(S(=O)(=O)N2CCN(C(=O)N3CCN(C)CC3)CC2)cc1
170. Cc1ccccc1Oc1ccc(-c2ccccc2)nc1
171. COc1ccc(S(=O)(=O)N2C[C@@H](OC)[C@@H](OC)C2)cc1
172. O=C(Nc1ccc(Cl)cc1)N1CCN(Cc2ccccc2)CC1
173. O=C(Nc1nc2ccccc2s1)c1ccc2c(c1)NC(=O)C2
174. CC(C)[C@H](NC(=O)[C@H](Cc1ccccc1)NC(=O)OCc1ccccc1)C(=O)O
175. Cc1ccccc1S(=O)(=O)N1CCN(C2CCCCC2)CC1
176. CC(C)[C@H](NC(=O)c1ccc(-c2ccccc2)cc1)C(=O)O
177. O=C(Nc1nnc(-c2ccccc2)s1)c1ccccc1
178. NC(=O)c1ccc(Nc2nccc(-c3ccc4c(c3)OCO4)n2)cc1
179. COc1ccc(CNC(=O)c2cc3c(cc2OC)C(=O)N(C)C3=O)cc1
180. CC(C)(C)NC(=O)c1ccc(NS(=O)(=O)c2ccc(C(F)(F)F)cc2)cc1
181. CN1CCN(c2ccc(-c3cnc4[nH]c5c(F)cccc5c4c3)cc2)CC1
182. NC(=O)c1ccc2c(c1)nc(NCc1ccc(Cl)cc1)c1cc(C(N)=O)ccc1C2
183. COc1ccc2c(c1)[C@H]1CC[C@H](C2)N1C
184. O=C(O)c1ccc(Nc2nccc(-c3ccccc3)n2)cc1
185. O=C(Nc1ccc(-c2ccccc2)cc1)N1CCC(Oc2ccc(Cl)cc2)CC1
186. NC(=O)c1cnc(-c2ccc3c(c2)CNC3=O)[nH]1
187. CNC(=O)[C@@H]1C[C@H]2O[C@@](C)([C@@H]1OC)n1c3ccccc3c3c4c(c5c6ccccc6n2c5c31)C(=O
)NC4
188. C[C@]12CC[C@H]3[C@@H](CC[C@H]4CCC(=O)CC[C@@]43C)[C@@H]1CC[C@@H]2O
189. COc1cc(Nc2ncc3ccc(F)cc3n2)ccc1F
190. COc1ccc(Cl)c(Cl)c1C1c2c(c(OC)c1OC)n(C)n2C(=O)c1ccc(F)cc1
191. COc1cc2c(cc1OC)C(=O)C(C(=O)Nc1ccccc1)C(=O)N2
192. COc1ccc2c(c1)C(=O)C(CC1CCN(Cc3ccccc3)CC1)C2
193. COc1cc2c(cc1OC)C(=O)C(C)(C)CCC2NC(=O)c1ccc(-c2ccc(C(=O)O)cc2)cc1
194. O=C(O)c1cccc(Nc2nc3cccc(C(F)(F)F)c3c(=O)[nH]2)c1
195. CCCCCCCCCCCC(=O)NCCCCCCCC(=O)OC(C)C
196. CCCCCCCCCCCCCCCCCCCCCCCCCCCC(=O)O
197. COc1ccc(C[C@]2(c3ccccc3)[C@@H](c3ccccc3)N[C@@H](C)C2)cc1
198. COc1cccc(C(=O)Nc2cccc(C(F)(F)F)c2)c1
199. COc1ccc(NC(=O)NCCc2ccccc2)cc1
200. COc1ccc(CN(C(=O)c2ccc(N(C)C(F)(F)F)cc2)C(C)(C)C(=O)O)cc1
201. COc1ccc2c(c1)C(=O)C(C)(C)N2c1ccc(S(N)(=O)=O)cc1
202. COc1cc(NC(=O)[C@@H](C)NC(=O)c2ccccc2Cl)ccc1C(F)(F)F
203. COc1cc(N2CCN(C(=O)OC(C)(C)C)CC2)ccc1S(C)(=O)=O
204. Cc1cc(C(=O)Nc2cccc(C(F)(F)F)c2)cc(N2CCC(N3CCOCC3)CC2)n1
205. COc1ccc2c(c1)C(=O)/C2=C\c1ccc(C(=O)O)cc1
206. COc1cc2c(cc1OC)C(=O)N(Cc1ccccc1)CC2
207. O=C(O)c1ccccc1-c1ccc2c3[nH]ncc3c(=O)n(CC(F)(F)F)c2c1
208. COc1ccc2c(c1)C(=O)/C(=C/c1ccc(S(N)(=O)=O)cc1)O2

209. O=C(Nc1ccc(N2CCN(C(=O)c3ccc(Cl)c(Cl)c3)CC2)cc1)N1CCOCC1
210. O=C(Nc1ccc(Cl)c(Cl)c1)c1ccc(NS(=O)(=O)c2cccs2)cc1
211. O=C(Nc1ccc(Cl)c(Cl)c1)c1ccc(Cl)c(Cl)c1
212. Cc1cc(-c2ccc3c(c2)OC[C@@H](c2ccccc2C(F)(F)F)[C@@H](O)[C@@H]3O)ccc1C(F)(F)F
213. COc1ccc(NC(=O)/C=C2\C(=O)Nc3ccccc32)cc1
214. COc1cc2c(cc1C(=O)Nc1ccccc1)C(=O)N2
215. O=C(NC1CCN(Cc2ccc(C(F)(F)F)cc2)CC1)c1ccc(F)cc1
216. COc1ccccc1N1CCN(C(=O)c2c(-c3ccccc3)[nH]c3ccccc23)CC1
217. COc1ccccc1N1CCN(CCCNC(=O)c2nc(C)n(-c3ccccc3)c2C)CC1
218. O=C(O)c1ccc(C(=O)OCc2ccccc2)cc1
219. CC(C)(C)c1cccc(NC(=O)Nc2cccc(C(F)(F)F)c2)c1
220. O=C(O)c1ccc(C(F)(F)F)cc1-c1ccc2c(c1)OCO2
221. COc1ccc(C(=O)N2CCC(CNC(=O)c3c(Cl)cc(Cl)cc3Cl)CC2)cc1
222. COc1ccc(S(=O)(=O)Nc2ccc(Cl)c(Cl)c2)cc1
223. CC(C)(C)c1ccc(NC(=O)Nc2ccc(C(F)(F)F)cc2)cc1
224. O=C(N[C@@H]1CC[C@H](N2CCCCC2)C1)c1ccc(-c2nc3ccccc3s2)cc1
225. O=C(N[C@@H]1COCC[N+]1)c1ccc(F)cc1
226. COc1ccc(-c2nc(NC(=O)c3ccc(N(C)C)cc3)c3ccc(F)cc3n2)cc1
227. N#Cc1ccc(C(=O)Nc2ccccc2-c2ccccc2)cc1
228. CC(C)(C)c1ccccc1NC(=O)C1CC1
229. O=C(NC1CCCCC1)c1ccc(-c2ccccc2)cc1
230. COc1ccc(NC(=O)Nc2ccc(C(F)(F)F)cc2)cc1
231. COc1ccc2c(c1)C(=O)C(CC1CCCCC1)C2
232. O=C(Nc1ccc(Cl)cc1)N1CCC(C(O)(c2ccc(Cl)cc2)c2ccc(Cl)cc2)CC1
233. O=C(O)c1cc(-c2ccncc2)cc(NC2CCN(CCO)CC2)n1
234. Cc1ccccc1N1CCC(CNC(=O)c2ccccc2Cl)CC1
235. O=C(Nc1ccc(Cl)cc1)N1CCC(Nc2nccc(-c3ccccc3)n2)CC1
236. COc1cc(Nc2ncc3c(n2)C(=O)/C3=C\c2ccccc2)cc(OC)c1OC
237. CC(=O)N1CCN(C(=O)N2CCCCC2)CC1c1ccc(Cl)cc1Cl
238. O=C(O)C1=C(c2ccccc2)C(=O)N(Cc2ccccc2)C1=O
239. O=C(Nc1ccc(-c2ccccc2)cc1)[C@H]1CCCN(Cc2ccccc2)C1
240. COc1ccccc1C(=O)Nc1ccccc1
241. COc1cc(N2CCN(C)CC2)cc(Nc2ncc(Br)c(N[C@H](C)C(=O)O)n2)c1
242. O=C(Nc1ccc(C(F)(F)F)cc1)[C@H]1C[C@@H]2[C@@H](O)[C@@H](O)[C@H]2O1
243. COc1ccc(NC(=O)Nc2cccc(C(F)(F)F)c2)cc1
244. COc1cc(Nc2nc(Nc3ccc(Cl)c(Cl)c3)nc(Nc3ccc(N4CCN(C)CC4)cc3)n2)ccn1
245. O=C(Nc1ccccc1)N1CCN(Cc2ccccc2)CC1
246. COc1ccc(S(=O)(=O)N2CCN(C(=O)C3CC3)CC2)cc1
247. C[C@]12CC[C@H]3[C@@H](C[C@@H](O)C4CC4)C[C@@H]3[C@@H]1CCC2=O
248. COc1ccc2c(c1)C(C)(C)C2
249. COc1ccccc1NC(=O)CN1CCN(c2nc3ccccc3s2)CC1
250. COc1cc2c(cc1OC)C(=O)C(C(=O)O)C(=O)N2
251. N#Cc1ccc(NC(=O)Nc2ccc(C(F)(F)F)cc2)cc1
252. COc1ccc2c(c1)NC(=O)/C(=C/c1ccc(N(C)C)cc1)C(=O)N2

253. COc1ccc(-c2cc3c(c(=O)n2-c2ccc(F)cc2)CNC3)cc1
254. O=C1NC(=O)c2cc(-c3ccc(O)cc3)ccc2N1c1ccc(Cl)cc1
255. CC(C)(C)OC(=O)N[C@@H]1CCC[C@H]1NC(=O)OCc1ccccc1C(F)(F)F
256. COc1cc(Nc2ncc(C(=O)Nc3cccc(C(F)(F)F)c3)s2)ccc1N1CCN(C)CC1
257. COc1cccc(C(=O)Nc2ccccc2C(=O)Nc2ccc(C(=O)O)cc2)c1
258. COc1ccc(Nc2ncc3c(c2)NC(=O)/C3=C\c2ccc(OC)cc2)cc1
259. COc1ccc(C[C@H](C)NC(=O)c2ccccc2)cc1
260. COc1ccc(-c2nn(CC(=O)O)c3ccc(F)cc23)cc1OC
261. NS(=O)(=O)c1ccc(C(=O)NCc2ccc(C(F)(F)F)cc2)cc1
262. COc1ccccc1N1CCN(C(=O)c2cc3ccccc3[nH]2)CC1
263. COc1cc(C(=O)N[C@@H]2CC[C@H](C(=O)O)CC2)ccc1OC
264. CC(C)(C)c1nc(C(=O)Nc2nc(C(F)(F)F)cs2)cs1
265. CC(C)C[C@H](NC(=O)[C@H](Cc1ccccc1)NC(=O)[C@@H]1CCCN1C(=O)[C@H](CCCNC(=N)N)NC(=O)[C@H](CCCN)NC(=O)[C@@H](N)Cc1ccccc1)C(N)=O
266. O=C(O)c1ccc(Cl)cc1S(=O)(=O)Nc1ccc(Cl)cc1
267. O=C(O)C1=C(c2ccc(Oc3ccc(Cl)cc3)cc2)CCC1
268. COc1ccc2c(c1)C(=O)C(C)(C)C(=O)N2c1ccccc1
269. O=S(=O)(c1ccc(Cl)cc1)C1CCN(S(=O)(=O)c2ccccc2)CC1
270. O=C(O)c1ccc(C(=O)Nc2ccc(C(F)(F)F)cc2)cc1
271. O=C(Nc1ccc(Cl)c(Cl)c1)N1CCCC1
272. CCCS(=O)(=O)c1ccc(CN(Cc2ccc(OC)c(OC)c2)CC2CCCCC2)cc1
273. O=C(O)c1ccc(NC(=O)Nc2ccc(C(F)(F)F)cc2)cc1
274. O=C(N[C@@H]1C[C@H]2O[C@@H](n3cnc4ccccc43)[C@@H](O)[C@@H]2[C@@H]1O)c1ccccc1
275. COc1cc(Nc2ncc(Br)c(Nc3ccc(C(=O)O)cc3)n2)ccc1N1CCN(C)CC1
276. O=C(N[C@H]1CC[C@H](N2CCCC2)[C@H](O)C1)c1ccc(F)cc1
277. COc1ccc2c(c1)C(=O)N(C)/C2=C\c1ccc(/C=C/C(=O)O)cc1
278. COc1ccc(C(=O)Nc2ccc(-c3ccc(OC)cc3)cc2)cc1
279. COc1cccc(C(=O)Nc2ccccc2C(=O)Nc2cccc(C(=O)O)c2)c1
280. Cc1cc(C(F)(F)F)ccc1-c1cnc2c(c1)OCO2
281. O=C(O)c1ccc(Nc2nc3cc(Cl)ccc3s2)cc1
282. COc1cc(Nc2ncc3c(n2)C(c2ccccc2C(F)(F)F)CC3)ccc1C(N)=O
283. COc1ccc2c(c1)[C@]1(C)CCN(C(=O)OC(C)(C)C)[C@@H]1N2C
284. O=C(N[C@@H](Cc1ccccc1)C(=O)N1CCC[C@H]1c1ccccc1)C(=O)N1CCC[C@H]1c1ccccc1
285. COc1cc(-c2cnc(N)s2)cc(C(=O)O)c1
286. CC(C)(C)c1ccc(NC(=O)c2ccccc2)cc1
287. CC(C)(C)c1ccc(NC(=O)Nc2ccc(-c3cnc(N)nc3)cc2)cc1
288. COc1cc(-c2ccc(OC)cc2)cc(-c2ccccc2)c1
289. COc1ccc(-c2ccc(C(=O)O)c(C)c2)cc1
290. O=C(O)c1cc(-c2ccccc2)cc(-c2ccccc2)c1
291. CCCCCCCCCCCCCCCNC(=O)[C@@H]1O[C@@H](C(=O)N[C@H]2CCC(C)(C)C2)[C@H](O)[C@H]1O
292. NS(=O)(=O)c1ccc(NC(=O)c2ccccc2)cc1
293. COC(=O)c1cccc(NC(=O)C2(N)CCN(c3cccc(Cl)c3)CC2)c1
294. CC(C)(C)OC(=O)Nc1ccc(S(=O)(=O)Nc2cccc(Cl)c2)cc1

295. CC(C)C[C@H]1NC(=O)[C@H](Cc2ccccc2)NC(=O)[C@H](Cc2ccccc2)NC(=O)[C@H](Cc2ccccc2)NC(=O)[C@H](Cc2ccccc2)NC1=O
296. C[C@@H]1C[C@H]2O[C@@](C)([C@@H]1OC)n1c3ccccc3c3c4c(c5c6ccccc6n2c5c31)C(=O)NC4
297. Cc1ccc(-c2ccc(C(=O)Nc3ccc(C(=O)N(C)C)cc3)cc2)cc1
298. COC(=O)c1ccc(Sc2cnc(C(=O)N3CCN(c4ccccc4)CC3)o2)cc1
299. O=c1[nH]c2ccccc2[nH]1
300. O=C(Nc1cc(-c2ccc3c(c2)OCO3)ccc1O)c1ccc(F)cc1
301. O=C(NO)c1ccc2c(Nc3ccc(Cl)cc3)ncnc2c1
302. C[C@]12CC[C@H]3c4cc(Cl)ccc4CC[C@H]3[C@@H]1CC[C@@H]2NS(=O)(=O)c1ccccc1
303. CC(C)C[C@H]1O[C@@H](O)[C@@H](O)[C@H](O)[C@H](O)[C@H](O)[C@H]1O
304. COc1ccc(-c2nc3nc(-c4ccc(OC)cc4)oc3n3ccnc23)cc1
305. COc1ccc2[nH]cc(CCN3CCC(c4ccccc4)CC3)c2c1
306. O=C(O)C[C@H]1[C@@H](O)CO[C@H]1[C@@H](O)CC[C@@]1(O)CC[C@@H](O)[C@H]1O
307. COc1ccc(C(=O)N[C@@H]2C[C@@H](O)[C@@H](O)[C@@H]2O)cc1
308. Cc1cc(-c2cc(C(F)(F)F)nc(-c3ccc4c(c3)OCO4)n2)nc(-c2ccc(Cl)c(Cl)c2)n1
309. COc1ccc(-n2nnnc2C)cc1-c1ccc(S(=O)(=O)N2CCCCCC2)cc1
310. CC(C)(C)c1ccc(S(=O)(=O)N2CCN(c3cccc(C(F)(F)F)c3)CC2)cc1
311. CC(C)(C)c1cnc(C2CC2)nc1
312. COc1ccc(-c2nc(C(=O)Nc3ccc(OC)cc3)[nH]c2-c2ccc(OC)cc2)cc1
313. O=C1NC(=O)/C(=C/c2cnn3c(N4CCNCC4)cc(-c4ccc(C(F)(F)F)cc4)cc23)N1
314. CS(=O)(=O)c1ccc(NC(=O)NC2CCCCC2)cc1
315. CC(C)(C)OC(=O)N1CCC(c2cc(-c3ccc4nc(-c5nnc(N)o5)[nH]c4c3)ccc2C#N)CC1
316. COc1ccc(S(=O)(=O)N2CCN(C(=O)c3ccc(C(F)(F)F)cc3)CC2)cc1
317. CN1C(=O)[C@]2(C)Oc3ccccc3N(C)C12
318. O=C(Nc1ccc(-c2ccccc2)cc1)c1ccc(O)cc1
319. CC(C)(C)OC(=O)N1CC=C(c2c[nH]c3ncc(Cl)cc23)CC1
320. O=C(Nc1cc(-c2ccccc2)ncn1)c1ccccc1
321. COc1cc(-c2ccncc2)cc(C(=O)O)c1
322. O=C(O)c1ccc(Nc2nccc(-c3cccc(N4CCOCC4)c3)n2)cc1
323. COc1ccc(C(=O)Nc2ccc(C(=O)O)cc2)c(Oc2ccc(Cl)cc2)c1
324. O=C(O)c1ccc(-c2ccc(-c3ccc(F)cc3)cc2)cc1
325. COc1ccc(NC(=O)N[C@@H]2Cc3ccccc3O2)cc1
326. COc1cc2c(cc1OC)C(=O)C1=C(C(=O)O)C(C(=O)N(C)C)CC1C2
327. O=C(O)c1ccc(C2CC2)cc1-c1ccc2c(c1)CCCC2
328. O=C(O)c1ccc(-c2ccc3c(-c4ccccc4)c[nH]c3c2)cc1
329. CN1CCN(c2ccc(-c3c(-c4ccccc4)c[nH]c3-c3ccccc3)cc2)CC1
330. O=C(N[C@@H]1C[C@@H](O)[C@H]1OCc1ccc(F)cc1)c1ccc(Cl)cc1
331. O=C(N[C@@H]1CC[C@H]1OCc1ccccc1)c1ccccc1
332. O=C(N[C@@H]1C[C@H](O)CC[C@@H]1OCc1ccccc1)c1ccccc1
333. O=C(Nc1ccc(F)cc1)Nc1ncc(Cl)c(Nc2ncnc3ccccc23)n1
334. O=C(NCC#Cc1ccc2ncnc(Nc3cccc(F)c3)c2c1)c1cccc(F)c1
335. O=C(N[C@@H]1CNC[C@H]1O)c1ccc2c(c1)[nH]c1ccccc12
336. O=C(N[C@@H](Cc1ccccc1)C(=O)Nc1ccccc1)c1ccccc1
337. COc1ccc(-c2nc([C@H]3CC[C@H](NC(=O)C4CCCCC4)CC3)c(C(F)(F)F)[nH]2)cc1

338. O=C(O)c1ccc2c(c1)C(c1ccccc1)CCC2
339. CC(C)(C)c1ccc(C(=O)NCCCN2CCN(c3nc4ccccc4s3)CC2)cc1
340. CC(C)(C)OC(=O)c1ccc(-c2ccc(C(F)(F)F)nc2)cc1
341. O=C(N[C@@H]1C[C@H](O)CN1)c1cc(F)c(F)c(F)c1
342. Cc1cc(-c2ccc(F)c(F)c2)c(-c2ccc(F)c(C(F)(F)F)c2)o1
343. COc1cc2c(cc1OC)C(=O)N(C)C(=O)N2
344. O=c1[nH]c(=O)[nH]c2ccccc12
345. CN1CCN(c2ccc(C(=O)Nc3ccc(Cl)cc3Cl)cc2)CC1
346. CC(C)(C)c1ccc(S(=O)(=O)N2CCN(c3nc4ccccc4[nH]3)CC2)cc1
347. COc1cc(C(=O)Nc2ncc(-c3ccc(C(=O)O)cc3)s2)ccc1Cl
348. CN1CCN(c2ccc(C(F)(F)F)cc2)CC1
349. CCCCCCCCCCCCCCCCCCCCP(=O)([O-])O[N-]
350. COc1cc2c(cc1OC)C(=O)N(C)c1ccc(-c3ccc(C(F)(F)F)cc3)cc1C2
351. O=C1CC2(CCCC2)[C@@H](O)[C@@H]1O
352. CC(C)[C@H]1NC(=O)[C@H](CC(=O)O)NC(=O)[C@H](Cc2ccccc2)NC(=O)[C@H](Cc2ccccc2)NC(=O)[C@H](Cc2ccccc2)NC(=O)[C@H](Cc2ccccc2)NC1=O
353. COc1ccc(-c2cc(C(=O)O)ccc2Cl)cc1
354. CC(C)(C)c1ccc(NC(=O)c2ccc(-c3ccccc3C(=O)O)cc2)cc1
355. CC(C)C[C@H](NC(=O)[C@H](Cc1ccccc1)NC(=O)[C@H](CC(=O)O)NC(=O)[C@H](Cc1ccccc1)NC(=O)OCc1ccccc1)C(=O)O
356. CC(C)(C)c1ccc(-c2ccccc2C(=O)O)cc1
357. O=C(NC1CC1)c1ccc(Nc2ncc3ccc(Cl)cc3n2)cc1
358. COc1ccc(C(=O)Nc2ccc(C(F)(F)F)cc2)cc1C(=O)Nc1ccc(C(F)(F)F)cc1
359. Cc1cc(C(F)(F)F)cc(C(F)(F)F)c1C(=O)N[C@@H]1CCCC[C@H]1Nc1ccccc1
360. CC(C)(C)OC(=O)N[C@H](Cc1ccccc1)C(=O)N[C@@H](CCCN=C(N)N)C(=O)O
361. CC(C)C[C@@H]1O[C@@H](c2ccccc2)[C@H](O)[C@@H]1O
362. Cc1cc(Cl)cc(Cl)c1C(=O)N[C@H]1CC[C@@H](CO)O1
363. O=C(N[C@@H]1CC[C@H](O)[C@H]1OC(=O)[C@H](Cc1ccccc1)NC(=O)OCc1ccccc1)C(F)(F)F
364. O=C(O)c1ccc(-c2ccc3c(c2)OCO3)cc1
365. COc1cc2c(cc1O)CCC2=O
366. CC(C)(C)OC(=O)N[C@@H]1C[C@H](Oc2ccc(-c3ccccc3C(=O)O)cc2)C[C@H]1O
367. CC(C)(C)OC(=O)c1ccc(NC(=O)[C@H]2CC[C@H](NC(=O)[C@H]3C[C@@H](N)CC3)CC2)cc1
368. O=C(Nc1ccc(Oc2ccc(O)cc2)cc1)c1ccccc1
369. O=C(c1ccc(Cl)cc1)N1CCN(C2CCCCC2)CC1
370. CCCCCCCCCCCCCCCCCCCCP(=O)(O)O
371. O=C(O)c1ccc(Cc2cc(Cc3ccccc3)[nH]n2)cc1
372. COc1cc(NC(=O)[C@@H](Cc2ccc(OC)cc2)NC(=O)c2ccccc2)ccn1
373. COc1cc(N2CCC(N(C)C)CC2)ccc1Nc1ncc(Cl)c(Cl)c1Cl
374. COc1ccc(S(=O)(=O)Nc2ccc(C(F)(F)F)cc2)cc1
375. COc1cc(Nc2c(C#N)cnc3cc(OC)c(O)cc23)cc(OC)c1OC
376. COc1cc(COc2ccc([C@@H](O)C(=O)O)cc2)cc(C(=O)O)c1O
377. CC(C)(C)OC(=O)N[C@@H](Cc1c[nH]c2ccccc12)C(=O)N[C@@H](C(=O)O)C(C)C
378. COc1ccc(NC(=O)c2cncc(N)n2)cc1Cl
379. c1ccc2c(N3CCNCC3)nccc2c1

380. CC(=O)N[C@@H]1CC[C@H]2O[C@@](C)([C@@H]1OC)n1c3ccccc3c3c4c(c5c6ccccc6n2c5c31)C(=O)NC4
381. O=C(N[C@H]1CC[C@H](c2ccccc2)CC1)c1ccc(N2CCOCC2)nc1
382. COc1cc(C(=O)Nc2ccc([N+](=O)[O-])c([N+](=O)[O-])c2)c(OC)c(OC)c1OC
383. Cc1cnc(C(=O)Nc2cc(C(F)(F)F)cc(C(F)(F)F)c2)c(C)c1
384. CC(=O)N[C@@H](CC(C)C)C(=O)N[C@@H](Cc1ccccc1)C(=O)N[C@@H](CC(C)C)C(=O)N[C@@H](CC(C)C)C(=O)O
385. COc1ccc(S(=O)(=O)N2CCN(C(=O)OCc3ccc(Cl)cc3)CC2)cc1
386. COc1ccc(S(=O)(=O)N2CCC(Oc3ccc(Cl)cc3)CC2)cc1
387. O=C(O)[C@H](Cc1ccccc1)[C@H](O)C(=O)N(Cc1ccc(Cl)cc1)C1CC1
388. O=C(Nc1cc(-c2ccc(O)cc2)nc2ccccc12)c1ccccc1
389. CC(=O)N[C@@H](Cc1ccccc1)C(=O)N[C@@H](Cc1ccccc1)C(=O)N[C@@H](CC(C)C)C(=O)O
390. COC(=O)[C@H](Cc1ccccc1)NC(=O)[C@H](Cc1c[nH]c2ccccc12)NC(=O)[C@H](Cc1c[nH]c2ccccc12)NC(=O)OC(C)(C)C
391. CC(C)(C)c1cc(C(=O)N[C@@H](CC(=O)O)C(=O)N[C@@H](CC2CCCCC2)C(=O)N[C@@H](CC(C)C)C(=O)N[C@@H](CC(C)C)C(=O)O)ccc1Cl
392. O=C(O)c1c(-c2ccccc2)c2ccccc2n1C1CC1
393. COc1ccc2c(c1)C(c1ccccc1)=NCCN2
394. COC(=O)c1ccc(NC(=O)Nc2ccc([N+](=O)[O-])cc2)cc1
395. COc1ccc(S(=O)(=O)Nc2cccc(C(=O)O)c2)cc1OC
396. COc1cc(N2CCOCC2)cc(C)c1-n1c(C)nnc1C
397. CCOC(=O)c1ccc(NC(=O)N[C@@H](CC2CCCCC2)C(=O)O)cc1
398. CCN(C)C/C=C/C(C)=C/C(=O)OC(C)(C)C
399. COc1ccc(S(=O)(=O)N2CCC[C@H]2C(=O)N(C)C)cc1OC
400. COc1ccc(S(=O)(=O)N(Cc2ccccc2)c2ccccc2)cc1
401. Cc1ccc(NC(=O)C(C)(C)C)cc1-c1ccc(S(=O)(=O)Nc2ccccc2C(F)(F)F)cc1
402. COc1cc(NC(=O)c2ccc(OC(F)(F)F)cc2)cc2c1O[C@@H]1O[C@@H](C(F)(F)F)[C@@H](O)[C@]12C
403. COc1ccc(-c2cc(C(=O)O)cc(C(=O)O)c2)cc1C(=O)O
404. C[C@@H]1C[C@H]2C[C@H](CC[C@@H]3[C@@H](O)CC[C@@H]32)[C@H]2C[C@@H]1CC[C@]2(C)C
405. CN(C)C(=O)c1ccc2c(c1)Nc1ccccc1C2=O
406. CC(=O)N[C@@H](Cc1ccccc1)C(=O)N[C@@H](CC(C)C)C(=O)N[C@@H](Cc1ccccc1)C(=O)O
407. C[C@]12O[C@H](C[C@]1(O)CO)n1c3ccccc3c3c4c(c5c6ccccc6n2c5c31)C(=O)NC4
408. COc1cc(Nc2ncnc3cc(-c4ccc(OC)cc4)[nH]c23)cc(OC)c1OC
409. O=C(O)c1ccc(C(=O)N2CCC(CCc3ccccc3)CC2)cc1
410. CCCCCCCCCCCCC(=O)C(F)(F)F
411. N#Cc1ccc2c(C3CC3)n[nH]c2c1
412. Cc1cc(NC(=O)c2ccc(Cl)cc2)c2ccccc2n1
413. CC(C)[C@H](NC(=O)[C@@H](Cc1ccccc1)NC(=O)OCc1ccccc1)C(=O)N[C@@H](CC1CCCCC1)C(=O)O
414. Cc1nc2ccccc2c2c(=O)[nH]c12
415. COc1ccc(-c2ccc(N3CCC(C(=O)O)CC3)cc2)cc1
416. O=C1c2ccccc2C(=O)N1CC1CCN(Cc2ccc(C(F)(F)F)cc2)CC1
417. COc1cccc(-c2cc(COc3ccc4c(c3)OCO4)c(CO)c(OC)c2OC)c1
418. Cc1nnc(N2CCC(N3CCN(C)CC3)CC2)c(N2CCN(C(=O)Nc3cccc(Cl)c3Cl)CC2)n1

419. COc1ccc(-c2nc(N3CCN(C)CC3)c(C(=O)O)[nH]2)cc1
420. COc1ccc(C2=C(N3CCN(C(=O)OC)CC3)C(=O)OC2c2ccccc2C(=O)O)cc1
421. C[C@@H]1OC(=O)n2c(C#N)cnc2Nc2ccc(Cl)cc2S(=O)(=O)N1C
422. C[C@H]1C[C@@H](NC(=O)c2ccc(OC(F)(F)F)c(C(F)(F)F)c2)[C@H](c2ccc(Cl)cc2)C1
423. COc1cccc(NC(=O)[C@H](Cc2ccccc2)NC(=O)[C@H](Cc2ccccc2)NC(=O)OCc2ccccc2)c1
424. O=C(O)c1ccc2c(O)cccc2c1O
425. CC(C)[C@@H](Oc1ccccc1)C(=O)N1CCCCC1
426. Cc1cc(C(=O)N[C@@H](Cc2ccccc2)C(=O)N[C@H](Cc2ccccc2)C(=O)N[C@H](C(=O)O)C(C)C)cc(O)c1O
427. COc1cc(NC(=O)c2cccc(-c3ccc4c(c3)OCO4)c2)cc(OC)c1OC
428. COc1cc(-c2cc(N3CCN(C)CC3)ccc2O)ccc1OCCO
429. O=C1Nc2c(Cl)cccc2C(=O)Nc2ccc(Cl)c(Cl)c21
430. Cc1ccc(NC(=O)c2ccc(C(C)C)cc2C(=O)Nc2ccccc2C(C)C)cc1
431. COc1ccc(C(=O)Nc2cccc(C(=O)Nc3cccc(C(=O)O)c3)c2)cc1
432. COC(=O)c1cc(Cn2c(=O)n(Cc3ccccc3)c3ccccc3c2=O)ccn1
433. CC(C)[C@H](O)[C@H](C)NC(=O)[C@H](CC(C)C)NC(=O)[C@H](CC(C)C)NC(=O)[C@H](Cc1ccccc1)NC(=O)OC(C)(C)C
434. C[C@]12CC[C@@H]3c4ccc(O)cc4CC[C@H]3[C@@H]1CCC2=O
435. COc1ccc(C[C@H]2CC[C@H](NC(=O)c3ccc(C(C)(C)C)cc3)CC2)cc1
436. COc1cc(N2CCN(C3CCCCC3)CC2)c2ncn(C3CCCCC3)c2c1
437. CCCN(CCC)C(=O)C(C)(C)NC(=O)OCc1ccccc1
438. CC(=O)N1CCC(NC(=O)C2(NC(=O)[C@@H](Cc3ccccc3)NC(=O)c3ccccc3Cl)CC2)CC1
439. CCCNC(=O)c1ccc(C(=O)Nc2cccc([N+](=O)[O-])c2)cc1
440. COc1c(N)ncnc1N1CCC(c2nc(-c3ccc(F)c(C(F)(F)F)c3)cn2CCN2CCCC2)CC1
441. CN(C)C(=O)C1CCN(Cc2ccc(OC(=O)Nc3ccccc3)cc2)CC1
442. Cc1cc(N2CCOCC2)ccc1-c1nc2ccccc2[nH]1
443. CN(C)C(=O)c1cccc(-c2ccc(N3CCN(c4ncnc5ccccc45)CC3)nc2)c1
444. COc1cc(-c2ccc(NC(=O)C3CC3)cc2)ccc1OCCN1CCOCC1
445. CC(=O)N[C@@H](Cc1ccccc1)C(=O)N[C@H](Cc1ccccc1)CP(=O)(O)O
446. COc1ccc(Cc2c(O)c(O)c(O)c(O)c2O)cc1
447. CC(=O)N[C@@H](Cc1ccccc1)C(=O)N[C@H](Cc1ccccc1)C(=O)O
448. CC(C)[C@@H](O)[C@H](Cc1ccc(Cl)c(Cl)c1)c1ccc(C2CC2)cc1
449. CCN(CC)CCNC(=O)c1c(C)[nH]c(/C=C2\C(=O)Nc3ccc(F)cc32)c1C
450. N[C@@H]1C[C@H]2O[C@@](C)([C@@H]1OC)n1c3ccccc3c3c4c(c5c6ccccc6n2c5c31)C(=O)NC4
451. COc1ccc(S(=O)(=O)N2CCN(Cc3ccccc3)CC2)cc1
452. COc1cc(C(=O)N2CCC(C(=O)O)CC2)ccc1-c1ccc(OC)cc1
453. CN(C)CCCNC(=O)c1cc(-c2ccc(Oc3ccccc3)cc2)[nH]n1
454. CN1CCN(c2ccc(-c3c[nH]c4ccccc34)cc2)CC1
455. CNC(=O)c1c(C)c2ccccc2n1CC(=O)NC1CCN(CCCc2c[nH]c3ccccc23)CC1

Supplementary Table S1. SMILES representation for generated compounds.

### Graph representation of all the generated molecules

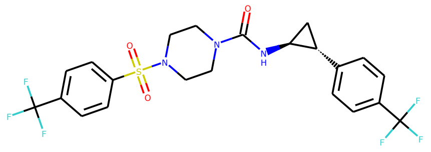

Generated molecule 1

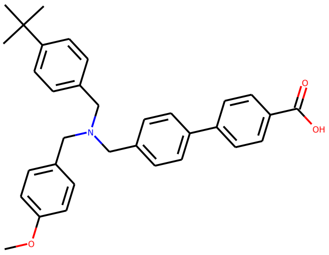

Generated molecule 2

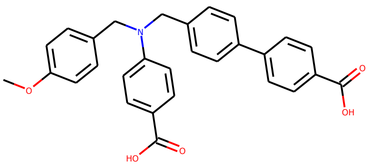

Generated molecule 3

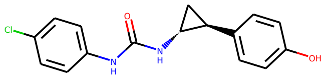

Generated molecule 4

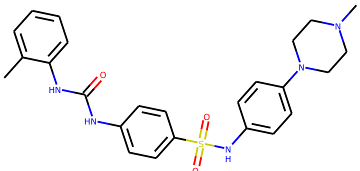

Generated molecule 5

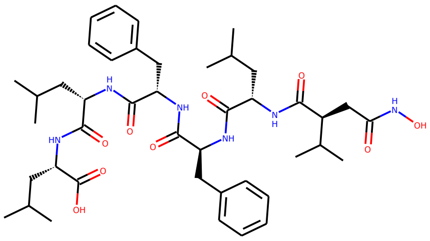

Generated molecule 6

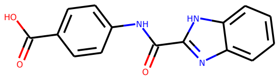

Generated molecule 7

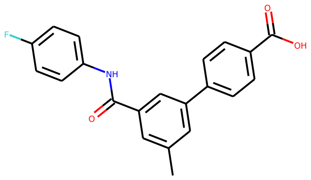

Generated molecule 8

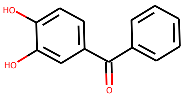

Generated molecule 9

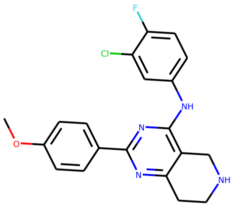

Generated molecule 10

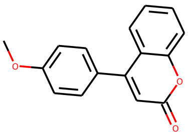

Generated molecule 11

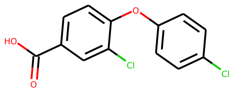

Generated molecule 12

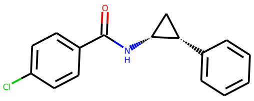

Generated molecule 13

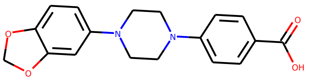

Generated molecule 14

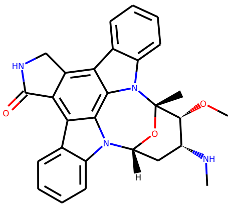

Generated molecule 15

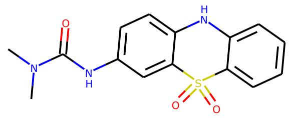

Generated molecule 16

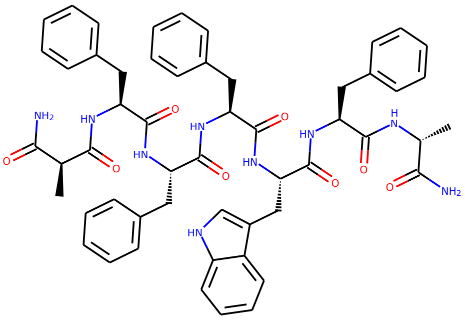

Generated molecule 17

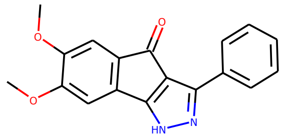

Generated molecule 18

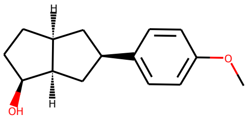

Generated molecule 19

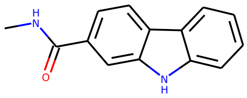

Generated molecule 20

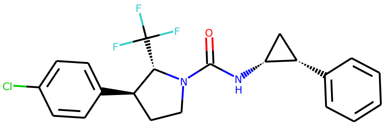

Generated molecule 21

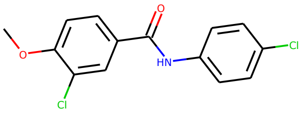

Generated molecule 22

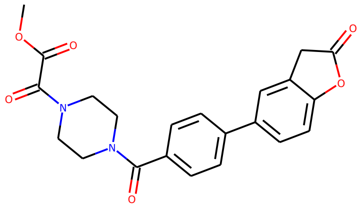

Generated molecule 23

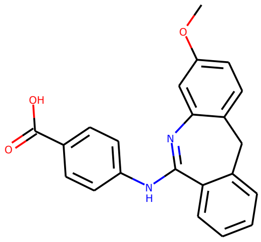

Generated molecule 24

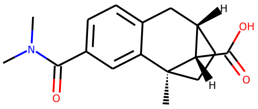

Generated molecule 25

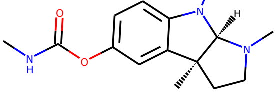

Generated molecule 26

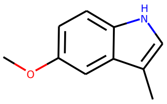

Generated molecule 27

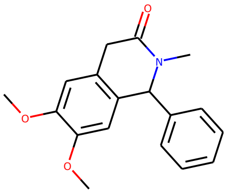

Generated molecule 28

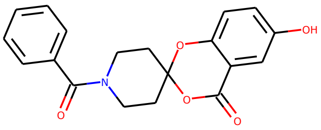

Generated molecule 29

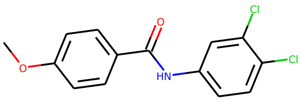

Generated molecule 30

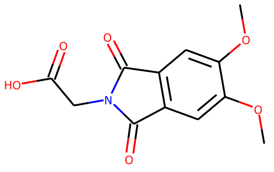


Generated molecule 31

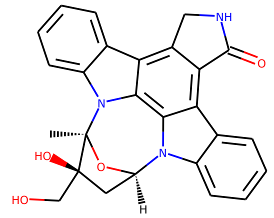


Generated molecule 32

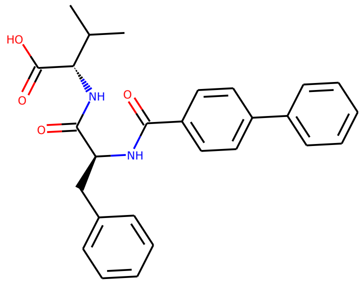


Generated molecule 33

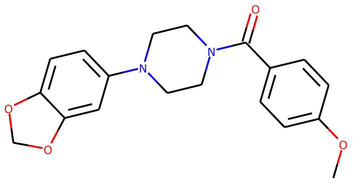


Generated molecule 34

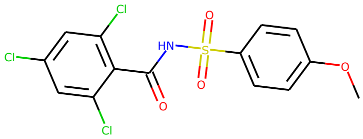


Generated molecule 35

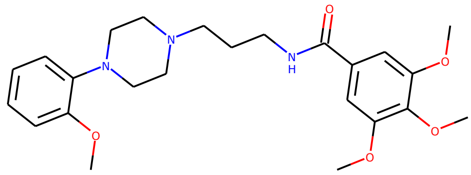


Generated molecule 36

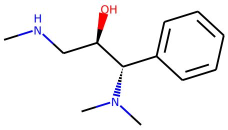


Generated molecule 37

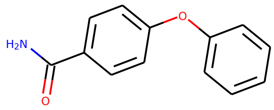


Generated molecule 38

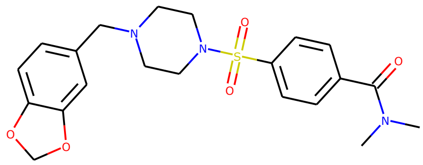


Generated molecule 39

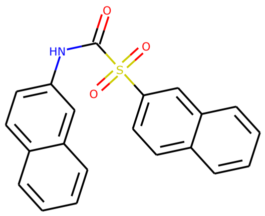


Generated molecule 40

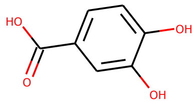


Generated molecule 41

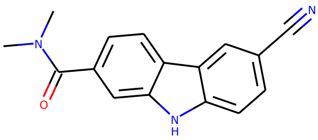


Generated molecule 42

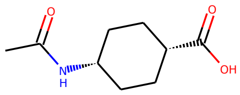


Generated molecule 43

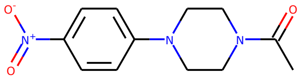


Generated molecule 44

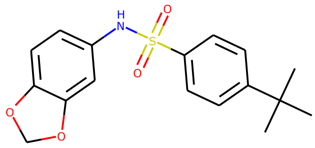


Generated molecule 45

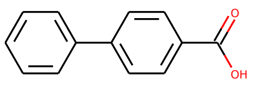

Generated molecule 46

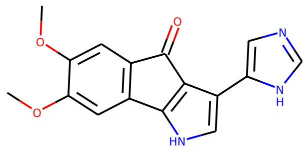

Generated molecule 47

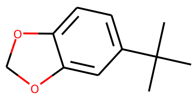

Generated molecule 48

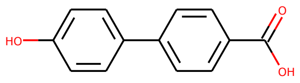

Generated molecule 49

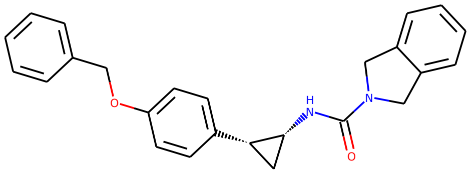

Generated molecule 50

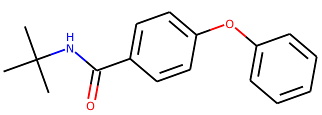

Generated molecule 51

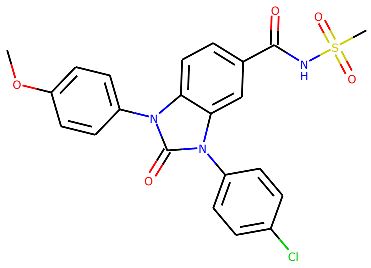

Generated molecule 52

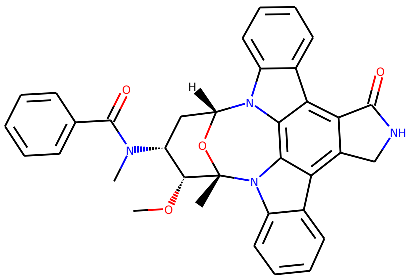

Generated molecule 53

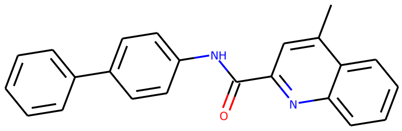

Generated molecule 54

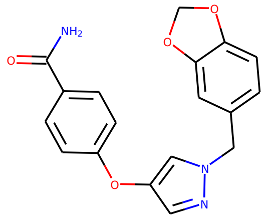

Generated molecule 55

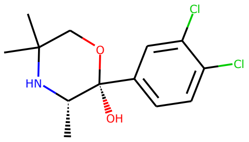

Generated molecule 56

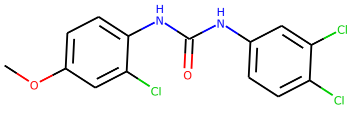

Generated molecule 57

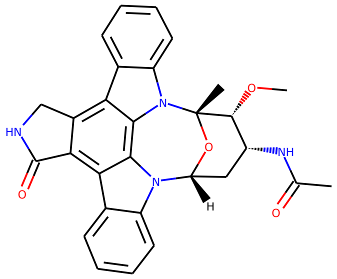

Generated molecule 58

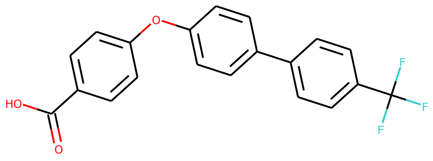

Generated molecule 59

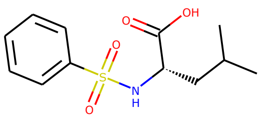

Generated molecule 60

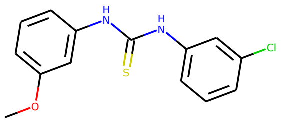

Generated molecule 61

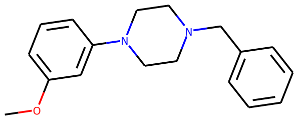

Generated molecule 62

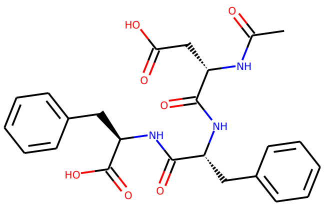

Generated molecule 63

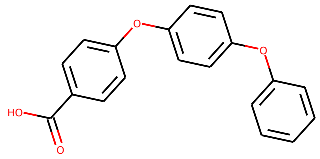

Generated molecule 64

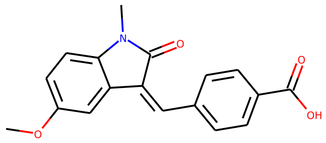

Generated molecule 65

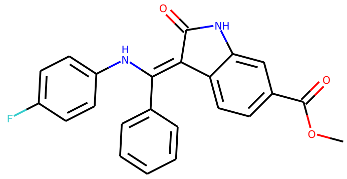

Generated molecule 66

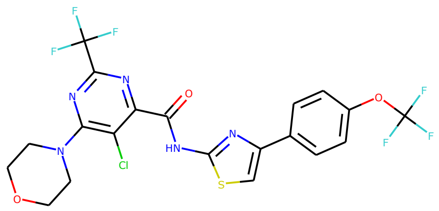

Generated molecule 67

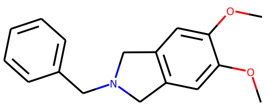

Generated molecule 68

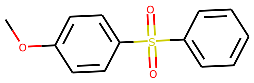

Generated molecule 69

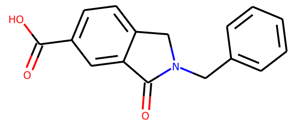

Generated molecule 70

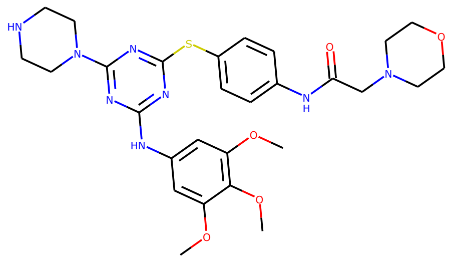

Generated molecule 71

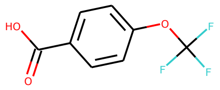

Generated molecule 72

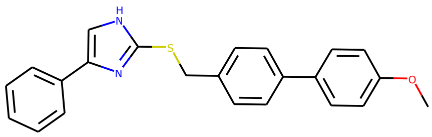

Generated molecule 73

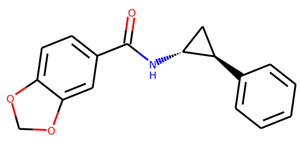

Generated molecule 74

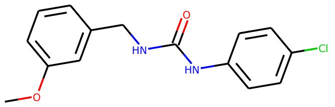

Generated molecule 75

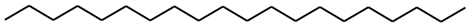

Generated molecule 76

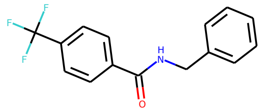

Generated molecule 77

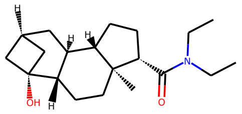

Generated molecule 78

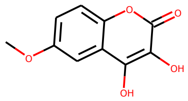

Generated molecule 79

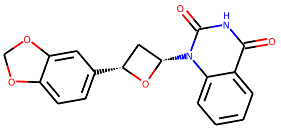

Generated molecule 80

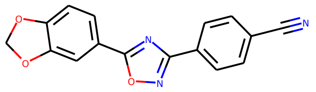

Generated molecule 81

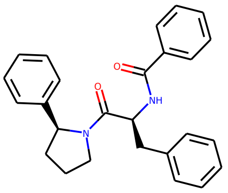

Generated molecule 82

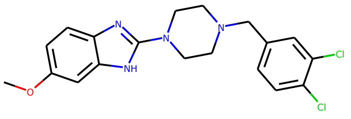

Generated molecule 83

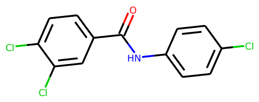

Generated molecule 84

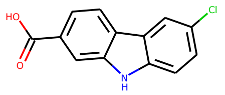

Generated molecule 85

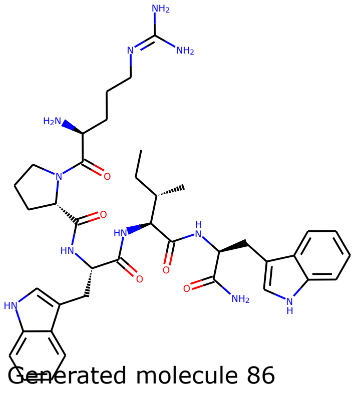

Generated molecule 86

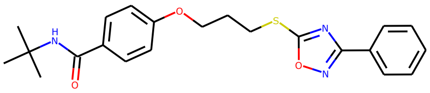

Generated molecule 87

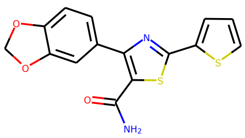

Generated molecule 88

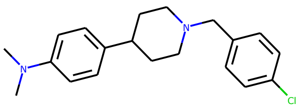

Generated molecule 89

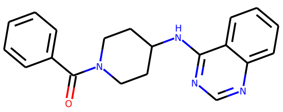

Generated molecule 90

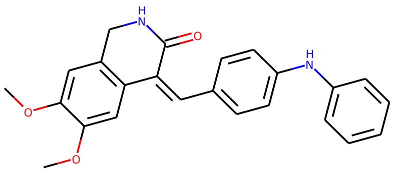

Generated molecule 91

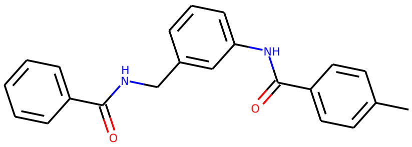

Generated molecule 92

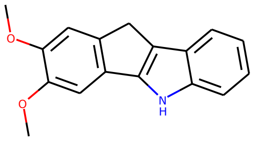

Generated molecule 93

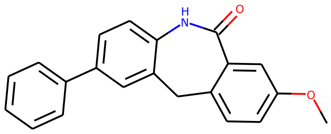

Generated molecule 94

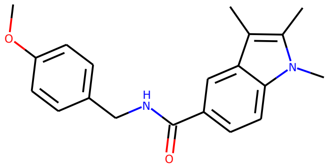

Generated molecule 95

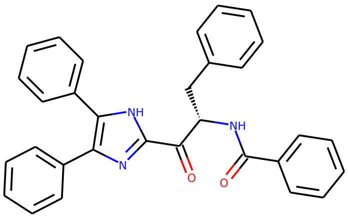

Generated molecule 96

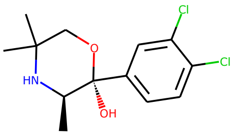

Generated molecule 97

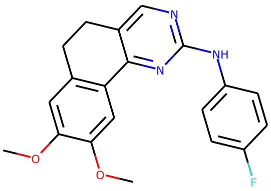

Generated molecule 98

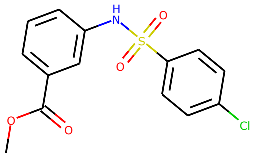

Generated molecule 99

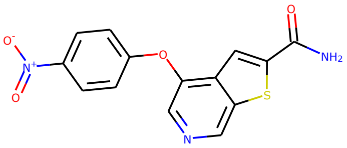

Generated molecule 100

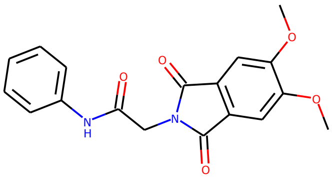

Generated molecule 101

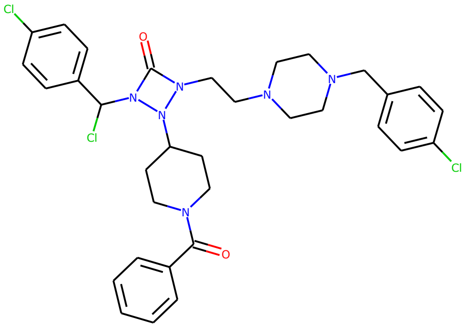

Generated molecule 102

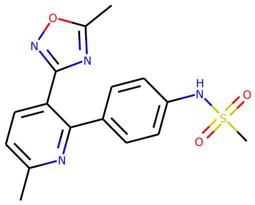

Generated molecule 103

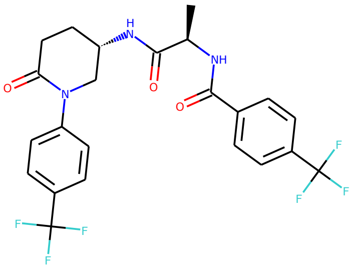

Generated molecule 104

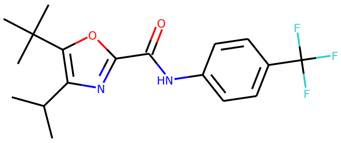

Generated molecule 105

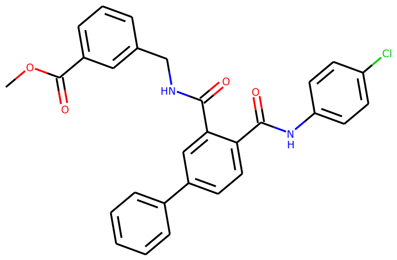


Generated molecule 106

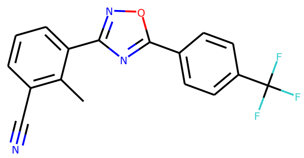


Generated molecule 107

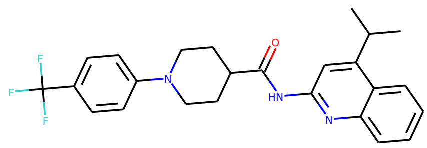


Generated molecule 108

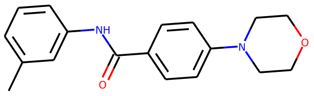


Generated molecule 109

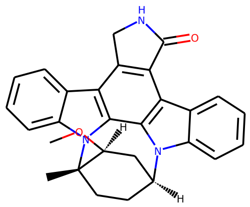


Generated molecule 110

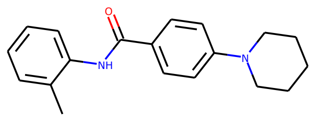


Generated molecule 111

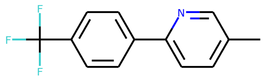


Generated molecule 112

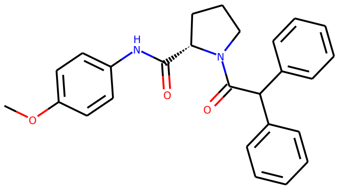


Generated molecule 113

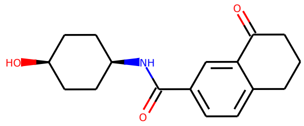


Generated molecule 114

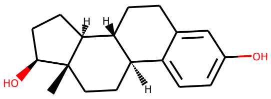


Generated molecule 115

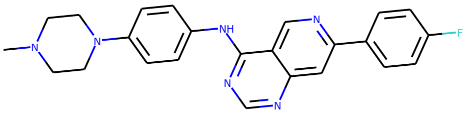


Generated molecule 116

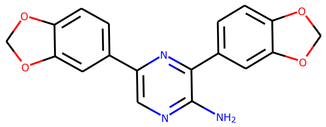


Generated molecule 117

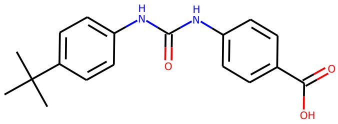


Generated molecule 118

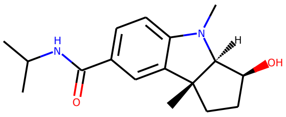


Generated molecule 119

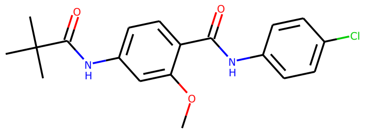


Generated molecule 120

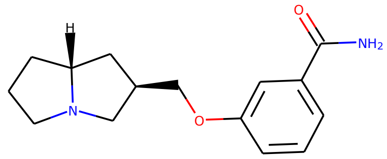

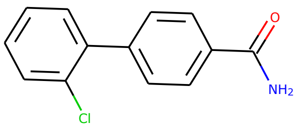

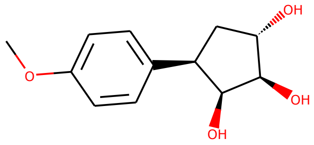

Generated molecule 121

Generated molecule 122

Generated molecule 123

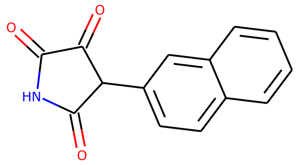

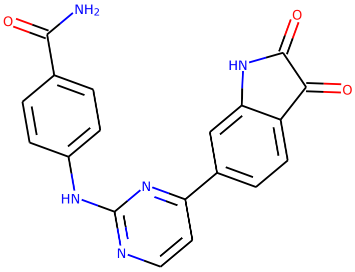

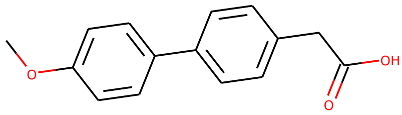

Generated molecule 124

Generated molecule 125

Generated molecule 126

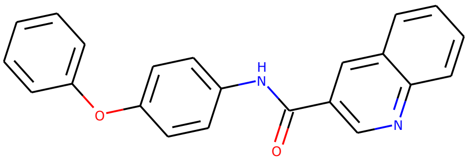

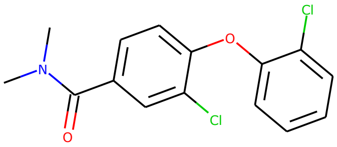

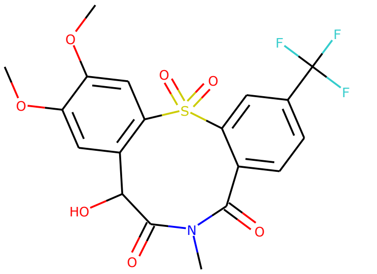

Generated molecule 127

Generated molecule 128

Generated molecule 129

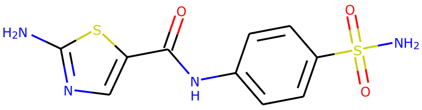

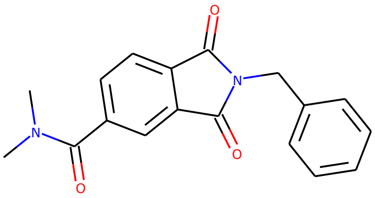

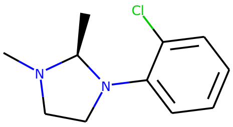

Generated molecule 130

Generated molecule 131

Generated molecule 132

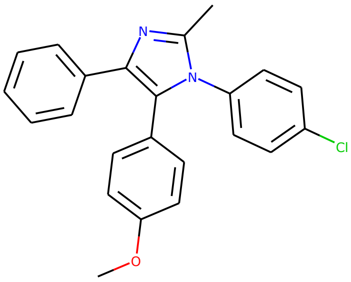

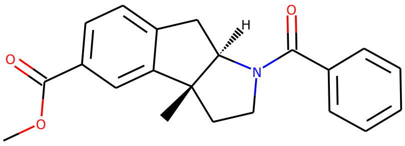

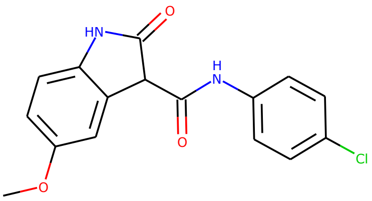

Generated molecule 133

Generated molecule 134

Generated molecule 135

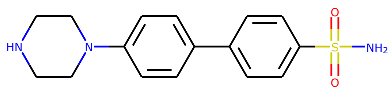


Generated molecule 136

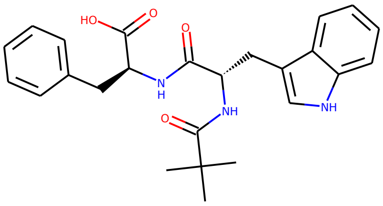

Generated molecule 137

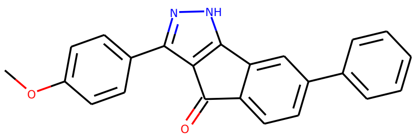

Generated molecule 138

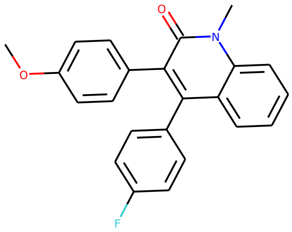

Generated molecule 139

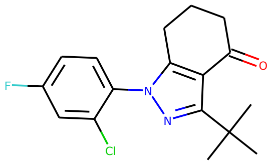

Generated molecule 140

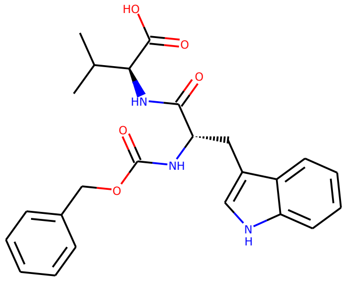

Generated molecule 141

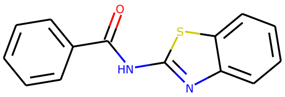

Generated molecule 142

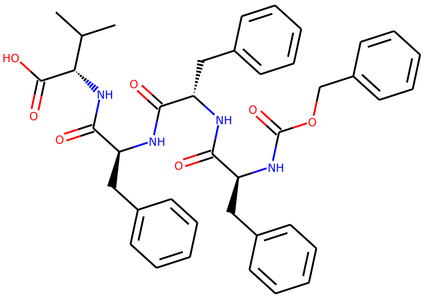

Generated molecule 143

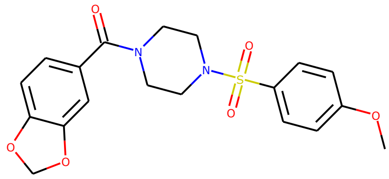

Generated molecule 144

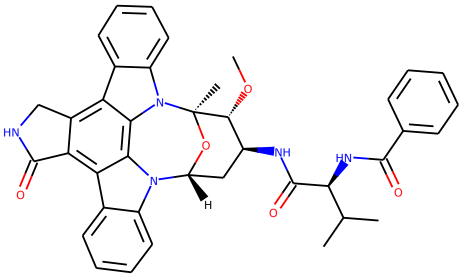

Generated molecule 145

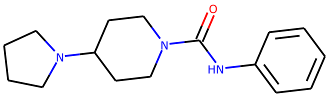

Generated molecule 146

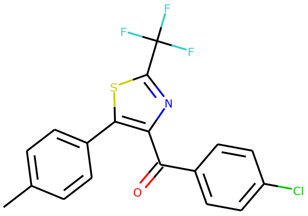

Generated molecule 147

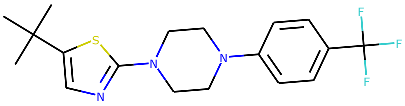

Generated molecule 148

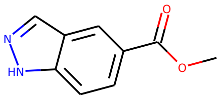

Generated molecule 149

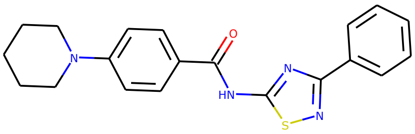

Generated molecule 150

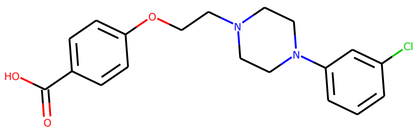

Generated molecule 151

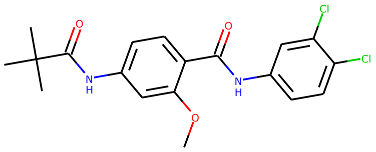

Generated molecule 152

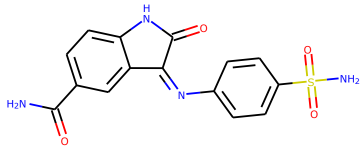

Generated molecule 153

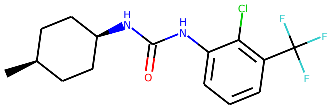

Generated molecule 154

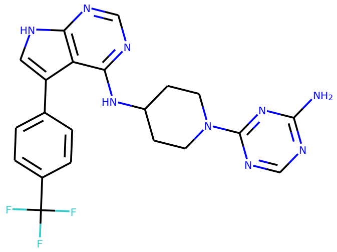

Generated molecule 155

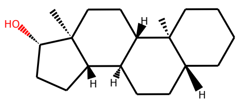

Generated molecule 156

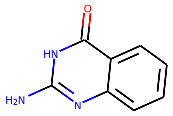

Generated molecule 157

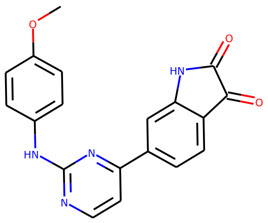

Generated molecule 158

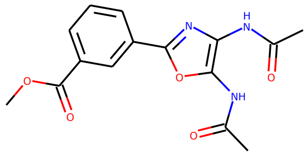

Generated molecule 159

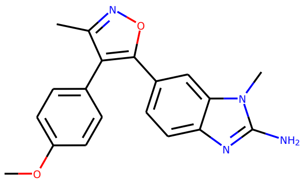

Generated molecule 160

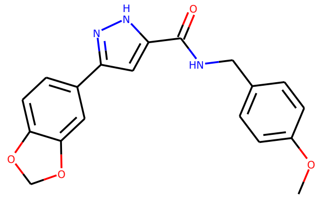

Generated molecule 161

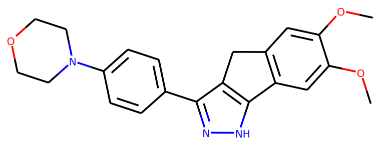

Generated molecule 162

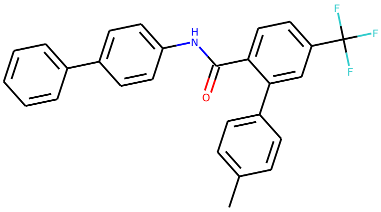

Generated molecule 163

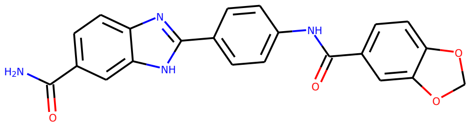

Generated molecule 164

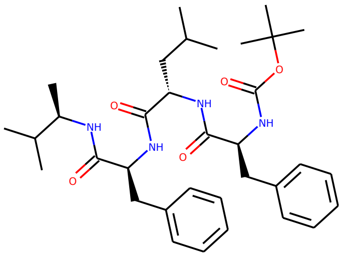

Generated molecule 165

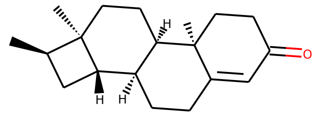

Generated molecule 166

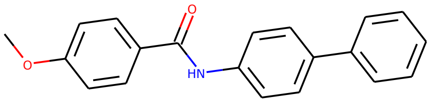

Generated molecule 167

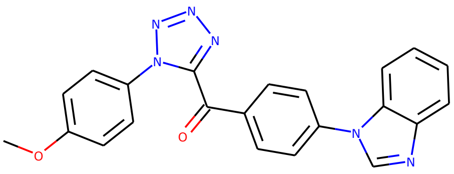

Generated molecule 168

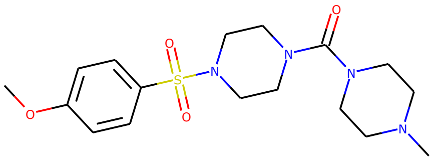

Generated molecule 169

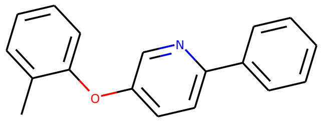

Generated molecule 170

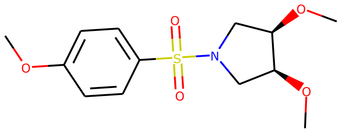

Generated molecule 171

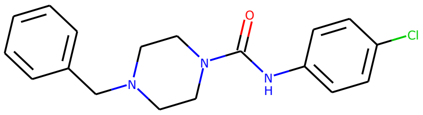

Generated molecule 172

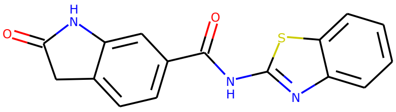

Generated molecule 173

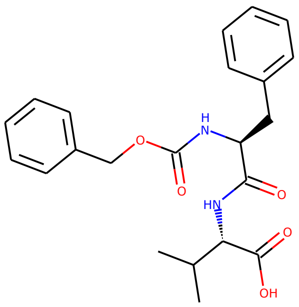

Generated molecule 174

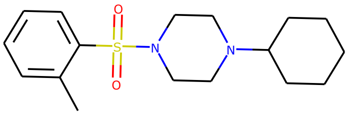

Generated molecule 175

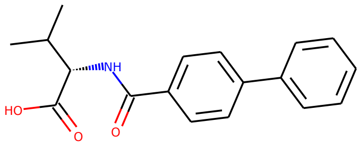

Generated molecule 176

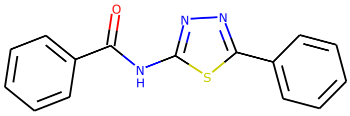

Generated molecule 177

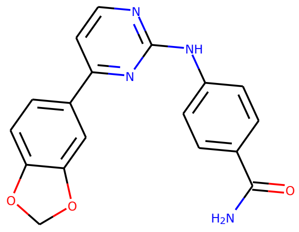

Generated molecule 178

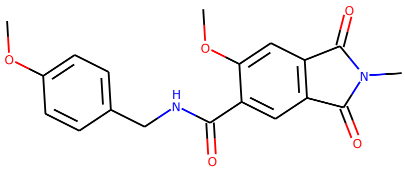

Generated molecule 179

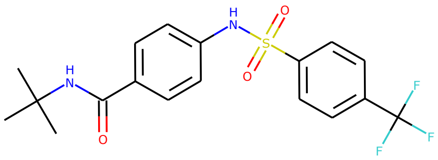

Generated molecule 180

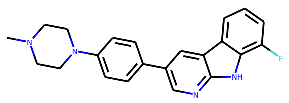

Generated molecule 181

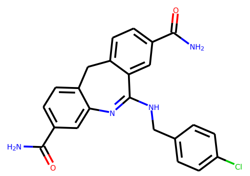

Generated molecule 182

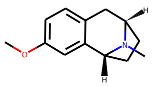

Generated molecule 183

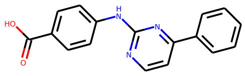

Generated molecule 184

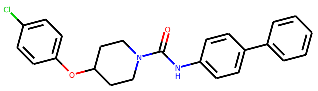

Generated molecule 185

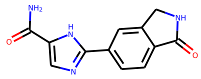

Generated molecule 186

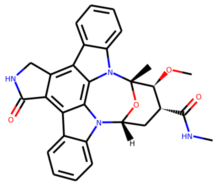

Generated molecule 187

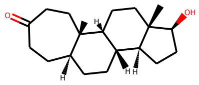

Generated molecule 188

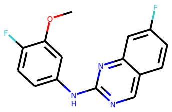

Generated molecule 189

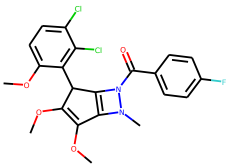

Generated molecule 190

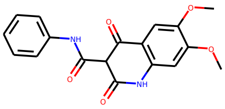

Generated molecule 191

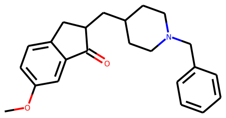

Generated molecule 192

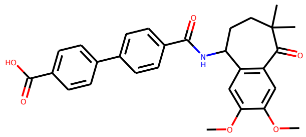

Generated molecule 193

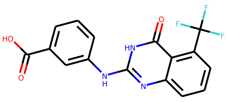

Generated molecule 194

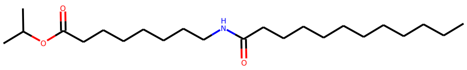

Generated molecule 195

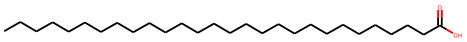

Generated molecule 196

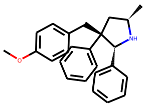

Generated molecule 197

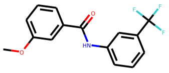

Generated molecule 198

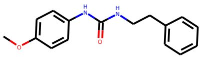

Generated molecule 199

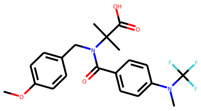

Generated molecule 200

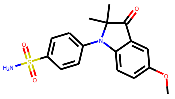

Generated molecule 201

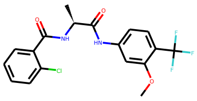

Generated molecule 202

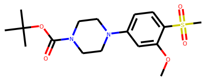

Generated molecule 203

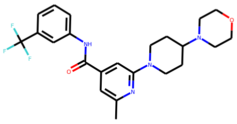

Generated molecule 204

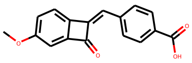

Generated molecule 205

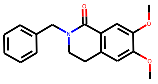

Generated molecule 206

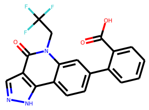

Generated molecule 207

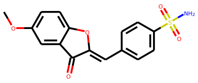

Generated molecule 208

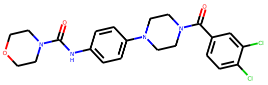

Generated molecule 209

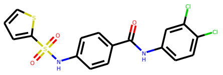

Generated molecule 210

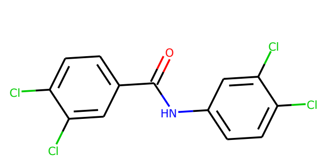

Generated molecule 211

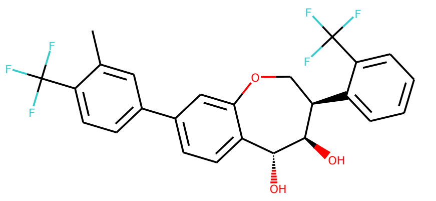

Generated molecule 212

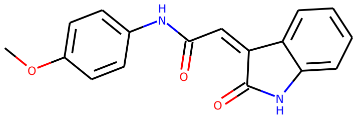

Generated molecule 213

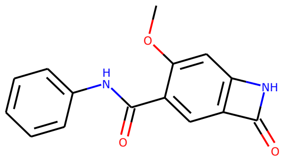

Generated molecule 214

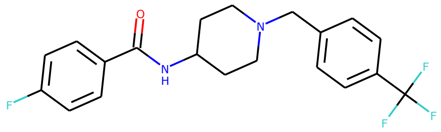

Generated molecule 215

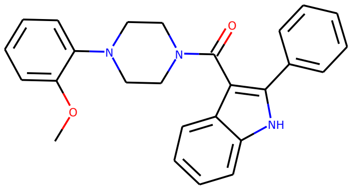

Generated molecule 216

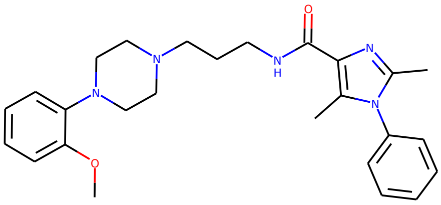

Generated molecule 217

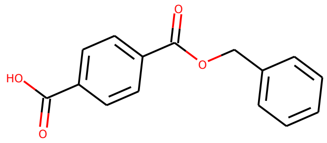

Generated molecule 218

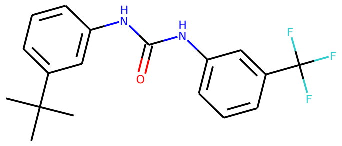

Generated molecule 219

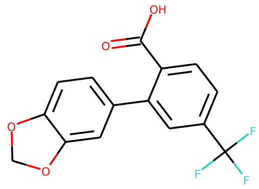

Generated molecule 220

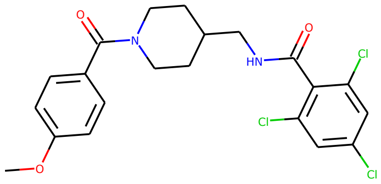

Generated molecule 221

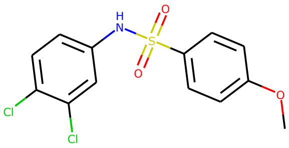

Generated molecule 222

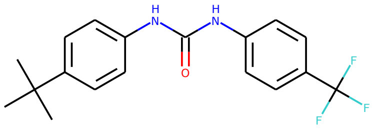

Generated molecule 223

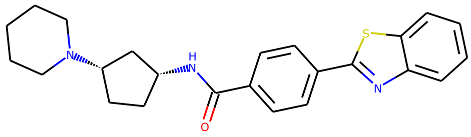

Generated molecule 224

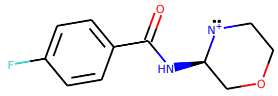

Generated molecule 225

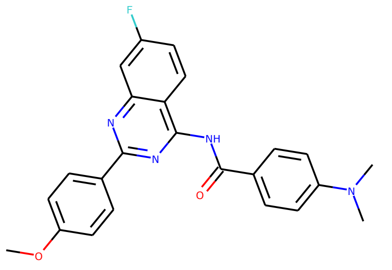

Generated molecule 226

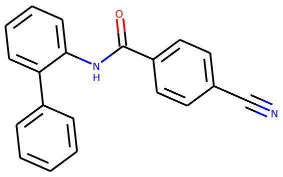

Generated molecule 227

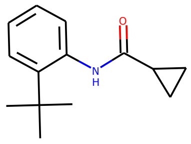

Generated molecule 228

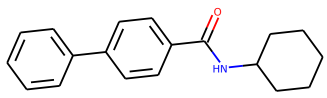

Generated molecule 229

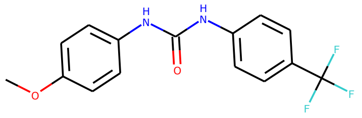

Generated molecule 230

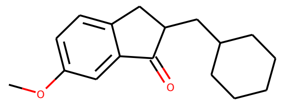

Generated molecule 231

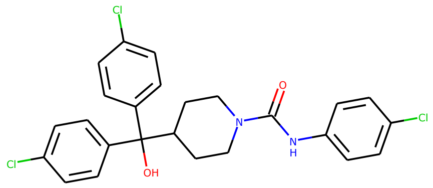

Generated molecule 232

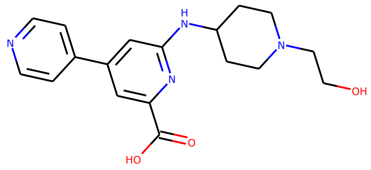

Generated molecule 233

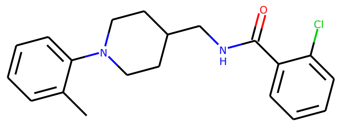

Generated molecule 234

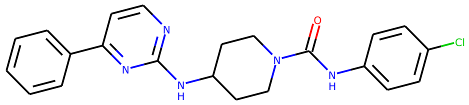

Generated molecule 235

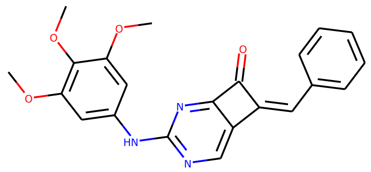

Generated molecule 236

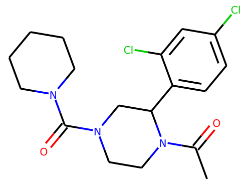

Generated molecule 237

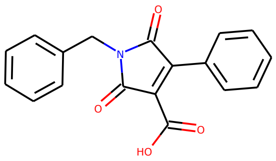

Generated molecule 238

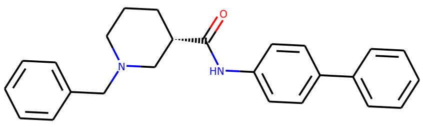

Generated molecule 239

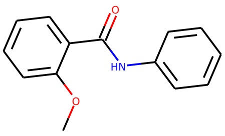

Generated molecule 240

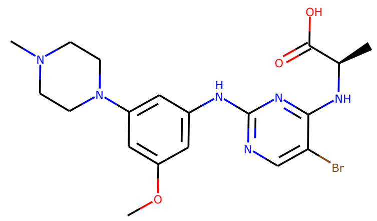

Generated molecule 241

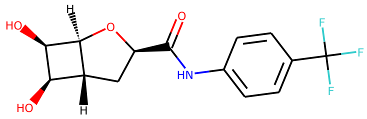

Generated molecule 242

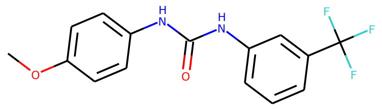

Generated molecule 243

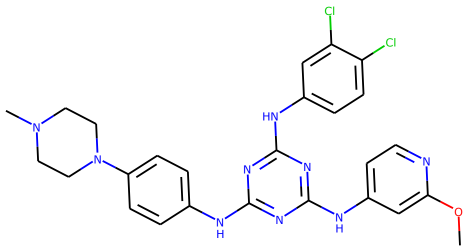

Generated molecule 244

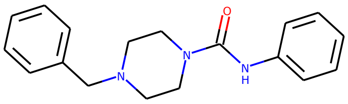

Generated molecule 245

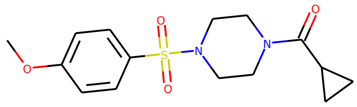

Generated molecule 246

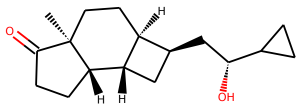

Generated molecule 247

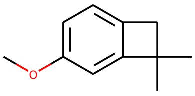

Generated molecule 248

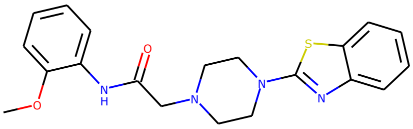

Generated molecule 249

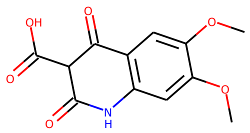

Generated molecule 250

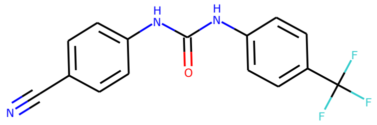

Generated molecule 251

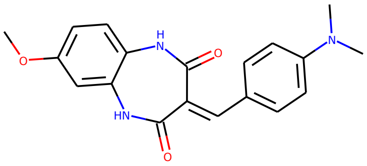

Generated molecule 252

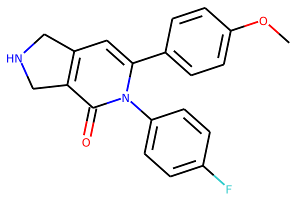

Generated molecule 253

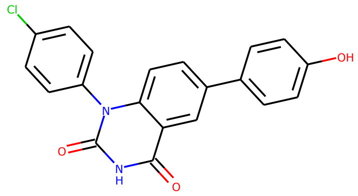

Generated molecule 254

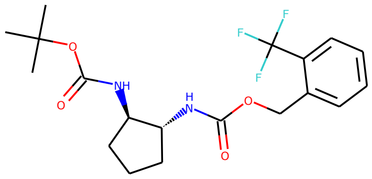

Generated molecule 255

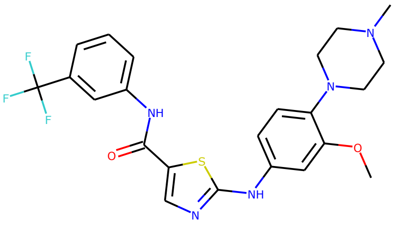

Generated molecule 256

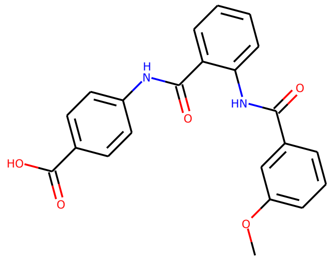

Generated molecule 257

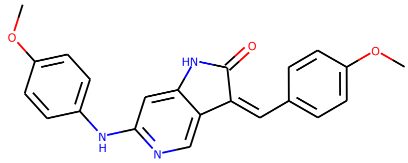

Generated molecule 258

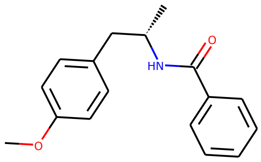

Generated molecule 259

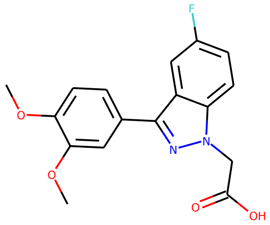

Generated molecule 260

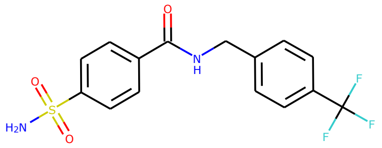

Generated molecule 261

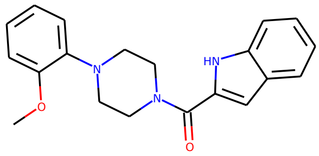

Generated molecule 262

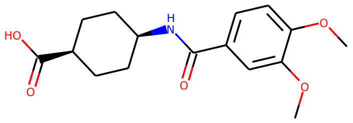

Generated molecule 263

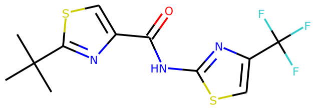

Generated molecule 264

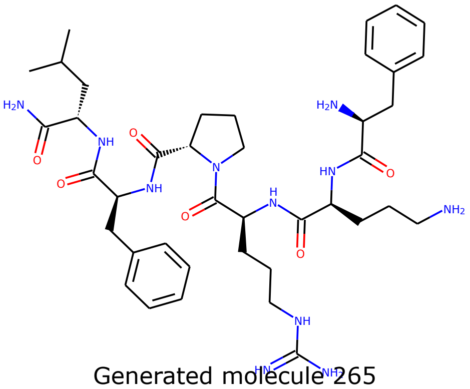

Generated molecule 265

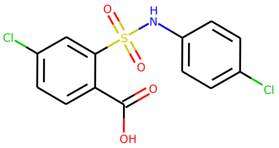

Generated molecule 266

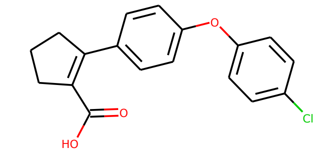

Generated molecule 267

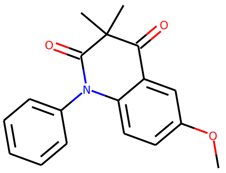

Generated molecule 268

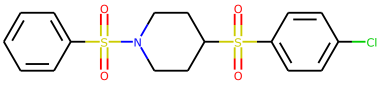

Generated molecule 269

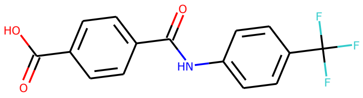

Generated molecule 270

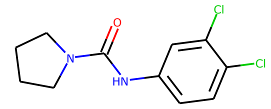

Generated molecule 271

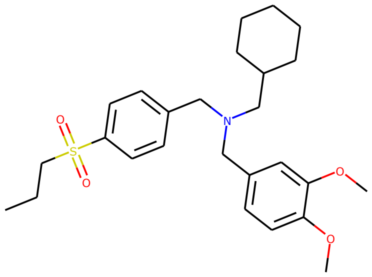

Generated molecule 272

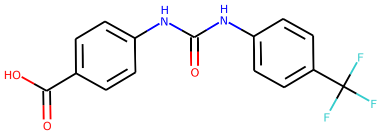

Generated molecule 273

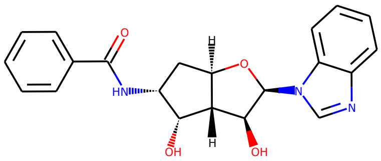

Generated molecule 274

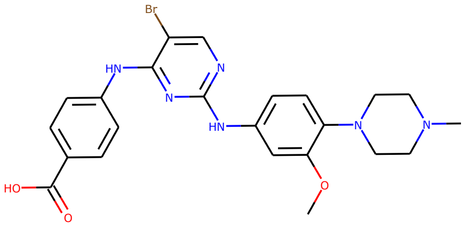

Generated molecule 275

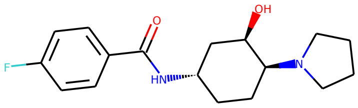

Generated molecule 276

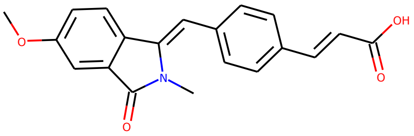

Generated molecule 277

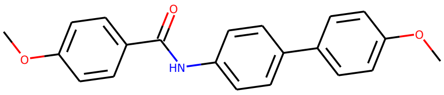

Generated molecule 278

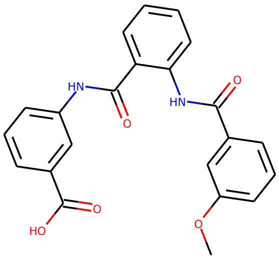

Generated molecule 279

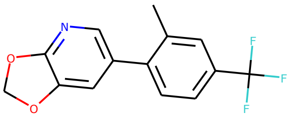

Generated molecule 280

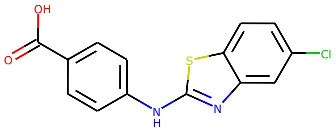

Generated molecule 281

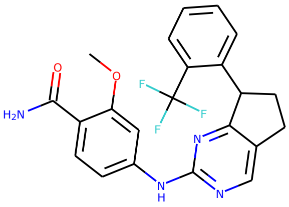

Generated molecule 282

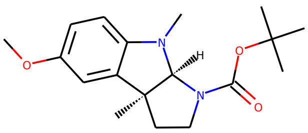

Generated molecule 283

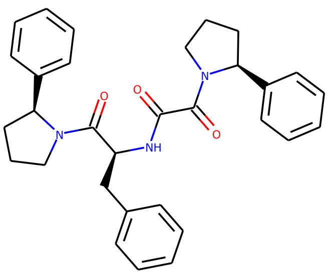

Generated molecule 284

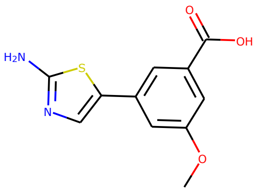

Generated molecule 285

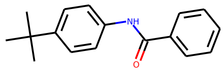

Generated molecule 286

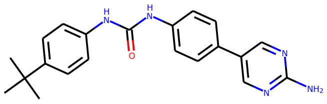

Generated molecule 287

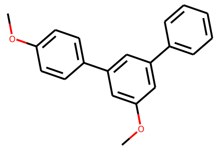

Generated molecule 288

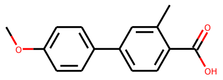

Generated molecule 289

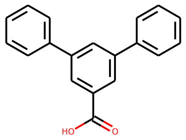

Generated molecule 290

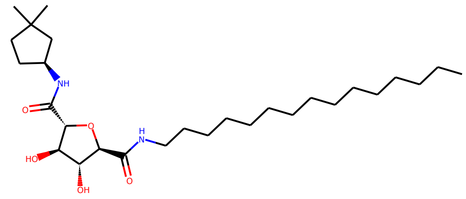

Generated molecule 291

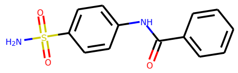

Generated molecule 292

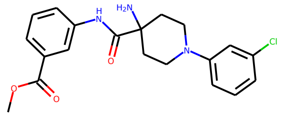

Generated molecule 293

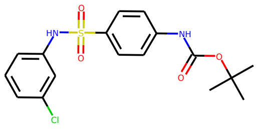

Generated molecule 294

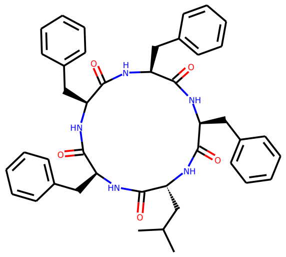

Generated molecule 295

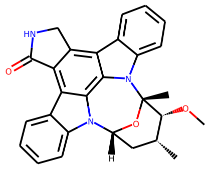

Generated molecule 296

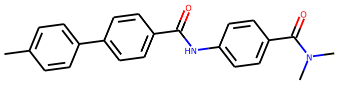

Generated molecule 297

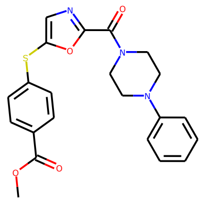

Generated molecule 298

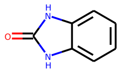

Generated molecule 299

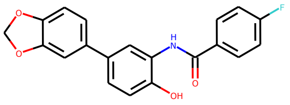

Generated molecule 300

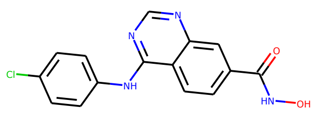

Generated molecule 301

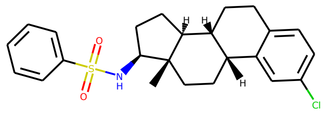

Generated molecule 302

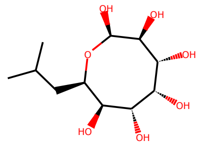

Generated molecule 303

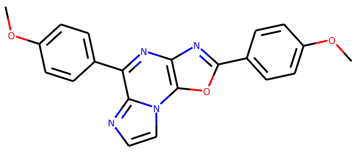

Generated molecule 304

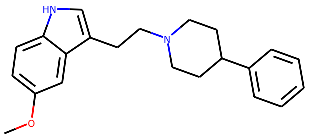

Generated molecule 305

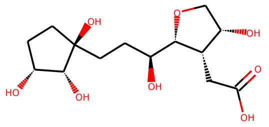

Generated molecule 306

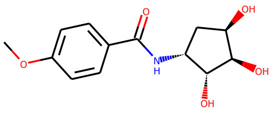

Generated molecule 307

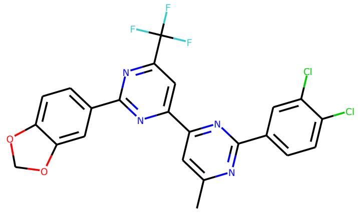

Generated molecule 308

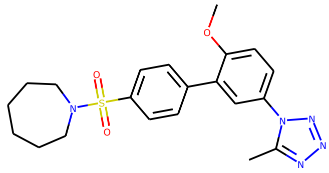

Generated molecule 309

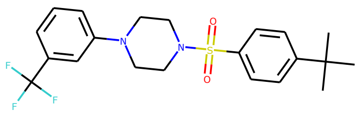

Generated molecule 310

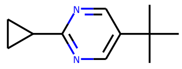

Generated molecule 311

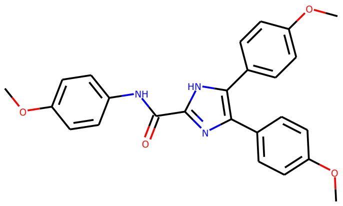

Generated molecule 312

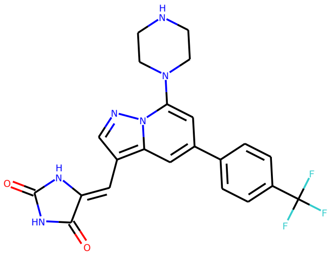

Generated molecule 313

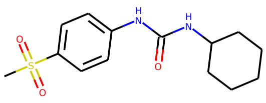

Generated molecule 314

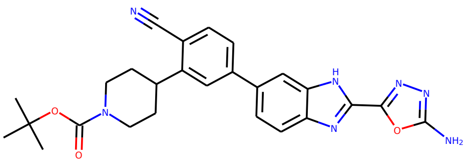

Generated molecule 315

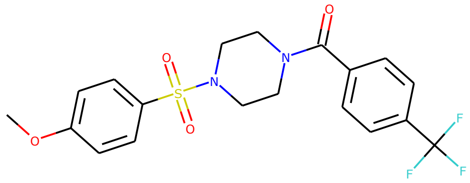

Generated molecule 316

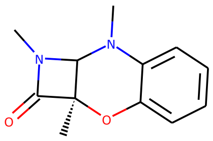

Generated molecule 317

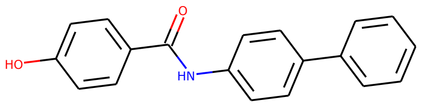

Generated molecule 318

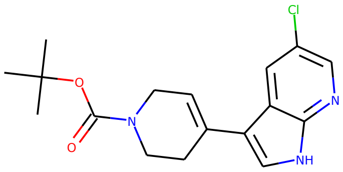

Generated molecule 319

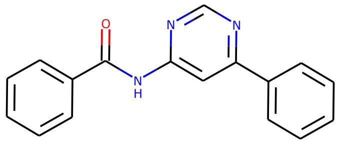

Generated molecule 320

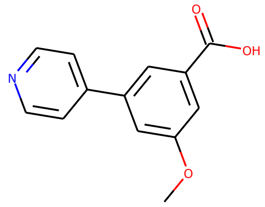

Generated molecule 321

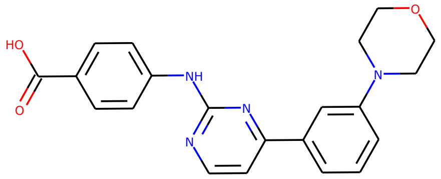

Generated molecule 322

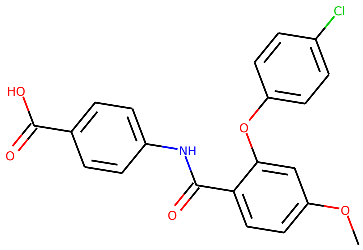

Generated molecule 323

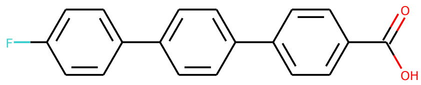

Generated molecule 324

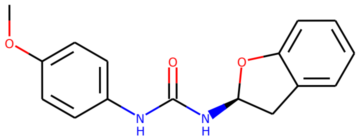

Generated molecule 325

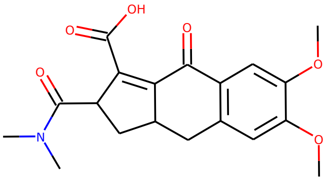

Generated molecule 326

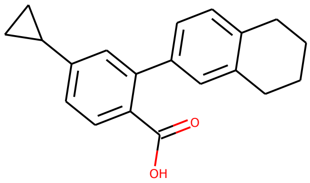

Generated molecule 327

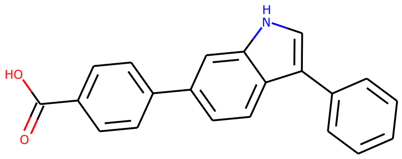

Generated molecule 328

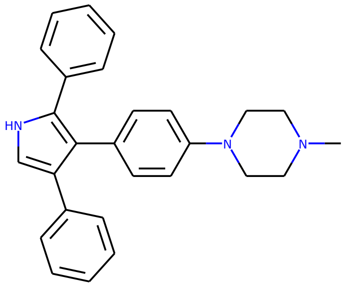

Generated molecule 329

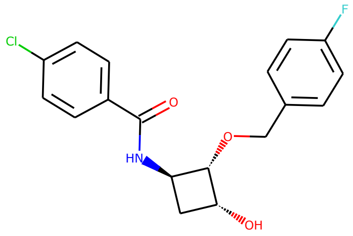

Generated molecule 330

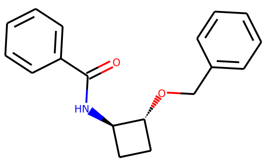

Generated molecule 331

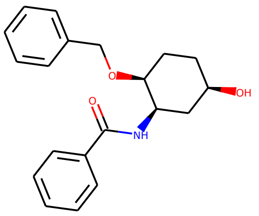

Generated molecule 332

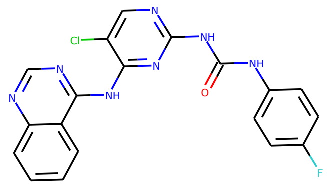

Generated molecule 333

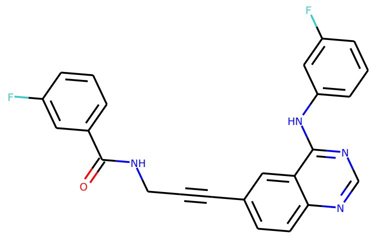

Generated molecule 334

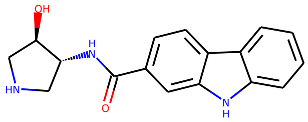

Generated molecule 335

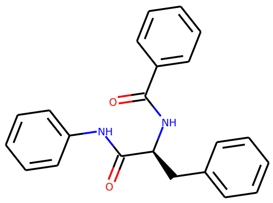

Generated molecule 336

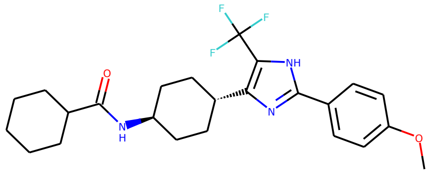

Generated molecule 337

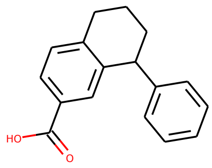

Generated molecule 338

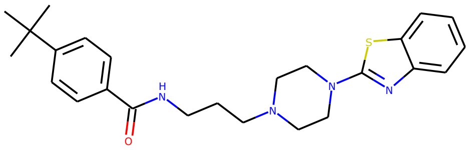

Generated molecule 339

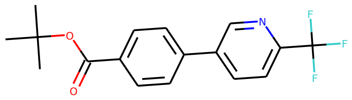

Generated molecule 340

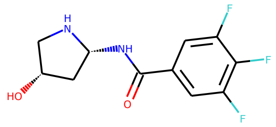

Generated molecule 341

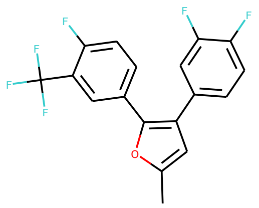

Generated molecule 342

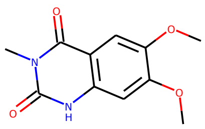

Generated molecule 343

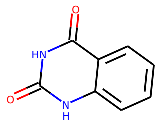

Generated molecule 344

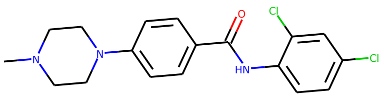

Generated molecule 345

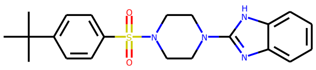

Generated molecule 346

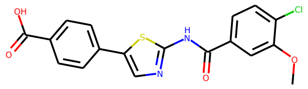

Generated molecule 347

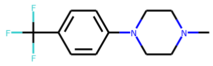

Generated molecule 348

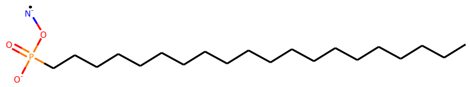

Generated molecule 349

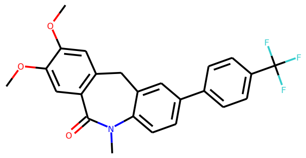

Generated molecule 350

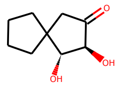

Generated molecule 351

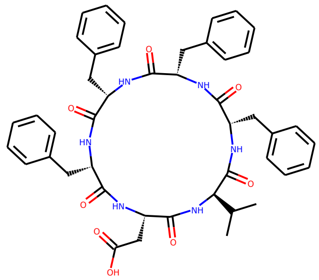

Generated molecule 352

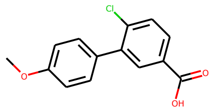

Generated molecule 353

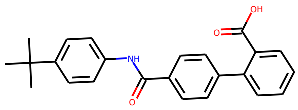

Generated molecule 354

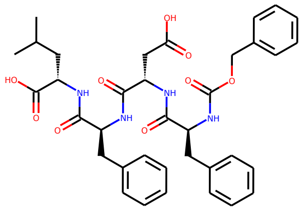

Generated molecule 355

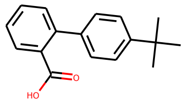

Generated molecule 356

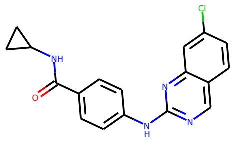

Generated molecule 357

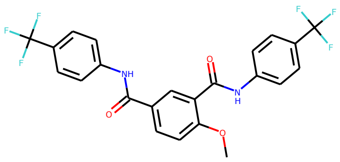

Generated molecule 358

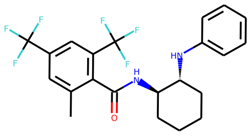

Generated molecule 359

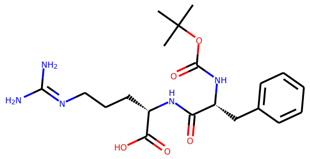

Generated molecule 360

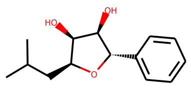

Generated molecule 361

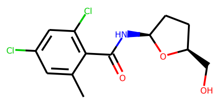

Generated molecule 362

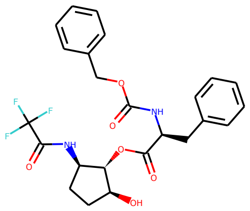

Generated molecule 363

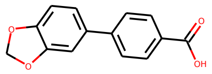

Generated molecule 364

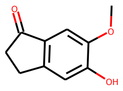

Generated molecule 365

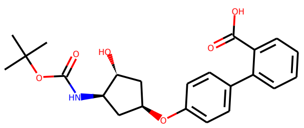

Generated molecule 366

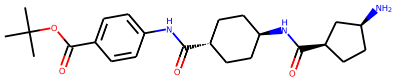

Generated molecule 367

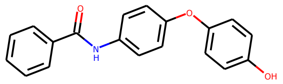

Generated molecule 368

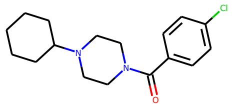

Generated molecule 369

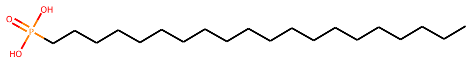

Generated molecule 370

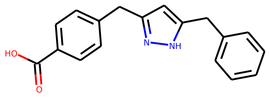

Generated molecule 371

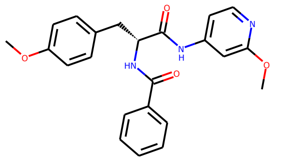

Generated molecule 372

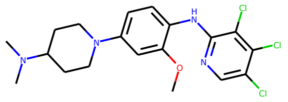

Generated molecule 373

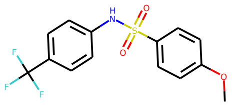

Generated molecule 374

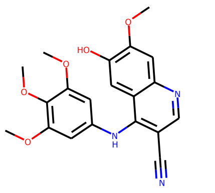

Generated molecule 375

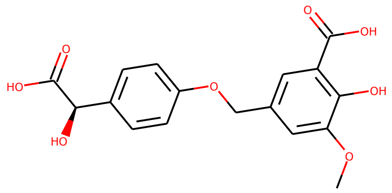

Generated molecule 376

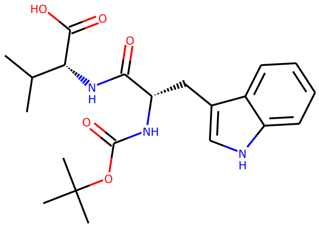

Generated molecule 377

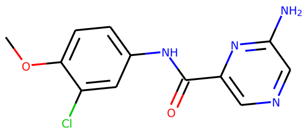

Generated molecule 378

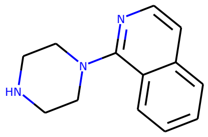

Generated molecule 379

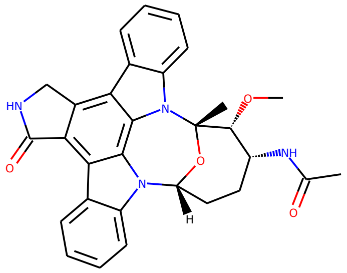

Generated molecule 380

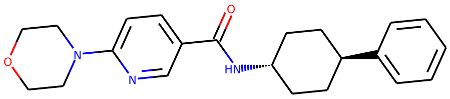

Generated molecule 381

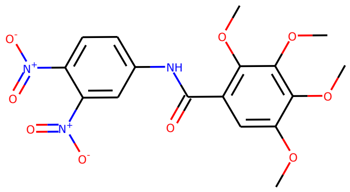

Generated molecule 382

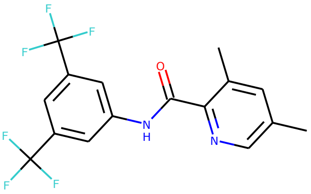

Generated molecule 383

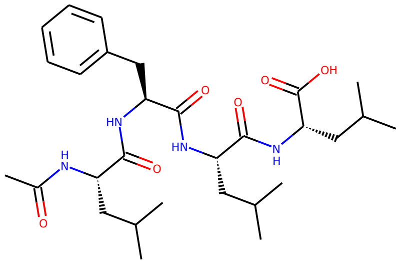

Generated molecule 384

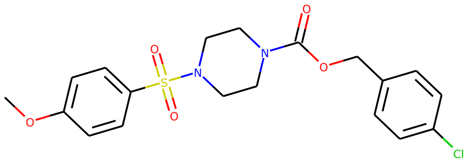

Generated molecule 385

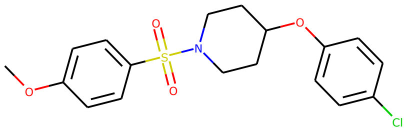

Generated molecule 386

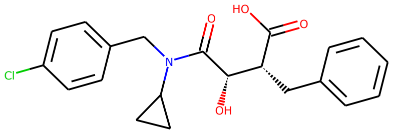

Generated molecule 387

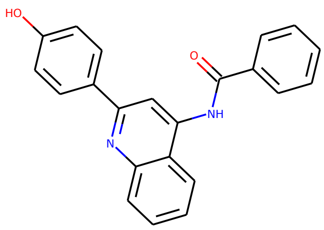

Generated molecule 388

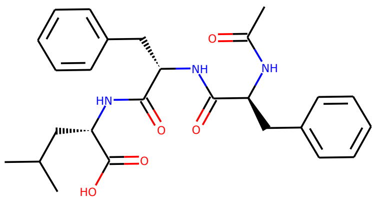

Generated molecule 389

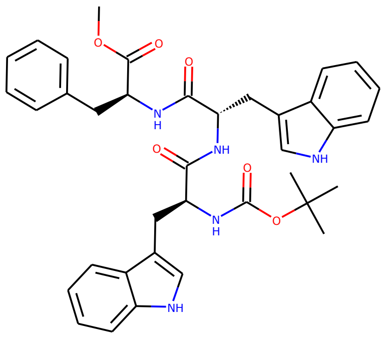

Generated molecule 390

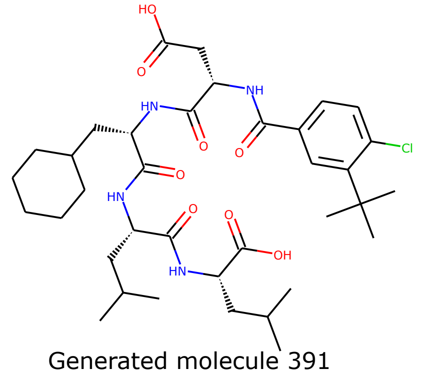

Generated molecule 391

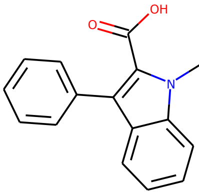

Generated molecule 392

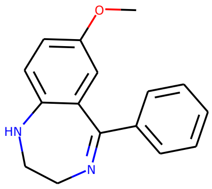

Generated molecule 393

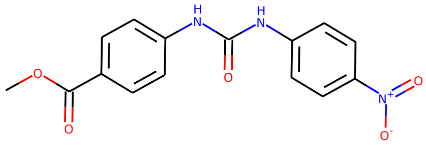

Generated molecule 394

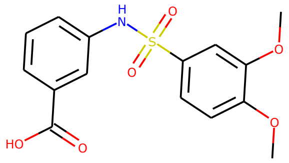

Generated molecule 395

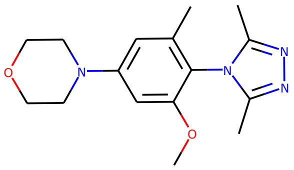

Generated molecule 396

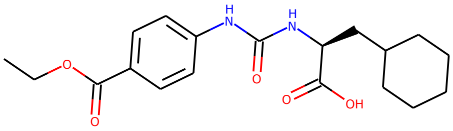

Generated molecule 397

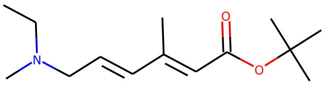

Generated molecule 398

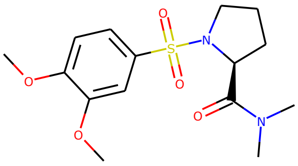

Generated molecule 399

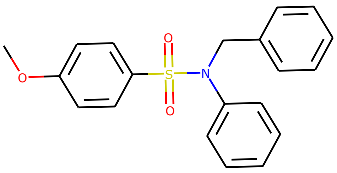

Generated molecule 400

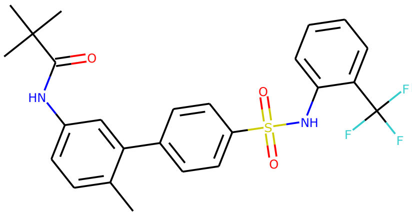

Generated molecule 401

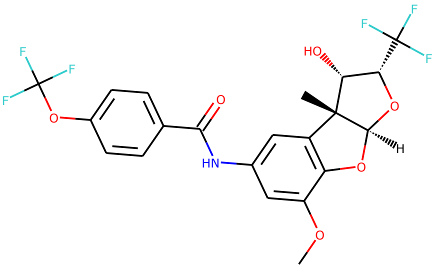

Generated molecule 402

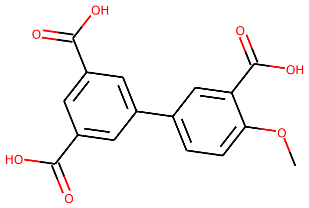

Generated molecule 403

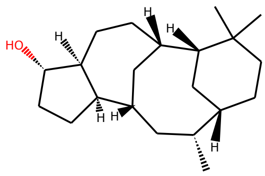

Generated molecule 404

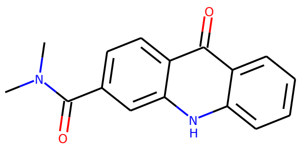

Generated molecule 405

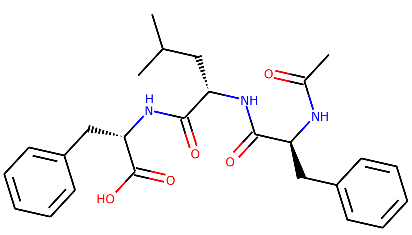

Generated molecule 406

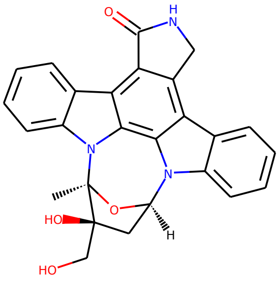

Generated molecule 407

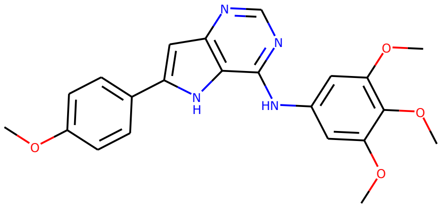

Generated molecule 408

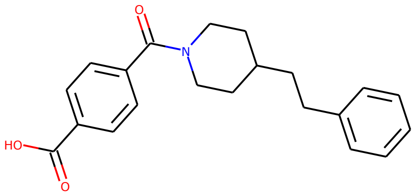

Generated molecule 409

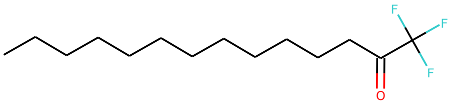

Generated molecule 410

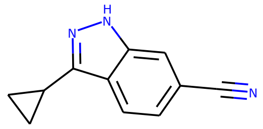

Generated molecule 411

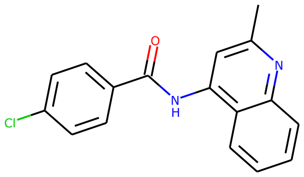

Generated molecule 412

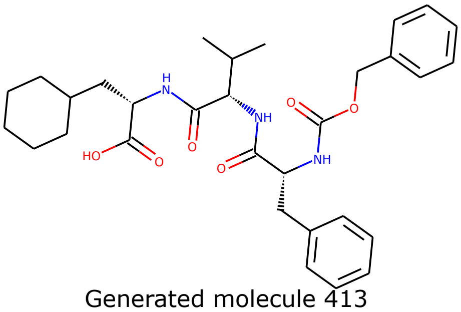

Generated molecule 413

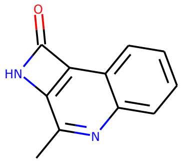

Generated molecule 414

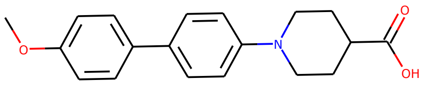

Generated molecule 415

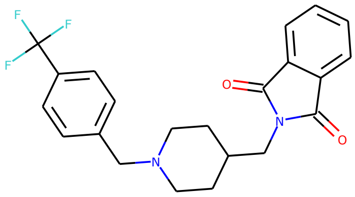

Generated molecule 416

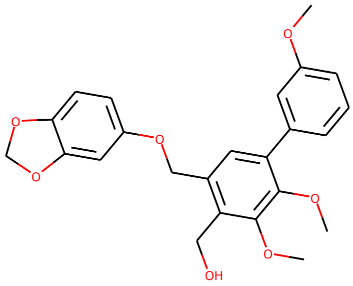

Generated molecule 417

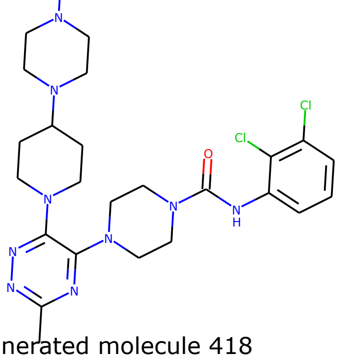

Generated molecule 418

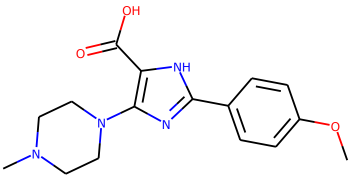

Generated molecule 419

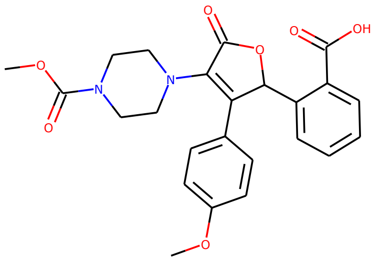

Generated molecule 420

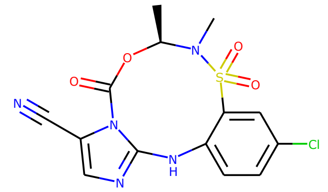
Generated molecule 421

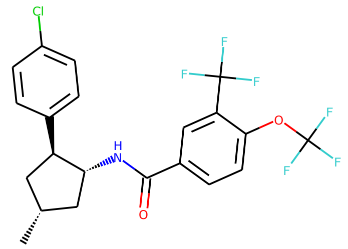
Generated molecule 422

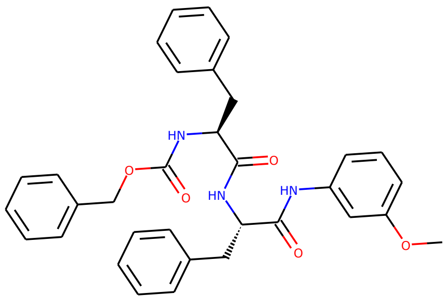
Generated molecule 423

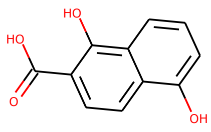
Generated molecule 424

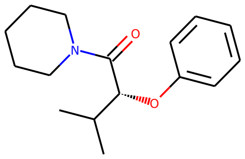
Generated molecule 425

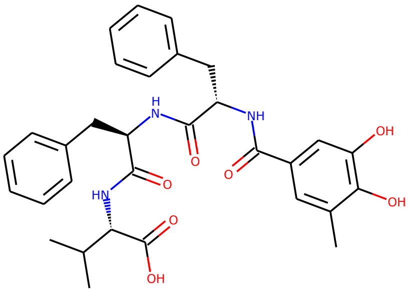
Generated molecule 426

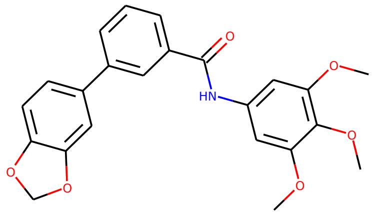
Generated molecule 427

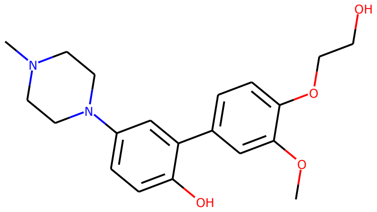
Generated molecule 428

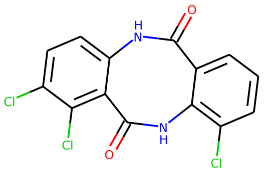
Generated molecule 429

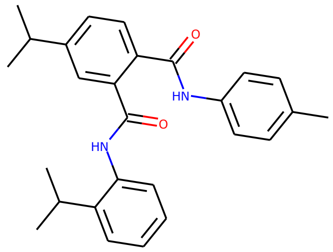
Generated molecule 430

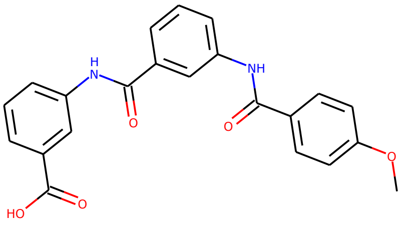
Generated molecule 431

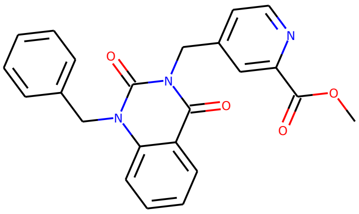
Generated molecule 432

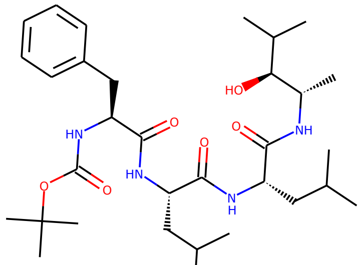
Generated molecule 433

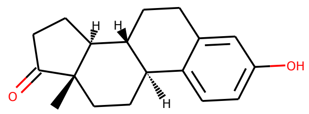
Generated molecule 434

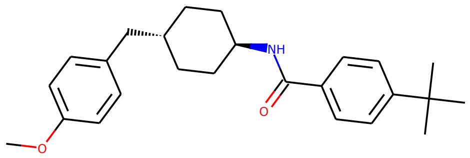
Generated molecule 435

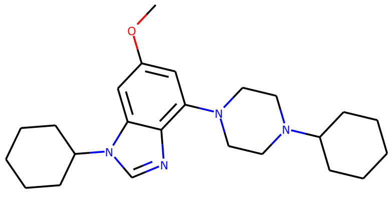

Generated molecule 436

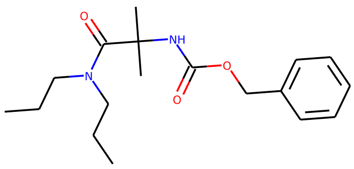

Generated molecule 437

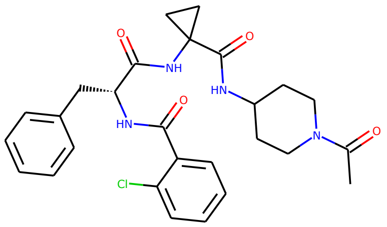

Generated molecule 438

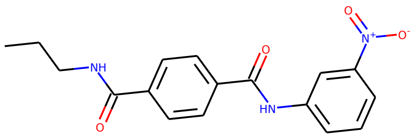

Generated molecule 439

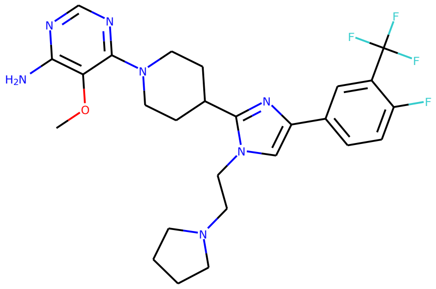

Generated molecule 440

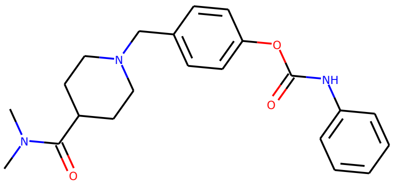

Generated molecule 441

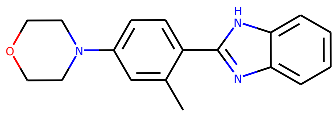

Generated molecule 442

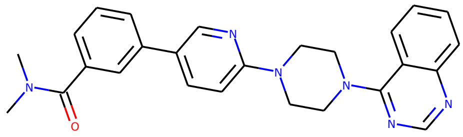

Generated molecule 443

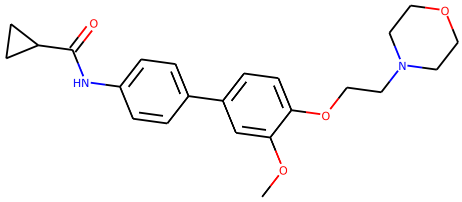

Generated molecule 444

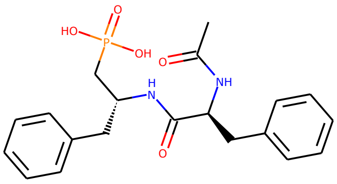

Generated molecule 445

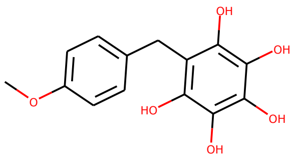

Generated molecule 446

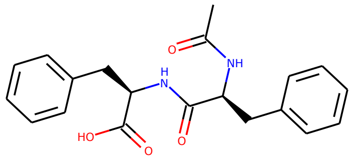

Generated molecule 447

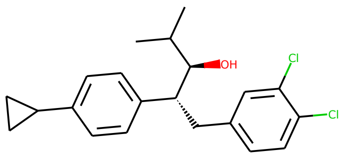

Generated molecule 448

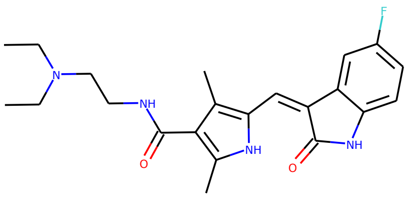

Generated molecule 449

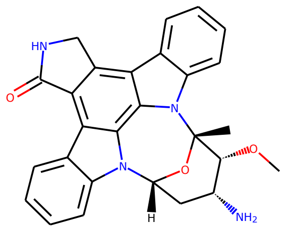

Generated molecule 450

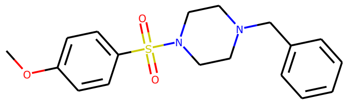

Generated molecule 451

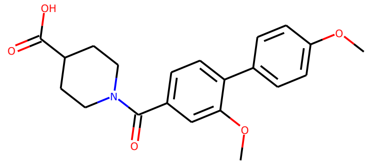

Generated molecule 452

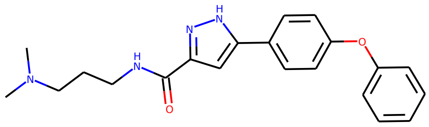

Generated molecule 453

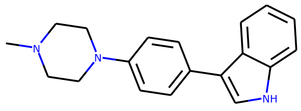

Generated molecule 454

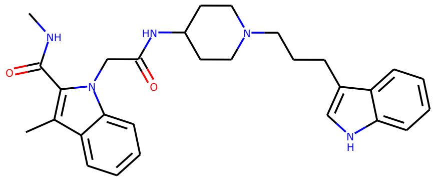

Generated molecule 455

Supplementary Figure S2. Graph representation for generated compounds.

### ROC curves and corresponding AUC for the structures of Insulin-like growth factor 1 receptor (IGF-1R) and Vascular endothelial growth factor receptor 2 (VEGFR2).

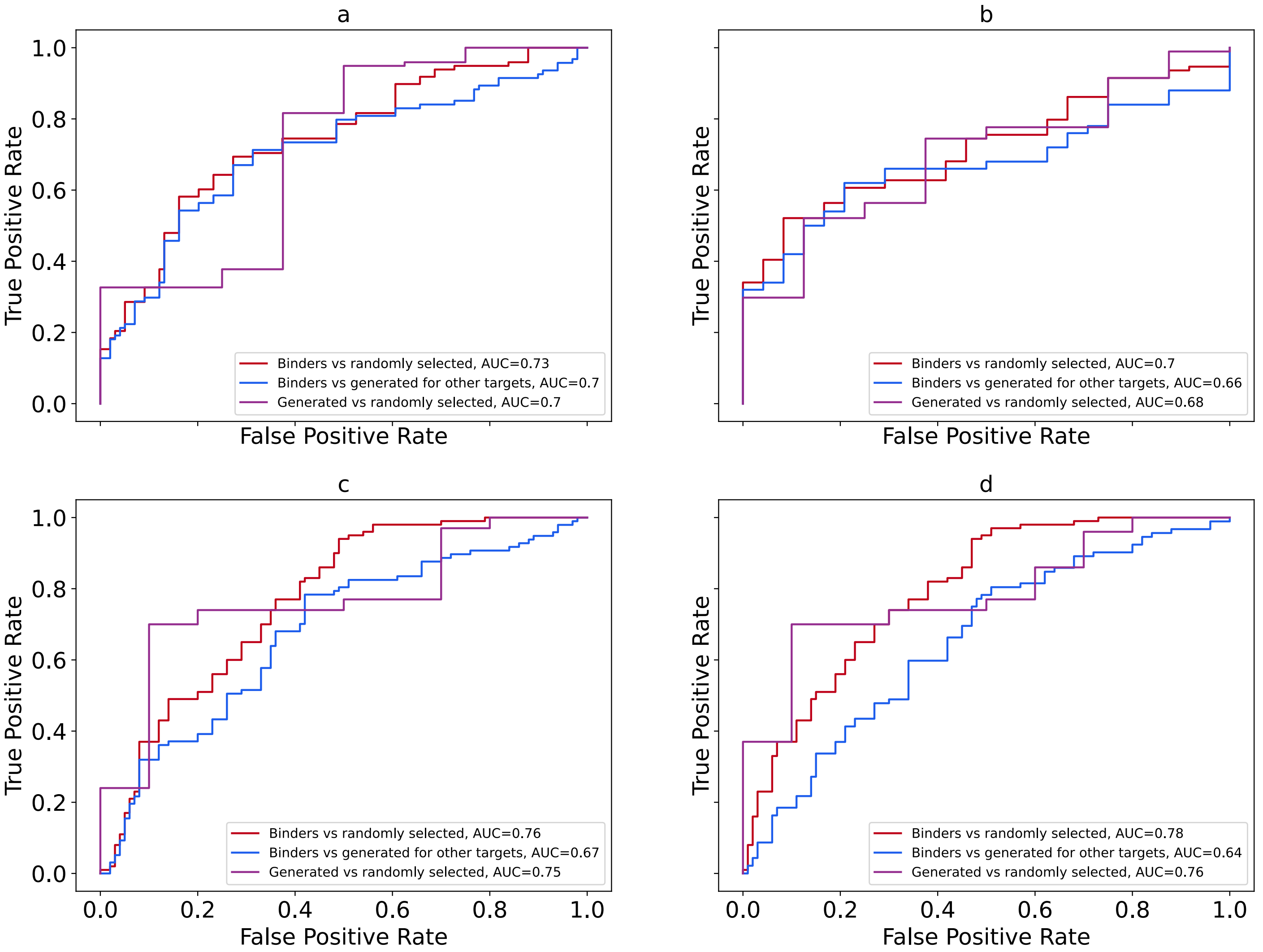


Supplementary Figure S3. ROC curves and corresponding AUC for the following structures:
(a) structure of IGF-1R with PDB code 4D2R
(b) structure of IGF-1R with PDB code 2OJ9
(c) structure of VEGFR2 with PDB code 4ASE
(d) structure of VEGFR2 with PDB code 2P2H

### ROC comparison of known binders versus generated molecules and corresponding AUC for the structures of Insulin-like growth factor 1 receptor (IGF-1R) and Vascular endothelial growth factor receptor 2 (VEGFR2).

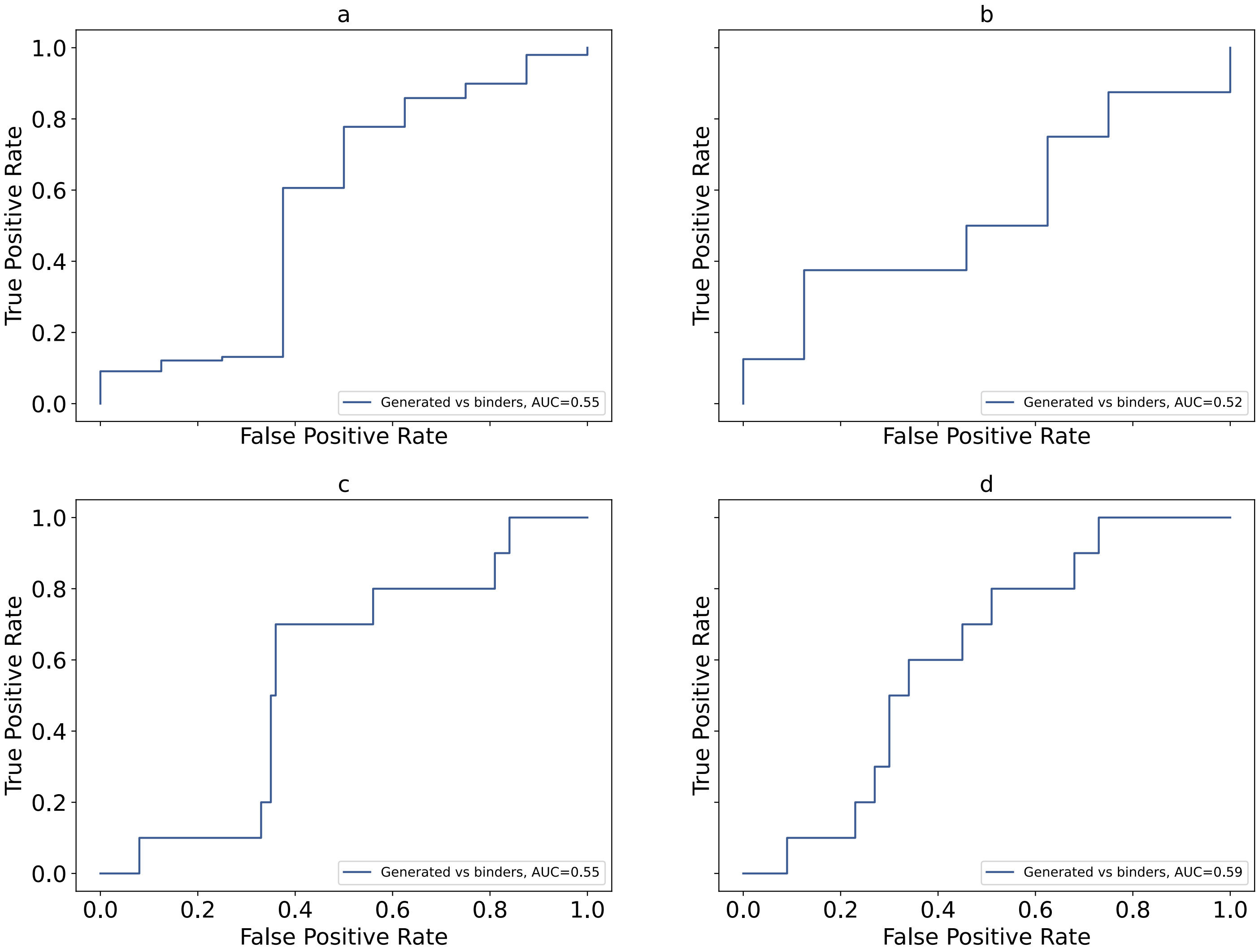


Supplementary Figure S4. ROC comparison of known binders versus molecules generated for IGF-1R and VEGFR2 and corresponding AUC for the following structures:

(a) structure of IGF-1R with PDB code 4D2R
(b) structure of IGF-1R with PDB code 2OJ9
(c) structure of VEGFR2 with PDB code 4ASE
(d) structure of VEGFR2 with PDB code 2P2H
